# Ribosome demand links transcriptional bursts to protein expression noise

**DOI:** 10.1101/2023.10.19.563206

**Authors:** Sampriti Pal, Upasana Ray, Riddhiman Dhar

## Abstract

Stochastic variation in protein expression generates phenotypic heterogeneity in a cell population and has an important role in antibiotic persistence, mutation penetrance, tumor growth and therapy resistance. Studies investigating molecular origins of noise have predominantly focused on the transcription process. However, the noise generated in the transcription process is further modulated by translation. This influences the expression noise at the protein level which eventually determines the extent of phenotypic heterogeneity in a cell population. Studies across different organisms have revealed a positive association between translational efficiency and protein noise. However, the molecular basis of this association has remained unknown. In this work, through stochastic modeling of translation in single mRNA molecules and empirical measurements of protein noise, we show that ribosome demand associated with high translational efficiency in a gene drives the correlation between translational efficiency and protein noise. We also show that this correlation is present only in genes with bursty transcription. Thus, our work reveals the molecular basis of how coding sequence of genes, along with their promoters, can regulate noise. These findings have important implications for investigating protein noise and phenotypic heterogeneity across biological systems.

## Introduction

Genetically identical cells often show variation in gene expression under identical environmental condition. Stochastic variation in gene expression, termed as expression noise, generates phenotypic heterogeneity which has important implications for many biological processes. Gene expression noise has been associated with persistence of microbial cells in antibiotics^1,2^, incomplete mutation penetrance^3–5^, cellular decision-making^6,7^, cancer progression, and anti-cancer therapy resistance^8–10^. Transcription occurring in bursts^11–13^, with a promoter transitioning between on- and off-states, is a major source of expression noise, and each promoter has its characteristic burst frequency and burst size^14–18^. Earlier studies have associated presence of TATA box motif^19,20^, occurrence of TFIID and SAGA co-activator complexes with TATA binding protein ^21^, nucleosome occupancy, and histone modifications with expression noise^22–25^. In addition, transcription factors (TFs) and the gene regulatory network play important roles in determining noise ^26–28^.

The expression noise generated by transcription is modulated by the translation process^13,29^. Thus, a gene that exhibits high expression noise at the mRNA level may not show high noise at the protein level. Two early studies observed a positive association between translational efficiency and expression noise in bacteria and yeast^30,31^, where an increase in mean protein expression, engineered by changes in synonymous codons, led to an increase in expression noise. This contrasted with the traditional inverse relationship between mean protein expression and noise observed in empirical measurements of noise in bacteria and yeast^14–16^. Salari *et al.*^32^, through a computational analysis of expression noise in yeast, observed an increase in positive correlation between tRNA adaptation index (tAI), a proxy for translational efficiency^33,34^, and expression noise for genes with tAI values up to 0.55. The correlation, however, decreased thereafter^32^. In addition, a recent study in *Arabidopsis* reported a positive correlation between translational efficiency and expression noise^35^, similar to what had been observed in microbes. Taken together, these studies showed that the translation process impacted protein expression noise in similar manner across biological systems.

What is the molecular basis of the positive correlation between translational efficiency and protein expression noise? The answer to this question remains unclear, although some mechanisms have been proposed. To achieve a specific protein expression level, a cell can produce many mRNA molecules and translate them at a low rate, or can start with a small number of mRNA molecules and translate them at a high rate^36^. It has been argued that the latter scenario requiring high translation rate may lead to more noise, due to constraints on availability of mRNA, or due to fluctuations in small mRNA numbers^36,37^. However, there has only been indirect evidence in support of this hypothesis^36^. Further, recent studies have shown that the translation process is bursty in nature and can switch between on- and off-states^38,39^, just like the transcription process. Whether translational bursting along with transcriptional bursting can explain the correlation between translational efficiency and noise remains unknown.

In this work, we show that fluctuations in mRNA levels, originating from bursty transcription, combined with high translational efficiency can lead to high protein noise. Through stochastic modeling of translation in single mRNA molecules, we uncovered that the extent of demand for the ribosomal machinery needed for translation at the level of individual genes determined how transcriptional noise was translated into protein noise. In addition, we found that the positive association between translational efficiency and protein noise was visible only for genes that showed bursty transcription. Consequently, the ribosome demand model predicted that a reduction in translational efficiency or a departure from bursty transcription to a more uniform rate of transcription would abolish the association between translational efficiency and protein noise. We validated both these predictions through an experimental noise measurement system in yeast, thereby establishing ribosome demand as the molecular link between transcriptional bursts and protein noise. Taken together, our work reveals the molecular basis of how coding sequence of genes can regulate protein noise via modulating the translation rate. These findings have implications for investigating protein noise and phenotypic heterogeneity across biological systems.

## Results

### High protein noise in genes with low transcription and high translation stems from high mRNA noise

We first tested the existing hypothesis which posits that the genes with low transcription and high translation would exhibit more protein expression noise than the genes with high transcription and low translation (Fig. 1A). Although it has been reported that essential genes in the yeast *Saccharomyces cerevisiae* employ high transcription and low translation strategy for expression, presumably because of detrimental effects of high noise in these genes^36^, there is no direct supportive evidence. To test this hypothesis, we compared protein expression noise of genes showing different levels of transcription and translation in the yeast *S. cerevisiae*. We obtained the mRNA expression data from the study of Nadal-Ribelles *et al.*^40^, who quantified mRNA expression at the level of single cells in yeast (Fig. S1A). We used the data on protein synthesis rate per mRNA from Riba *et al.*^41^ as a measure of translational efficiency (Fig. S1B). We obtained the protein expression noise data from Newman *et al.*^15^, and used their measure of Distance-to-Median (DM), that is corrected for the dependence of expression noise (coefficient of variation, CV = standard deviation of protein expression level/mean protein expression) on mean protein expression, as the measure of protein noise. We obtained protein noise values of 2763 genes from this dataset (Fig. S1C).

**Fig. 1.**
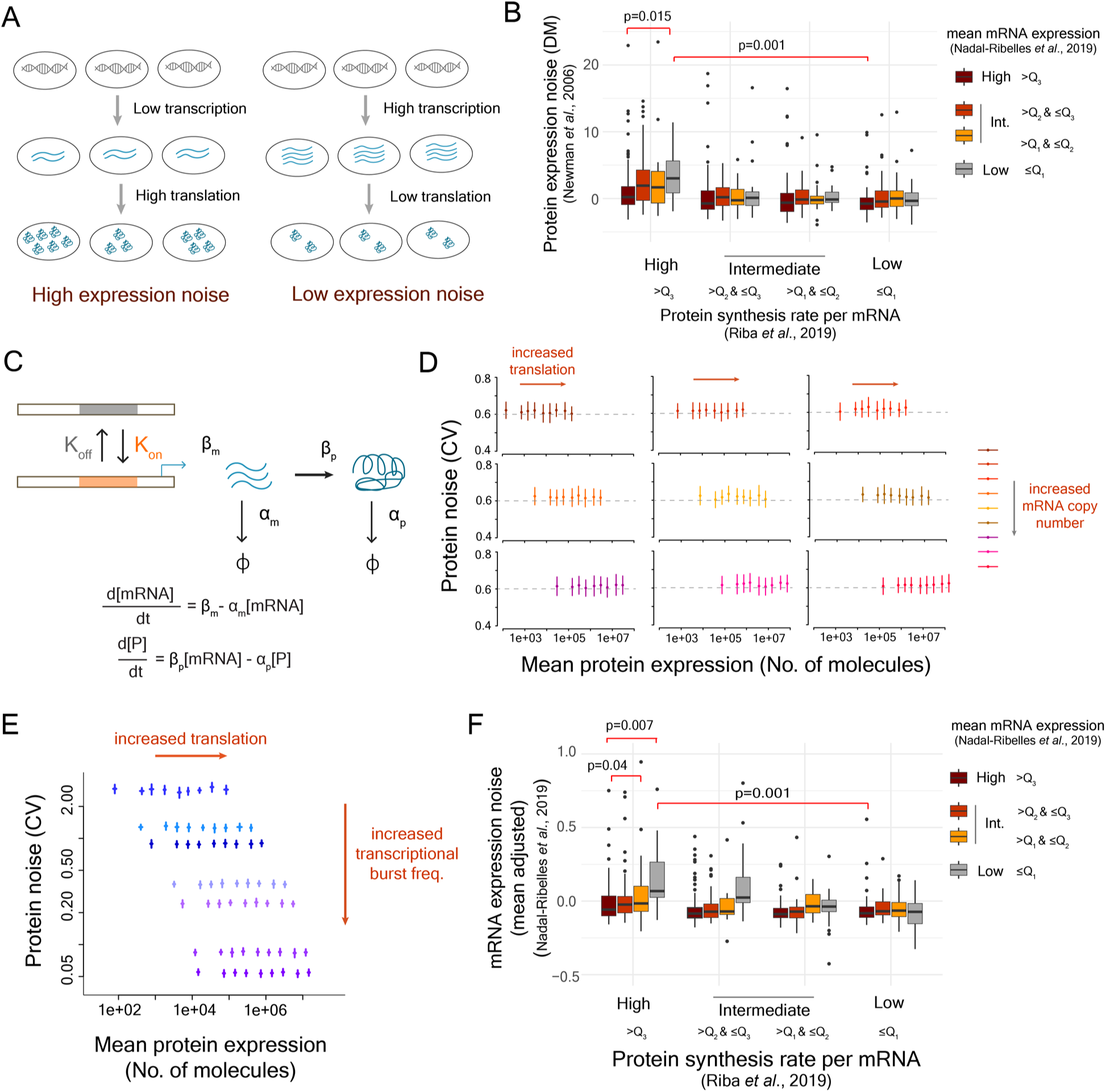
High mRNA noise combined with high translational efficiency leads to high protein noise. **(A)** Existing model describing the impact of translational efficiency on protein expression noise. **(B)** Genes were classified into 16 classes according to the quartiles of mean mRNA expression, calculated from Nadal-Ribelles *et al.*^40^, and then by the quartiles of protein synthesis rates per mRNA from Riba *et al*.^41^. The protein noise values for genes in each of the classes were obtained from Newman et al. (2006), and the measure distance-to-median (DM) value, as derived in their work, was considered as the measure of noise. **(C)** Two-state model of gene expression with the transition rates K_on_ and K_off_ between transcriptional ON and OFF states was used for stochastic simulations **(D-E)** Relationship between mean protein expression and protein noise (CV) obtained from stochastic modeling using the two-state model. The panel (D) describes the results of stochastic simulations obtained at different starting mRNA numbers of a gene to test whether the mRNA expression level of a gene can explain the positive relationship between mean protein expression and protein noise. The panel (E) shows the results obtained from stochastic simulations at different transcriptional burst frequencies, but keeping the starting mRNA number of the gene constant. **(F)** Mean-adjusted mRNA expression noise calculated from the single-cell RNA-seq data^40^ in 16 classes of genes classified according to the quartiles of mean mRNA expression and the quartiles of translational efficiency based on the data on protein synthesis rate per mRNA from Riba *et al*., 2019^41^. Q_1_, Q_2_, and Q_3_ represent first, second and third quartiles.

Mean mRNA expression and translational efficiency individually showed poor correlation with protein noise (Fig. S2). We therefore classified yeast genes based on the quartiles of mean mRNA expression values, and then by the quartiles of translational efficiency resulting in a total of 16 classes (Fig. 1B). The group of genes with low mean mRNA expression and high translational efficiency showed significantly higher protein expression noise (DM, median noise = 3.02) compared to the genes with high mRNA expression and low translational efficiency (median protein noise = -0.724; Mann-Whitney U-test, p=0.001; Fig. 1B), suggesting that low mRNA expression and high translational efficiency was indeed associated with high protein noise.

To better understand how low mRNA expression and high translational efficiency could lead to high protein noise, we employed a two-state model of gene expression^42^. Briefly, a gene can switch between on- and off-states with rates K_on_ and K_off_, respectively (Fig. 1C). In the on-state, the gene is transcribed at a rate β_m_, and the mRNA molecules produced are translated at a rate β_p_. Using stochastic simulations following Gillespie’s algorithm^43^, we modeled the changes in concentrations of mRNA and protein molecules over time individually in 1000 cells, and further calculated mean protein expression and noise values (coefficient of variation, CV) (see Methods). In all simulations, we quantified protein expression noise by coefficient of variation as it was not possible to calculate mean-adjusted noise or DM for a single gene. We modeled the impact of mRNA expression and translational efficiency on expression noise by varying the transcription (β_m_) and the translation (β_p_) rates over a wide range of values (Fig. 1D). However, we did not see any change in protein noise, either with changes in the translation rate for a specific mean mRNA expression, or with changes in the mean mRNA expression (CV= ∼0.62, Fig. 1D). Further, simulations over a wide-range of burst frequencies (K_on_) did not reveal any correlation between translational efficiency and protein noise (Fig. 1E). These results suggested that the differences in the mRNA levels and the translational efficiency could not explain the differences in noise values between low-mRNA high-translation class compared to the high-mRNA low-translation class.

Next, we tested whether higher heterogeneity in mRNA numbers associated with low mean mRNA expression could explain the higher protein noise in the genes of the low-mRNA high-translation class^37^ from the experimental data. To do so, we derived a measure of mean-adjusted mRNA expression noise^28^, that accounts for dependence of mRNA expression noise on mean mRNA expression, for 5475 genes from yeast single-cell RNA-seq data^40^ (Fig. S3A,B). Overall, some of the genes with low mean mRNA expression showed high mean-adjusted mRNA expression noise (Fig. S3C). Next, we compared mRNA expression noise of the 16 classes of genes as obtained based on the levels of mean mRNA expression and translational efficiency (Fig. 1B). The class of genes with low mean mRNA expression and high translational efficiency showed the highest mRNA noise among all classes (Fig. 1F). Specifically, this class of genes had significantly higher mRNA noise (median mRNA noise = 0.068) compared to the class with high mRNA level and low translational efficiency (median mRNA noise = -0.082; Mann-Whitney U-test, p=0.001, Fig. 1F). In general, mean-adjusted protein expression noise showed a moderate correlation with mean-adjusted mRNA noise (Pearson’s correlation = 0.44 and Spearman’s correlation = 0.29, Fig. S3D). These results suggested that high mRNA noise associated with low transcription, rather than the low mean mRNA level itself, combined with translational efficiency was a good determinant of protein noise.

### Bursty transcription combined with high translational efficiency leads to high protein noise

Based on the above observations, we put forward a new working model that highlighted the influence of mRNA noise in determining protein noise. Noise in mRNA expression is usually associated with bursty transcription, where low burst frequency leads to high mRNA expression noise and an increase in burst frequency gradually lowers mRNA noise as the transcription rate becomes more uniform. Therefore, we hypothesized that among genes that exhibited bursty transcription, the ones with high translational efficiency would show higher protein noise compared to the ones with low translational efficiency (Fig. 2A).

**Fig. 2.**
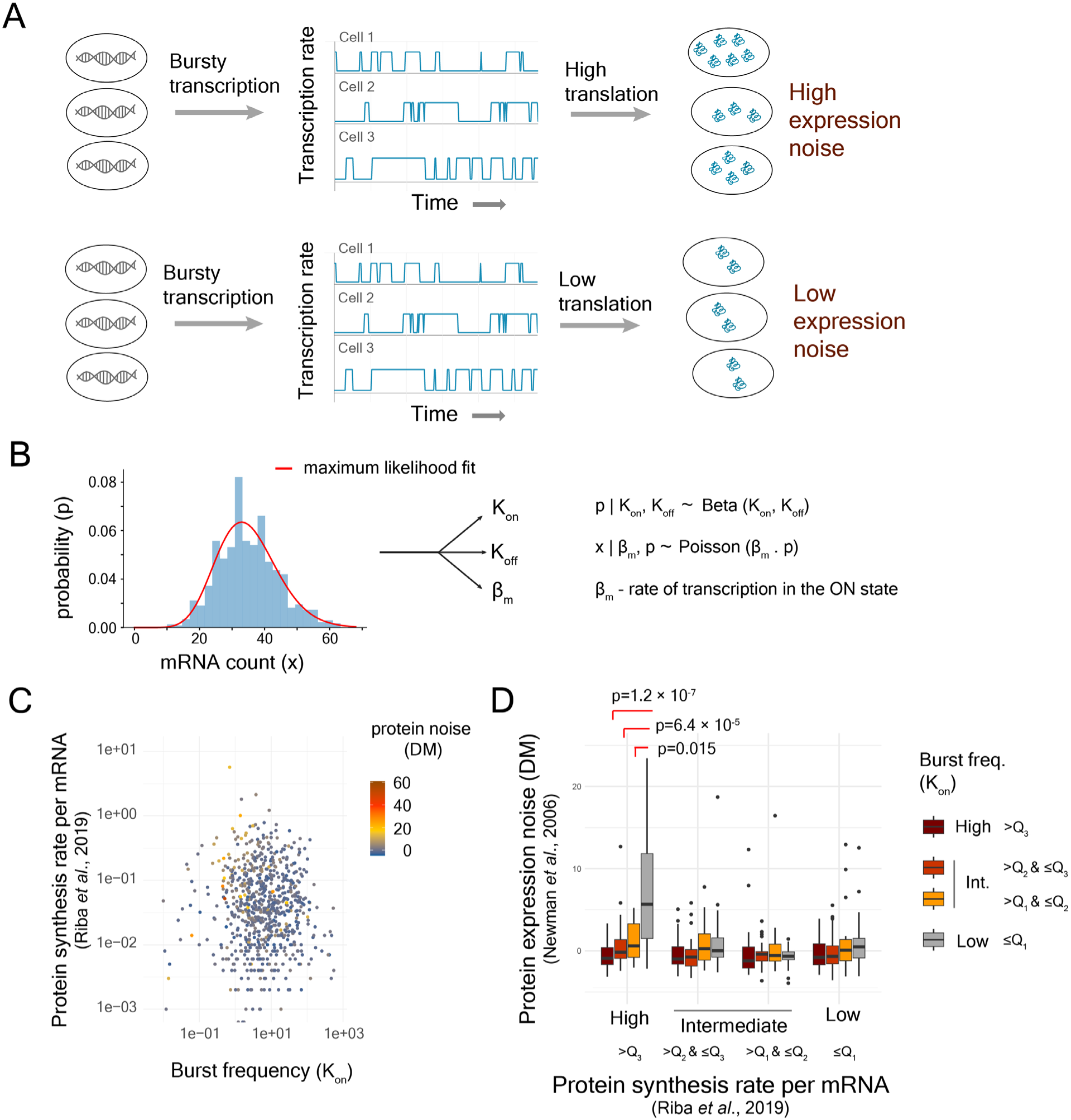
Stochastic fluctuation in mRNA expression, originating from transcriptional bursts, combined with high translational efficiency generates high protein noise. **(A)** The new working model postulated that the genes with bursty transcription (low transcriptional burst frequency) and high translational efficiency were likely to exhibit higher protein expression noise compared to the genes with bursty transcription but low translational efficiency. **(B)** Estimation of parameters of two-state model of gene expression from single-cell RNA-seq data as described by Kim and Marioni (2013)^44^. **(C)** Protein noise of genes with different levels of transcriptional burst frequencies and translational efficiency, estimated by protein synthesis rates per mRNA^40^. **(D)** Protein expression noise (DM values from Newman *et al*., 2006) of genes classified into 16 classes based on burst frequency and translational efficiency (protein synthesis rate per mRNA^41^). Q_1_, Q_2_, and Q_3_ represent first, second and third quartiles, respectively.

To test this new model, we estimated the parameters associated with transcriptional bursts such as on- and off-transition rates (K_on_ and K_off_) and transcription rate (β_m_) for >6000 genes in yeast from single-cell RNA-seq data^40^ (Fig. S4), using a maximum-likelihood approach based on a Poisson-Beta model^44^ (Fig. 2B). Genes with low transcriptional burst frequencies showed high mRNA noise (Fig. S5). However, the genes that exhibited high protein noise were associated with high translational efficiency (Fig. 2C). In addition, we classified genes into 16 classes according to the quartiles of burst frequency and the quartiles of translational efficiency^40^ (Fig. 2D). Genes with low burst frequency and high translational efficiency showed the highest protein noise among all 16 classes (Fig. 2D). This class of genes showed a mean protein noise (DM) value of 5.65, whereas, the class of genes with high burst frequency and high translational efficiency showed a mean protein noise (DM) value of -0.038 (Mann-Whitney U test, p=1.22 × 10^-7^; Fig. 2D). These observations confirmed that high translational efficiency, combined with bursty transcription, could lead to high protein noise (Fig. 2D). Analysis with tAI as the measure of translational efficiency showed similar trend (Fig. S6, S7), suggesting that these results were robust to the use of different measures of translational efficiency.

### Bursty translation does not explain the correlation between translational efficiency and protein noise

Even though results from the analysis of earlier experimental data agreed with the new working model, they did not explain how high translational efficiency combined with low burst frequency could lead to high protein noise, and how reducing translational efficiency could lower protein noise. Recent studies have observed that translation, like transcription, also occurs in bursts^38,39^. Specifically, Livingstone *et al*. ^39^, through single-cell mRNA tracking, measured the parameters of translational bursting and identified the roles of 5’UTR and 5’mRNA cap in modulation of translational burst frequency and burst amplitude.

Therefore, we asked whether incorporating translational bursting into our model could reveal the positive correlation between translational efficiency and protein noise. To do so, we built a TASEP based model^45–46^ of translation, with appropriate modifications, at the level of single mRNA molecules (Fig. 3A, see Methods). We assumed that an mRNA molecule could transition between on- and off-states^39^ with the rates TL_on_ and TL_off_, respectively, and in the on-state, translation initiation occurred at the rate of TL_init_ (Fig. 3A). As multiple ribosomes could translate a single mRNA molecule at the same time, a second translation initiation happened only when the preceding ribosome traversed at least 10 codons, to account for steric interaction between ribosomes^47,48^ (Fig. 3B). Several earlier studies have showed the presence of a gradually increasing profile of translational efficiency or ramp, although of different degree, in the 5’ end of coding regions of genes^34,49^. This meant that the translation speed was lower near the 5’ end of a gene and the speed gradually increased as a ribosome traversed from 5’ to 3’ end of an mRNA molecule. Taking this into account, we modeled the traversal speed of the ribosomes using a first-order Hill function where the traversal speed (V) was dependent on the position of the ribosome in the coding sequence (Eq. 10; Fig. 3B). An increase in the value of the parameter K_Hill_ led to an increased traversal time (Fig. 3C). The speed of traversal has an impact on translation initiation rate, as observed by Barrington *et al.*^50^. This meant that K_Hill_ was also related to the translation initiation rate, TL_init_. Higher translation initiation rate necessitated lower K_Hill_ to ensure faster traversal of the preceding ribosome through first 10 codons to avoid collision between two successive ribosomes (Fig. 3D). Similarly, lower K_Hill_ value led to faster traversal of ribosomes through an mRNA molecule, and thereby, permitted a greater number of translation initiation events per unit time (Fig. 3D). Thus, the translation initiation rate and the ribosome traversal time through an mRNA molecule were interconnected (Fig. 3D).

**Fig. 3.**
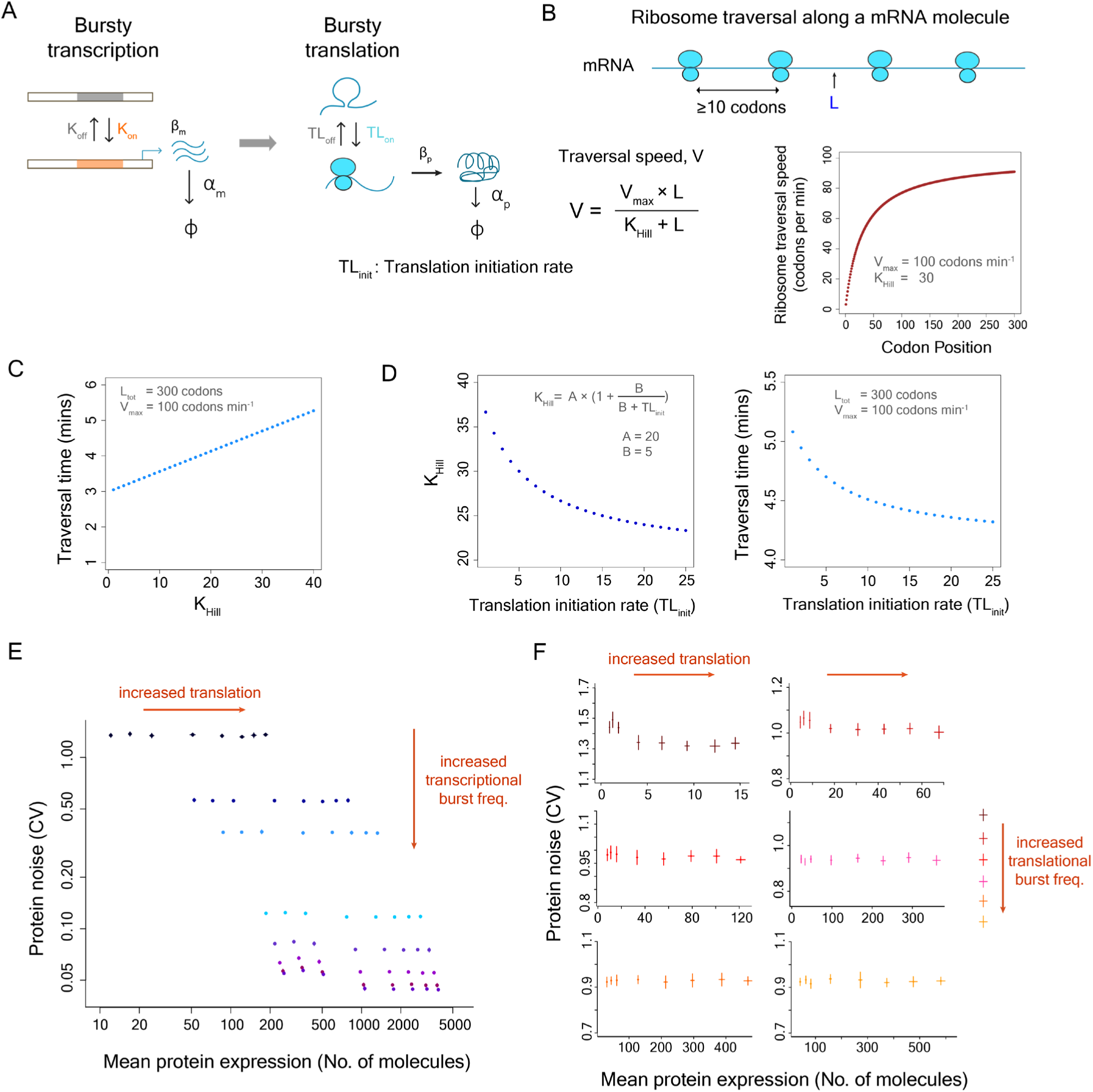
The model combining transcriptional and translational bursting does not explain the positive correlation between translational efficiency and protein noise. **(A)** Schematic diagram depicting the integrated model of transcriptional and translational bursting. **(B)** Ribosomal traversal speed along an mRNA molecule is given by V at a position L in the mRNA molecule. As multiple ribosomes could translate a single mRNA molecule at the same time, a second translation initiation happened only when the preceding ribosome traversed at least 10 codons, to account for steric interaction between ribosomes^47,48^. The ribosome traversal speed was modeled using a Hill function as several studies have shown the presence of a gradually increasing profile of translational efficiency or ramp in the 5’ end of coding regions of genes^34,49^ **(C)** Traversal time calculated as a function of K_Hill_ from the Hill function for a gene with 300 codons, and the maximum traversal speed of 100 codons per minute. **(D)** Relationship between K_Hill_, translation initiation rate and ribosome traversal time. Faster ribosome traversal enabled higher translation initiation rate (TL_init_)^50^. A and B are parameters of the model. **(E)** The results obtained from stochastic simulations using the combined model of transcriptional and translational bursting. Protein noise changes with changes in translational efficiency and transcriptional burst frequency, but do not reveal a positive correlation between mean protein expression and protein noise **(F)** Protein noise obtained from the combined model changes with changes in translation initiation rate and translational burst frequency, but do not explain the positive correlation between mean protein expression and protein noise.

We integrated the model of translation with the two-state model of transcription, and performed stochastic simulations in individually in 1000 cells to estimate mean protein expression and protein noise (coefficient of variation, CV). To test whether incorporation of the translational bursting could capture the positive correlation between mean protein expression and protein noise, we changed the translational efficiency by varying the translation initiation rate (TL_init_) in our simulations, thereby, also affecting the ribosome traversal time, while keeping all other parameters constant. For all transcriptional burst frequencies examined, we did not observe any change in protein noise with an increase in translational efficiency (Fig. 3E). We also tested the model at different translational burst frequencies. Although changes in translational burst frequencies altered the level of protein noise, we did not find any correlation between mean protein expression and protein noise (Fig. 3F). These results suggested that a simple model integrating transcriptional and translational bursting could not explain the observed relationship between translational efficiency and protein noise.

### Inclusion of ribosome demand reveals a positive correlation between translational efficiency and protein noise

Next, we analyzed several molecular models that could possibly explain the positive correlation between translational efficiency and protein noise. An increase in translational efficiency, leading to a higher translation rate, will require an increased supply of tRNA molecules and ribosomes. Could an increase in translation rate lead to a bottleneck in supply of tRNA, and generate variation in protein expression across cells? This seemed highly unlikely, as high translational efficiency is associated with presence of preferred codons, for which abundant tRNA molecules are available in a cell^33,34^. In comparison, rare codons have lower numbers of corresponding tRNA molecules available^33,34^. Thus, presence of rare codons would, in fact, cause a bottleneck in tRNA supply and hence, would lead to higher expression noise. This would further imply that a reduction in translational efficiency would generate higher expression noise, which would contradict the actual empirical observations^30,31^.

The translational efficiency is dependent on availability of ribosome machinery in the cell. We analyzed whether higher translational efficiency could lead to a bottleneck in the availability of ribosomes for mRNA molecules of a gene, cause variation in the translation process of a gene among cells and thereby, generate high expression noise. It has been reported that competition for ribosomes rather than tRNA, imposes constraints on translation in yeast^51^. In addition, studies in cell-free systems have shown that constraints on resources for transcription and translation can cause bursty expression^52^.

For a gene that is transcribed in bursts, there will be considerable temporal fluctuation in the number of mRNA molecules available for translation. When the mRNA copy number of that gene is high, they will require many ribosomes for translation. As the mRNA copy number drops due to stochastic fluctuations in transcription rate, some of the ribosomes earlier involved in translation of mRNA molecules of this gene become available, and can be allocated for translating mRNA molecules of other genes (Fig. 4A). At a subsequent time-point when the mRNA number of this gene rises again, it increases the demand for ribosomes (Fig. 4A). In actively growing cells, where the demand for ribosomal machinery is high, this can constrain translation initiation rate for these mRNA molecules. As the allocation of ribosomes to mRNA molecules is likely to be a stochastic process, this can generate variation in translation rates and consequently, protein expression of that gene between cells.

**Fig. 4.**
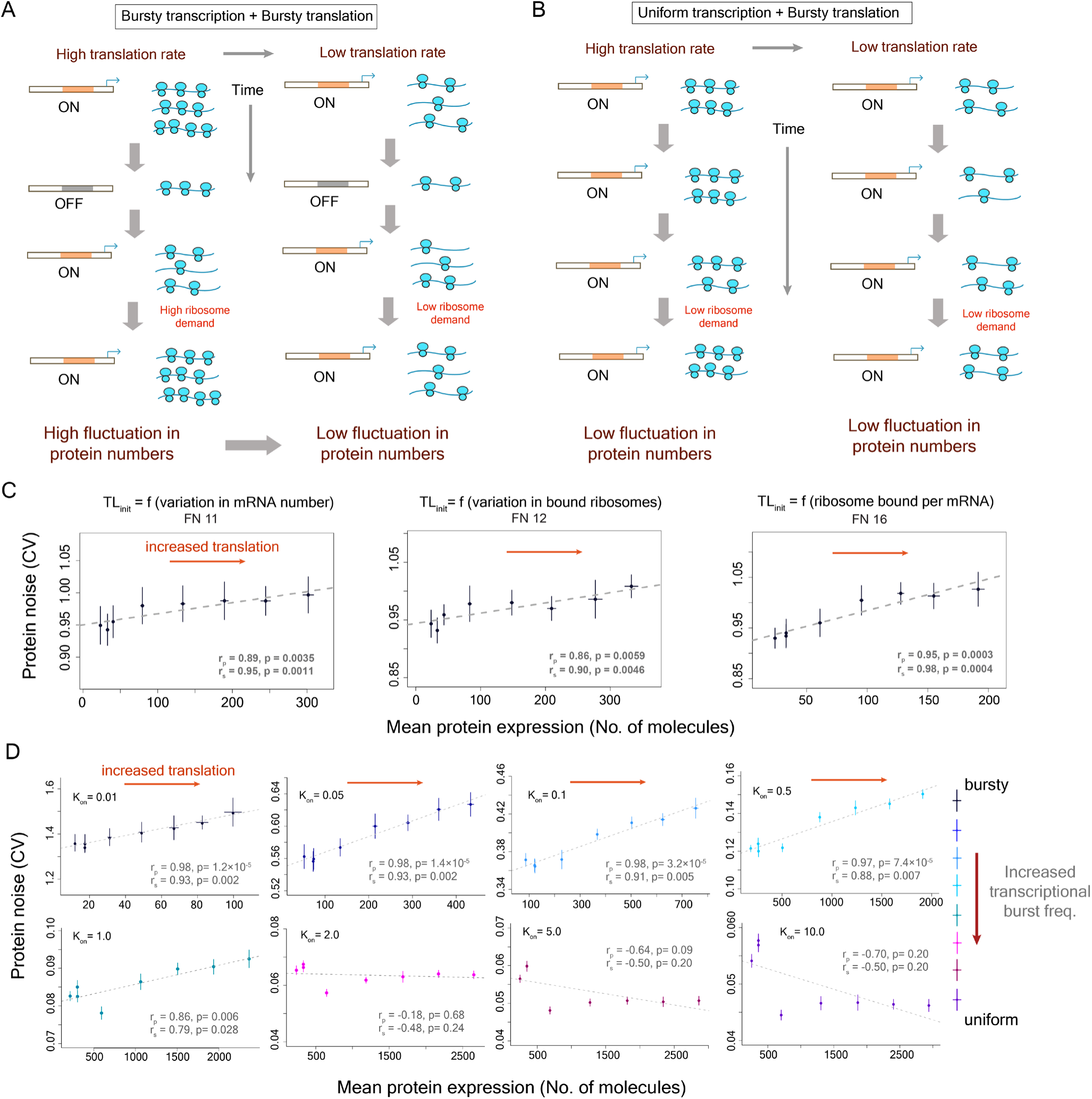
Inclusion of ribosome demand associated with translation of mRNA molecules of a gene can reveal positive correlation between translational efficiency and protein noise. **(A)** Schematic diagram depicting how ribosome demand for translation of mRNA molecules can vary with bursty transcription and bursty translation. As mRNA numbers of the gene fluctuate due to bursty transcription, high translational efficiency can lead to intermittent elevated ribosome demand for translation of the mRNA molecules of that gene. This can lead to increased inter-individual variation in protein numbers **(B)** Uniform transcription of a gene does not lead to a sudden elevated ribosome demand for translation, thereby reducing inter-individual variation in protein numbers **(C)** Results from simulations with three different functions (function 11, 12 and 16 from Table S1) that model the impact of ribosome demand on translational efficiency. Results from simulations with other functions to model ribosome demand are shown in Fig. S8. **(D)** The relationship between mean protein expression and protein noise at different transcriptional burst frequencies obtained from stochastic simulations with the model incorporating ribosome demand along with transcriptional and translational bursting. The ribosome demand was modeled using function 16 (Table S1). For each transcriptional burst frequency, the translational efficiency was altered by changing the translation initiation rate (TL_init_) while keeping rest of the parameters constant.

There are two predictions that come out of this model. First, it predicts that a reduction in the high demand for ribosome machinery during the low- to high-mRNA copy-number transition will lower protein expression noise (Fig. 4A). This can be achieved by lowering the translational efficiency of the gene. In this scenario, we will observe high protein noise at high translational efficiency and low noise at low translational efficiency, which will agree with the empirical observations.

Another way of minimizing the sudden demand for ribosomes is to reduce temporal fluctuations in mRNA numbers. This can be achieved at the level of transcription by employing a more uniform rate of transcription (Fig. 4B). Thus, according to the second prediction, protein noise of genes that are expressed in a uniform rate of transcription will be minimally affected by changes in the translational efficiency (Fig. 4B).

In addition, we explored whether co-translational protein folding, ribosome collision, translational errors, or changes in mRNA stability associated with translation rate could explain the positive association between translational efficiency and protein noise. Proteins are co-translationally folded as they exit through the ribosome tunnel after translation^53^. The speed of translation might affect the efficiency of co-translational folding, and perhaps, fast translation could lower efficiency of co-translational folding. This can generate misfolded proteins that are degraded. Since this process is stochastic in nature, it can lead to increased variation in protein expression among cells within a population. However, several studies have reported the exactly opposite phenomenon. A study by Spencer *et al.*^54^ showed that the use of optimal codons leads to better folding of the encoded polypeptide. Other studies have shown that fast translation can in fact help avoid misfolded states, and can help in better co-translational folding, especially for misfolding-prone regions^55,56^.

High translational efficiency means fast traversal of ribosomes through an mRNA molecule which would in turn increase the translation initiation rate. This can lead to a higher chance of collision between two successive ribosomes on an mRNA molecule in case of a slow-down in movement due to occurrence of non-preferred codons. Slower ribosome traversal reduces translation initiation rate and thus, lowers the chance of ribosome collision even in case of a slow-down. Ribosome collision leads to degradation of mRNA, rescue of bound ribosome and can also affect further translation initiation^57–59^. Therefore, this could be a possible reason for higher protein noise in genes that show high translational efficiency. However, in contrast to the ribosome demand model, this observation would hold for all genes with high translational efficiency, irrespective of their burst frequencies.

We further considered whether translational errors could explain the correlation between translation rate and protein expression noise. It is possible that fast translation can lead to more translational errors. This can generate proteins with wrong amino acids, which can result in protein misfolding and therefore, can lead to protein degradation. Since translational errors occur at random, this process will vary from one cell to another, which can lead to higher protein expression noise. However, there are conflicting reports regarding this. An earlier study reported that translational errors occurred at sites where the speed of ribosome was higher^60^. On the other hand, a recent study observed that preferred or optimal synonymous codons, where ribosome traversal is fast, were translated more accurately^61^.

Finally, we also considered whether the translation process itself could change mRNA stability and therefore, could generate cell-to-cell variation. It has been reported that increased translation can lead to higher mRNA destabilization^62^. In addition, codon optimality and thus elongation rate has also been suggested to be positively associated with mRNA stability^63^. A more recent study has however showed that translation initiation rate, but not elongation rate, is the major determinant of mRNA stability and inhibition of translation initiation has been shown to increase mRNA decay^64^. However, if changes in the translation rates altered mRNA stability and led to increased variation in cell-to-cell variation in protein concentration, this will be observed for all genes irrespective of their burst characteristics.

Since the ribosome demand model agreed with the earlier empirical observations, we tested whether incorporating ribosome demand in our mathematical model could capture the observed positive correlation between translational efficiency and protein noise. The ribosome demand for translating the mRNA molecules of a specific gene was modeled as a resource allocation problem that depended on the fluctuation in the number of mRNA molecules of that gene and the number of ribosomes that were available for translation of these mRNA molecules (Fig. 4A). Large variation in mRNA numbers led to large fluctuations in ribosome demand which also affected translation initiation rate. For example, a sudden surge in the number of mRNA molecules of the gene from one time point to another necessitated more ribosomes for translation, thereby, creating increased ribosome demand for translation of that gene. Similarly, large variation in the number of ribosomes required to be bound to mRNA molecules of that gene between two consecutive time points affected ribosome allocation and influenced translation initiation rate.

Therefore, we modeled ribosome demand using various mathematical functions (Table S1) where we considered it to be a function of variation in mRNA number, variation in ribosome bound per mRNA molecule or a combination of both (the parameter ‘av’, Table S1). The translation initiation rate (TL_init_) at a specific time point varied from the basal translation initiation rate according to Hill functions incorporating ribosome demand (Table S1). The Hill functions were of the order 10 to model a sharp decrease in translation initiation rate if the ribosome demand was too high. For each of the functions for modeling ribosome demand, we performed stochastic simulations and changed mean expression by varying the basal translation initiation rate while keeping other parameters constant (Fig. 4C, Fig. S8).

Interestingly, the functions where translation initiation rate was dependent on the number of ribosomes bound per mRNA molecule showed the most positive association between mean protein expression and protein noise (Fig. 4C, Fig. S8). We, therefore, considered one of these functions (function 16) for all subsequent analysis (Fig. 4C, Table S1). However, we observed similar positive association for several variants of these functions suggesting that the association found was robust to variations in the actual form of the function (Table S1, Fig. S8).

To further test whether our mathematical model could capture the two predictions of the ribosome demand model, we performed stochastic simulations for a wide range of transcriptional and translational burst frequencies (Fig. 4D). At low transcriptional burst frequencies, protein noise increased with an increase in translation rate (Fig. 4D). In addition, as we increased the transcriptional burst frequency and moved towards a more uniform rate of transcription, the translation rate had minimal effect on protein noise (Fig. 4D). Thus, the mathematical model could recapitulate both predictions of the ribosome demand model.

We also tested the impact of translational burst frequency on the association between translational efficiency and protein noise (Fig. 5A). At very low translational burst frequencies, protein noise did not show much change with an increase in translation rate (Fig. 5A). This was due to very low level of protein production at very low translational burst frequency, as mRNA molecule mostly resided in the inactive state. At higher translational burst frequencies, we again observed an increase in protein noise with an increase in the translation rate and hence with an increase in mean protein expression (Fig. 5A). We further tested the robustness of our stochastic modeling approach through a systematic variation in values of several model parameters (Fig. S9-S14), as well as explored different scenarios through random sampling of the parameter space (Fig. S15).

**Figure 5.**
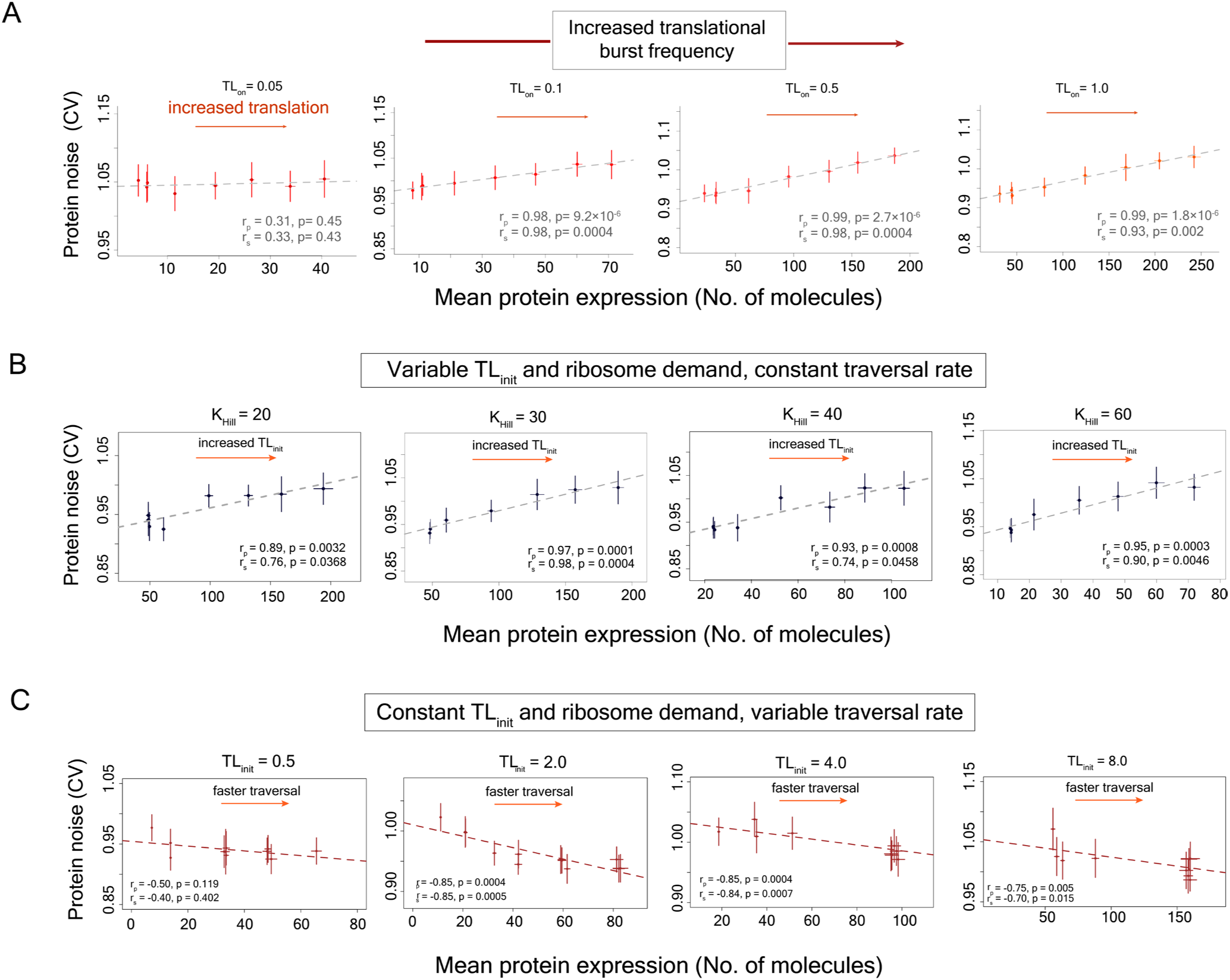
Ribosome demand is necessary for the positive correlation between mean protein expression and protein noise. **(A)** The relationship between mean protein expression and protein noise at different translational burst frequencies obtained from stochastic simulations with the model incorporating ribosome demand along with transcriptional and translational bursting. The ribosome demand was modeled using function 16 (Table S1). For each transcriptional burst frequency, the translational efficiency was altered by changing the translation initiation rate (TL_init_) while rest of the parameters were kept constant. **(B)** Stochastic simulations where mean protein expression was altered by changing base translation initiation rate (TL_init_), thus altering ribosome demand, but keeping ribosome traversal speed constant, maintained positive correlation between mean protein expression and noise. This was done by keeping K_Hill_ constant at specific values during simulations that constrained the traversal time (Eq. 10). **(C)** Stochastic simulations where mean protein expression was altered by changing the ribosome traversal speed (Eq. 10) but keeping the base translation initiation rate (TL_init_) constant, and thus, not allowing variation in ribosome demand with changes in ribosome traversal rate, abolished the positive correlation between mean protein expression and protein noise.

Next, we investigated whether elimination of the ribosome demand from our model could lead to a loss of the positive correlation between mean protein expression and protein noise. To do so, we tested two sets of models where we decoupled the translation initiation rate (TL_init_) from the ribosome traversal speed (V) and changed each of these parameters independently without varying the other. In the first set of models and stochastic simulations, we altered the mean protein expression of the gene by altering the base translation initiation rate but keeping the ribosome traversal speed constant. In these models, ribosome demand varied with changes in the basal translation initiation rate and maintained the positive correlation between mean protein expression and protein noise (Fig. 5B). The results were similar for ribosome traversal speed set at different values defined by the Hill coefficient (K_Hill_) according to the equation 10. In the second set of models and simulations, we changed the mean protein expression by altering the ribosome traversal speed but keeping the translation initiation rate constant. These models did not allow ribosome demand to vary according to the changes in the ribosome traversal speed and showed a loss of positive correlation between mean protein expression and protein noise (Fig. 5C and S16). Thus, these results suggested that the ribosome demand was necessary for maintaining the positive correlation between mean protein expression and protein noise.

### Impact of coding sequence mutations on protein noise is dependent on the burst characteristics of the promoter

To empirically test the predictions of the ribosome demand model, we set up an experimental assay to measure protein expression noise in the yeast model system (Fig. 6A). We built gene constructs where the expression of a green fluorescent protein (*GFP*) gene could be controlled by different promoters with different burst frequencies (Fig. S17). We then introduced several mutant variants of the GFP gene with altered translational efficiencies, under the regulation of these promoters. We integrated all constructs into the yeast genome, to eliminate expression noise arising from fluctuations in plasmid copy number (Fig. 6A). We measured GFP expression using flow cytometry (Fig. S18), and estimated noise from a homogeneous group of cells that showed very similar cell size and cell complexity^14^ (Fig.6B, Fig. S19).

**Fig. 6.**
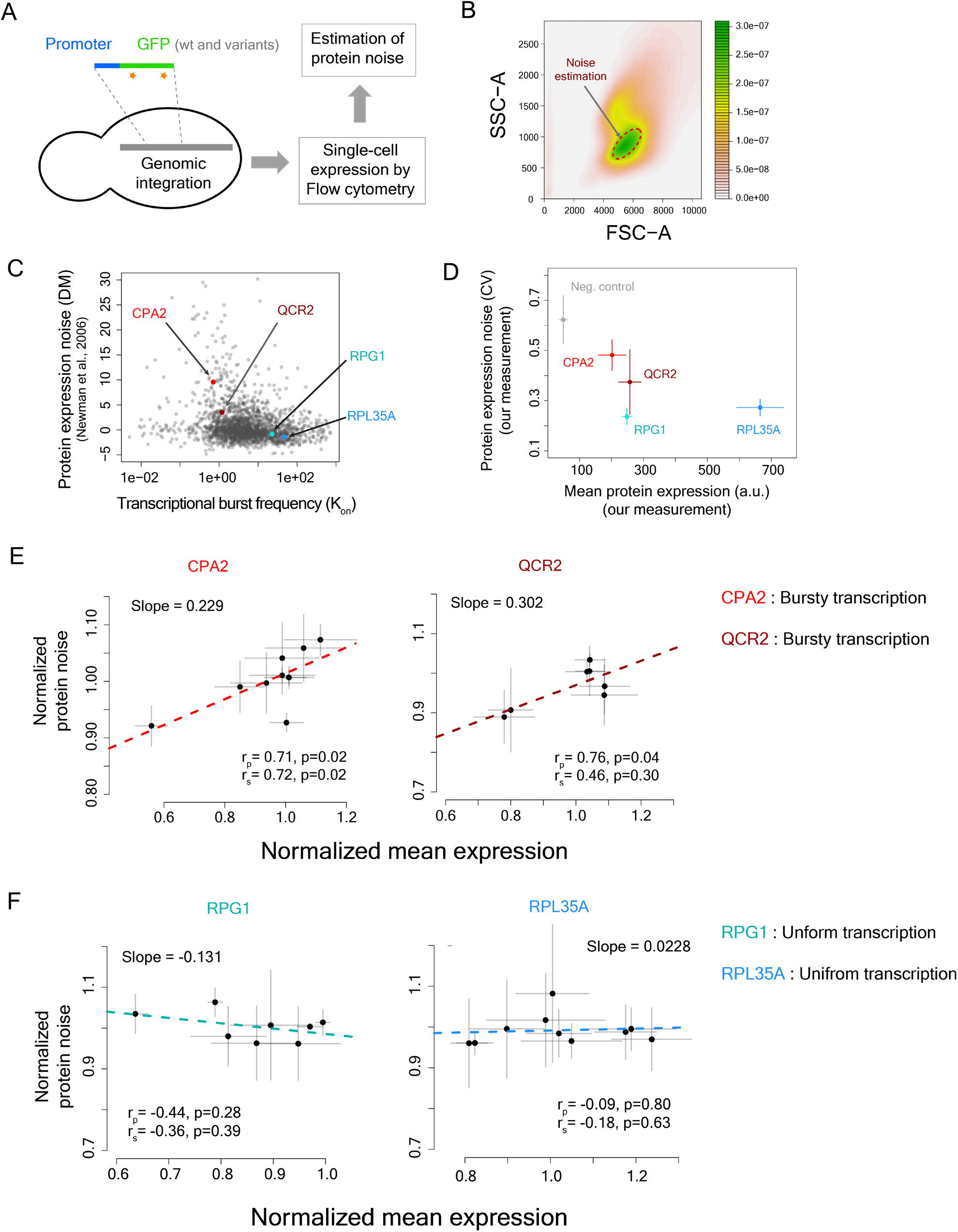
Impact of changes in translational efficiency on protein noise is dependent on the transcriptional burst characteristics of promoters. **(A)** Gene-promoter construct for genomic integration and noise measurement **(B)** Noise estimation from a homogeneous group of cells with similar cell size and complexity **(C)** Protein noise (DM)^15^ vs burst frequency values for yeast genes. Burst frequency values were estimated from single-cell RNA-seq data^40^ using the method described by Kim and Marioni (2013)^44^. **(D)** Measured mean protein expression and protein noise of the promoters of *RPL35A*, *RPG1*, *CPA2* and *QCR2* with the wild-type GFP gene. **(E)** Relationship between normalized protein noise and normalized mean expression of the GFP mutants in the bursty promoters *CPA2* and *QCR2*. (F) Relationship between normalized protein noise and normalized mean expression of the GFP mutants in the non-bursty promoters *RPG1* and *RPL35A*.

We picked four promoters for our constructs (Fig. 6C) - two promoters, of *RPG1* and *RPL35A* genes, with high burst frequencies (K_on_=22.855 and K_on_=47.464 respectively) and low protein noise (DM=-0.841 and DM=-1.4 respectively)^15^, and two promoters, of *CPA2* and *QCR2* genes, with low burst frequencies (K_on_=0.704 and K_on_=1.186 respectively) and high protein noise (DM=9.583 and DM=3.515 respectively)^15^. Our measurements also confirmed that the promoters *CPA2* and *QCR2* had higher protein noise compared to *RPG1* and *RPL35A* (Fig. 6D).

To change translational efficiency in all gene constructs in a similar manner, we built variants of the GFP gene that differed in their codon usage. This altered the translation elongation rate which in turn would also alter the translation initiation rate^50^. We constructed nine mutant variants of the *GFP* gene and cloned each of them under the control of these four promoters. The variants consisted of seven single mutants and two variants with multiple mutations (see Methods). Many of the codon substitutions led to usage of nonpreferred codons in yeast. We specifically focused on mutating N-terminal and C-terminal regions of the GFP protein so as not to affect its functional core. In addition, variants contained mostly synonymous mutations with two exceptions. We changed glutamic acids with aspartic acids at codon 5 and codon 234, and tyrosine with asparagine at codon 237 of the GFP gene. We created all constructs in three replicates, and estimated their noise through multiple independent experiments on different days.

The multi-mutant variant at the N-terminal end, containing five mutations (N-5), had the most number of rare codons and thus, was expected to show substantial reduction in mean protein expression owing to its low translational efficiency. We compared the impact of the coding region mutations across different promoters through the measures of normalized mean expression and normalized protein noise (Fig. S20A,B), to account for differences in mean protein expression and absolute protein noise among the four promoters. The N-terminal multi-mutant (N-5) showed substantial reduction in normalized mean protein expression (∼56-81% mean expression compared to the wild-type GFP gene) across all promoters (Fig. S20A). Very interestingly, the N-5 mutant also showed substantial reduction in protein noise values, but only when expressed under the bursty promoters *CPA2* and *QCR2* (Fig. S20B). This mutant showed 8% and 11 % reduction in normalized protein noise compared to the wild-type GFP gene when expressed under the *CPA2* and *QCR2* promoters respectively (Mann-Whitney U test, p=1.2×10^-4^ and p=6.7×10^-4^ respectively), but showed much lower change in normalized protein noise when expressed under the regulation of the promoters *RPG1* and *RPL35A* (∼3% increase and ∼4% decrease in CV respectively) (Fig. S20B), despite 36% and 19% decrease in mean expression under the regulation of these non-bursty promoters.

Analysis of the relationship between normalized protein noise and normalized mean expression across all GFP variants revealed very interesting results. The regression line fitted to the plot of normalized mean protein expression against normalized protein noise showed positive slopes only for bursty promoters *CPA2* and *QCR2* (slope of 0.229 and 0.302 respectively, Fig. 6E), like what had been observed before^30,31^. In contrast, the regression line for the promoters *RPL35A* and *RPG1* showed near-zero or negative slope (Slope of -0.131 and 0.0228, respectively, Fig. 6F). Genome-wide analysis of relationship between protein noise and protein synthesis rate per mRNA or tAI for groups of genes with different transcriptional burst frequencies also showed similar trend, with stronger positive relationship between protein noise and translational efficiency for genes with bursty transcription (K_on_ ≤ Q_1_, first quartile of burst frequency) (Fig. S21). These results were in line with the predictions from stochastic simulations, and therefore, suggested that variation in ribosome demand was indeed the molecular link between stochastic fluctuations in mRNA numbers and protein noise.

## Discussion

Taken together, our work provides unique molecular insights into the long-standing observation of positive correlation between translational efficiency and protein noise. We test the long-held assumption that low transcription rate combined with high translational efficiency can generate this positive correlation. We show that the high protein noise in genes with low transcription rate and high translational efficiency primarily stems from higher stochastic variation in mRNA numbers associated with low rate of transcription. We also show that demand for ribosomal machinery underlies translational modulation of transcriptional noise. Thus, our work reveals how the coding sequences of genes can influence expression noise, and how this contribution is dependent on the properties of the promoter. This work also highlights how dynamic properties of a system and shared resources influence gene expression, beyond genetic features such as promoter sequence or presence of specific motifs.

Our results also illustrate how the translation process can decouple mean protein expression and protein noise^27^, with a departure from the traditional inverse relationship between them. The nature and extent of translational decoupling is dictated by the translational efficiency of the coding sequence, as well as by the promoter and its mode of transcription.

There are several other factors besides codon usage that can impact translational efficiency. For example, presence of micro-ORF upstream of the start codon and presence of mRNA secondary structures in the 5’ UTR region can affect translational efficiency ^35,65^. In addition, presence of a stretch of positively-charged residues in the amino acid chain can slow down movement of the growing peptide chain through the ribosomal exit tunnel and can impact translational efficiency ^66^. Thus, the effective translational efficiency for a gene is a complex combination of multiple variables. It remains to be seen how these variables combine with translational bursts and impact expression noise.

In summary, our work identifies variation in ribosome availability as the key feature driving the translational modulation of transcriptional bursts. Such variation happens across organisms and therefore, these results have broad implications for studying protein noise across biological systems. Advancements in single-cell RNA sequencing have enabled us to study mRNA expression noise across diverse cellular processes in different organisms. However, noise at the protein level has more direct impact on cellular phenotypes and therefore, on phenotypic heterogeneity. Thus, a better understanding of how mRNA noise is linked to protein noise will help us better predict phenotypic heterogeneity. Our work further shows that combining estimates of burst parameters from single-cell RNA-sequencing data with tAI values of genes can help predict protein expression noise. As more single-cell RNA-seq data become available, this will greatly facilitate estimation of protein expression noise, and will enhance our understanding of its impact on cellular processes across different biological systems.

## Materials and Methods

### Datasets

The mean mRNA expression levels of yeast genes were calculated from the single-cell RNA-seq data in yeast^40^. Mean mRNA expression values for 5500 genes were obtained from this data. Noise in mRNA expression was calculated following the method described in Parab *et al*.^28^. Briefly, for each gene, mean expression, standard deviation and subsequently, coefficient of variation (CV) values were calculated. In the next step, polynomial functions of different degrees were fitted to the CV vs log-transformed mean mRNA expression plot, and the best fit was chosen. The polynomial function of order 5 was observed to give the best fit and was used for calculating mean-adjusted mRNA expression noise. For each gene, mean-adjusted mRNA noise was calculated by the distance of the vertical line drawn from its CV value to the fitted spline. Protein expression noise values for genes were obtained from Newman *et al.*^15^ and their Distance-to-Median (DM) values, that were corrected for dependence of CV on mean protein expression, in YPD medium were used. Protein noise of 2763 genes were obtained from this data. For translational efficiency, the data from Riba *et al.*^41^ were used, and for each gene, the real protein synthesis rate per mRNA value from their data was considered as its measure of translational efficiency. The tAI values for all yeast genes were calculated using tAI value of each codon in yeast from Sabi *et al.*^67^, according to the method described in Tuller *et al.*^34^.

### Two-state model for bursty transcription

Transcriptional bursts were modeled using a two-state model where a gene transitioned between on- and off-states^42^. The rate of transitions to on- and off-states were considered to follow Poisson distributions individually and thus, the time intervals between two on-transitions or off-transitions were exponentially distributed. The time intervals between successive events (on- or off-switching) were sampled from exponential distributions with rate parameters K_on_ and K_off_ respectively. Transitions to on-state resulted in production of mRNA at a rate β_m_ and translation of these mRNA molecules to proteins at a rate β_p._ These mRNA and protein molecules were considered to undergo removal, resulting from dilution due to cell growth and degradation, at the rates of α_m_ and α_p_ respectively.

The dynamics of transcription and translation were modeled using the following equations.

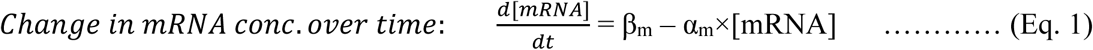

where β_m_ denoted the transcription rate per unit time (or burst size) and α_m_ denoted the removal rate of mRNA due to degradation and dilution.

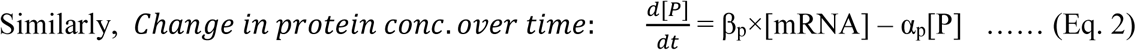

where β_p_ denoted the protein production rate from mRNA and α_p_ denoted the protein removal rate.

Stochastic simulations were performed using Gillespie’s algorithm^43^ to model changes in the numbers of mRNA and protein molecules over time. The behavior of the system was tracked at small discrete time intervals Δt from the initial time point t. These resulted in observations at ‘n+1’ time points t, t+ Δt, t+2×Δt, …, t+n×Δt. Any event of binding or unbinding occurring within a time interval was noted and resulted in changes in transcription rate which eventually led to a change in protein concentration. As the time interval Δt was small, the equations modeling the behavior of the systems was simplified as

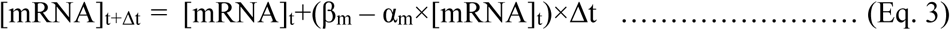

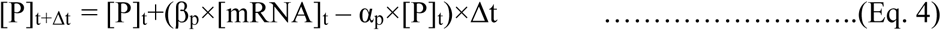

β_m_, α_m_, β_p_ and α_p_ were expressed in appropriate units for further simplification of these equations. The base parameter values for the model were set as: K_on_ = 0.02, K_off_ = 0.2, β_m_ = 10 per unit time, β_p_ = 100 per mRNA molecule per unit time, α_m_ = 0.07 per mRNA molecule per unit time, and α_p_ = 0.007 per protein molecule per unit time. The dynamics of transcription, variation in the mRNA concentration and variation in protein concentration with time were modeled across 10,000 cells. Noise was expressed as coefficient of variation (CV) from the calculation of mean and standard deviation in the protein level across these 10,000 cells across all time points.

To investigate how the mean mRNA expression level and the transcriptional burst frequencies combine with translation rate (β_p_) and impact protein noise, simulations were performed over a wide range of values of the parameters K_on_ and β_m_. The parameter K_on_ was varied between 0.001 and 10, and the parameter β_m_ was varied between 1 to 1000 per unit time. For each value of these parameters, translation rate β_p_ was varied between a range of 1 to 1000 per mRNA molecule per unit time.

### Estimation of burst parameters from single-cell RNA-seq data

The two-state model of gene expression enabled estimation of the parameters of transcriptional bursts from single-cell RNA-sequencing data using a maximum likelihood approach, as described by Kim and Marioni ^44^.

Specifically,

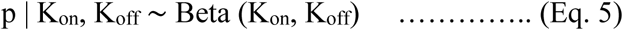

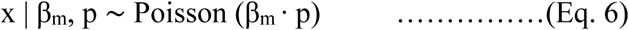

where ‘x’ is the number of RNA transcripts of a given gene within a cell and ‘p’ is an auxiliary variable following a beta distribution. The mean of the parameter ‘p’ is equal to the fraction of time that a gene spends in the active state. The resultant marginal distribution is known as Poisson-Beta distribution, denoted by P (x | K_on_, K_off_, β_m_), and gives the steady-state distribution for the mRNA copy numbers across cells. All parameters are further normalized by α_m_. K_on_ and K_off_ control the shape of the Beta distribution and represent the probability of a gene being in ‘on’ and ‘off’ states, respectively. β_m_ is the mean of Poisson distribution and represents the average rate of gene expression when the gene is in the ‘on’ state.

The yeast single cell RNA-seq data^40^ was given as the input to the model. The two-state model parameters *θ* = (*K_on_*, *K_off_*, β_*m*_) were inferred by maximizing the likelihood of the parameters of Poisson-Beta distribution for a given set of observed data points X. This can be represented as:

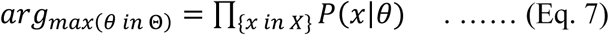

This was further represented as maximization of the log-likelihood, which was equivalent to minimization of the negative log-likelihood:

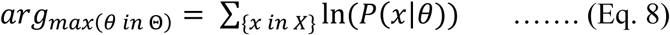

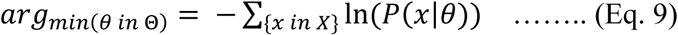

The negative log-likelihood was minimized using L-BFGS-B approach^68^ and the resulting distributions for parameters in *θ* were calculated.

### Mathematical model of translation at single mRNA level for capturing bursty translation

Several mathematical models for the translation process have been described, and they include the TASEP model, the Ribosome Flow Model (RFM) and their variants^45,46,69–72^. Totally Asymmetric Simple Exclusion Process or TASEP model assumes that an mRNA molecule is a one-dimensional lattice through which a ribosome can hop in only one direction from one site to the next, provided the latter is empty.

To capture the dynamics of translation, a simple TASEP-based model with appropriate modifications was developed and bursty translation was incorporated. To start with, translational bursts were assumed to arise due to stochastic transitions of an mRNA molecule between active and inactive states with the rates TL_on_ and TL_off_ respectively. In the active state, translation initiation on the mRNA molecule occurred at a rate TL_init_. The parameters TL_on_ and TL_off_ were assumed to follow Poisson distributions individually and hence, the time-intervals between two subsequent on-states (or off-states) were assumed to follow exponential distributions. The parameter values were chosen within the range of parameter values observed or estimated from experimental datasets, to build a realistic model. For yeast genes, maximum likelihood estimates of K_on_ and K_off_ from single-cell RNA-seq data^40^ were in the ranges of 0.005 to 744, and 0.001 to 1000, respectively. In yeast, the experimental estimates for mRNA synthesis rates ranged from ∼0.01 to 4 per min^73^, and protein synthesis rates per mRNA ranged from 0-13.6 per sec per mRNA^41^. Earlier Estimates for mRNA half-life ranged from 3 mins to 300 mins^74^ but a recent non-invasive measurement of mRNA half-life has found that majority of mRNA molecules in yeast have half-lives of less than 10 mins^64^. The estimates for protein half-life ranged from 2 mins to ∼16000 mins^75^. However, no estimates for TL_on_ and TL_off_ were available. Large values of K_on_, TL_on_, mRNA synthesis rate, protein synthesis rates, mRNA half-life and protein half-life made simulations extremely slow and computationally very demanding, as the model had to track many mRNA molecules and translation initiation events. Therefore, these values were not explored. In addition, genes with very stable mRNA molecules are unlikely to show fluctuation in mRNA numbers and therefore, are not of interest to this study.

As multiple ribosomes could translate a single mRNA molecule at the same time, a second translation initiation happened only when the earlier mRNA bound ribosome had moved by 10 or more codons on the mRNA molecule, thus, accounting for steric interactions between two ribosomes^47,48^. Movement of ribosome through an mRNA molecule depends on the individual codons that are present. Preferred codons for which tRNA molecules are readily available are traversed quickly, whereas the rare codons are read slower. Tuller *et al*.^34^ observed a specific and conserved translational profile across several species where translational efficiency was low at the start of the genes for about 30-50 codons, followed by a gradual increase to the maximum efficiency. Similarly, Weinberg *et al.*^49^ observed a slow rate of translation, although to a different degree, in the first 200 nucleotides of a gene. This enabled us to model ribosome traversal speed in a more general manner rather than focusing on the occurrence of specific codons that would vary from one gene to another. The ribosome traversal speed was modeled using a first order Hill function of the following form

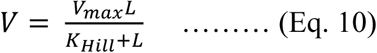

Where, V = traversal speed of the ribosome at codon L in the mRNA molecule, V_max_ = maximum traversal speed of the ribosome in the mRNA molecule, K_Hill_ = constant, and L = position in the mRNA. The values of V_max_ and K_Hill_ were chosen to be 100 codons per min and 30 respectively. All parameters were further varied and tested for model robustness as discussed below.

The value of K_Hill_ influenced the relationship between the traversal speed and the position L, and eventually determined the time taken for a ribosome to completely traverse through a mRNA molecule. High values of K_Hill_ reduced the number of translation initiation events, thus mimicking the scenario of lower translational efficiency. This is because a ribosome took longer time to traverse through the first 10 codons, which was required before the next translation initiation event could take place. A reduction in the value of K_Hill_ allowed higher translation initiation rate (TL_init_) and captured the behavior of the system in case of high translational efficiency. Conversely, increasing the translation initiation rate required the K_Hill_ value to be lower. Thus, the relationship between K_Hill_ and translation initiation rate (TL_init_) was modeled using the equation

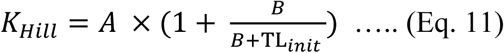

Where A and B were constants. The values of A and B shaped the relationship between K_Hill_ and β_p_, and in our model were chosen as 20 and 5 respectively. Higher translational efficiency led to faster traversal of ribosome through an mRNA molecule and allowed more translation initiation events leading to higher value of β_p_. The model was tested with different values of the parameters K_Hill_, A, B and V_max_ to test for robustness.

In the next step, the model of translation at the single mRNA level was integrated with the two-state model of transcription, and the combined model was utilized for subsequent stochastic simulations. The base parameter values for the model were chosen as: K_on_ = 0.02, K_off_ = 0.2, β_m_ = 4 per minute, TL_on_=0.5, TL_off_=0.5, A=20, B=5, K_Hill_=30, V_max_=100 codons per minute, Length of the gene (L_tot_) = 300 codons, mRNA half-life (HL_mRNA_) = 10 mins, and protein half-life (HL_prot_) = 100 mins.

Every mRNA molecule generated through transcriptional burst was tracked by the translation model at every minute over the life-time of the mRNA molecule, and the protein expression at every time-point was estimated. Gillespie’s algorithm was used for simulations in 1000 cells through which estimates for population-wide mean protein expression and protein noise were derived. The process was repeated for a wide-range of transcriptional and translational burst frequency values, as well as for other parameters of the models, to investigate whether positive correlation between mean protein expression and protein noise could be observed at specific parameter values. The K_on_ parameter was varied between 0.01 and 10, the K_off_ parameter was varied between 0.01 and 10, TL_on_ between 0.01 and 5, TL_off_ between 0.01 and 5, and β_m_ between 0.1 to 20 per min. For each of these parameter values, TL_init_ was varied between 0.1 and 10 per mRNA molecule per minute. The values of mRNA and protein half-lives were stochastically varied by ±10% of the chosen value, as the half-lives of individual mRNA and protein molecules can vary within a cell.

### Modeling and simulation of demand for ribosomal machinery

As multiple mRNA molecules share a common pool of ribosomal machinery for translation, the number of ribosomes available for translating a specific mRNA molecule can vary with time. Several studies have modeled the translation process with finite resources and competition, however, none of them had investigated the impact on protein expression levels or on heterogeneity. Only one earlier study, investing the stochastic nature of translation using mathematical models, predicted that use of rare codons will increase expression heterogeneity^76^, which contradicted the empirical observations.

Competition for a finite pool of ribosomal machinery would impact the translation initiation rate in an mRNA molecule. High availability of ribosomal machinery would lead to high translation initiation rate, whereas a scarcity of ribosomes would lower translation initiation rate. Transcriptional bursts generate stochastic fluctuations in mRNA numbers over time which can lead to a variable demand for ribosomal machinery required for translation. When the number of mRNA molecules of a gene increases suddenly due to a transcriptional burst, the demand for ribosomal machinery increases rapidly. Subsequently, when the mRNA numbers fall, this lowers demand and frees up ribosomes, which can then be utilized for translating mRNA molecules of other genes.

The demand for ribosomal machinery was incorporated in the model of translation so that its effect on translation initiation rate of an mRNA molecule could be captured. For a gene, the number of mRNA molecules produced by transcriptional bursting and the number of ribosomes bound to mRNA molecules were tracked over time. At every time point during simulation, the ribosome demand was modeled using different functions that incorporated the ratio of mRNA numbers at the current time point to the mRNA numbers at the previous time point, the ratio of current number of bound ribosomes to the number of bound ribosomes at the previous time point or a combination of both (parameter ‘av’, Table S1). These included functions where the translation initiation rate at a specific time point was equal to the basal translation initiation rate divided by the ribosome demand, or more complex Hill functions of order 10 (Table S1), that could induce sharp transitions in translation initiation rate when demand was too high. The parameter ‘av’ modeled the ribosome demand at a specific time point during the simulations, and the parameter ‘kc’ defined the threshold value for ribosome demand beyond which the translation initiation rate decreased sharply.

Higher number of mRNA molecules or number of ribosomes bound to mRNA at a time point compared to the previous time point meant lower availability of ribosomal machinery for translation and lower translation initiation rate. Among all functions tested, the one which considered both number of mRNA molecules and number of bound ribosomes at current and previous time points, and modeled the impact of ribosome availability on translation initiation rate through Hill function (Function 14 to Function 19, Table S1) generated the best positive correlation between mean protein expression and protein noise (Fig. S8). The function 16 was used for all subsequent analysis.

The integrated model of bursty transcription and bursty translation incorporating ribosome demand was tested in a wide range of values of several parameters. The impact of variation in values of the parameters A, B and maximum ribosome traversal rate (V_max_) individually on correlation between protein noise and translational efficiency was tested. The parameter A was varied between 10 and 50, B between 1 and 10, V_max_ between 50 and 150. For each of the parameter values, the translation initiation rate (TL_init_) was varied between 0.1 and 10 per mRNA molecule per min. For V_max_ = 50, no protein expression was observed. For the rest of the parameter values, no effect on correlation between translational efficiency and protein noise was observed (Fig. S9-S11).

In the next step, the model was tested with different values of the parameters K_on_, K_off_, TL_on_, TL_off_, and β_m_. Each of these parameters was varied individually and the impact of changes in the translation initiation rate (TL_init_) on protein noise was tested (Fig. 4D,E; Fig. S12-S14). The parameter K_on_ was varied between 0.01 and 10, K_off_ between 0.01 and 10, TL_on_ between 0.01 and 5, TL_off_ between 0.01 and 5, and β_m_ between 0.1 and 20. For each of these parameter values, TL_init_ was varied between 0.1 and 10 per mRNA molecule per min. Higher rates of β_m_ and TL_init_ made simulations extremely slow and computationally very demanding, as the model had to track a large number of mRNA molecules and translation initiation events, respectively. Therefore, these values could not be used for simulation.

Since it was not possible to explore all possible combinations of parameter values due to an extremely large number of combinations, random sampling of parameter values for K_on_, K_off_, TL_on_, TL_off_, β_m_, HL_mRNA_ and HL_protein_ were performed from a specified range of each parameter (Fig. S16). This also enabled us to study how different combinations of values of these parameters impact protein noise, and consequently, the relationship between translational efficiency and protein noise. Fifty random sampling of parameter values were done and for each set of parameter values, the translation rate (β_p_) was varied between 0.2 and 10 per mRNA molecule per min. The parameter K_on_ was sampled in the range of 0.01 to 2, K_off_ between 0.01 and 2, TL_on_ between 0.1 and 5, β_m_ between 0.1 and 10 per min, HL_mRNA_ between 1 and 20 mins, and HL_prot_ between 1 and 200 mins. Simulation results indicated interactions between different parameter values that also impacted the relationship between translational efficiency and protein noise (Fig. S15).

### Construction of GFP-promoter fragment and genomic integration in yeast

*Saccharomyces cerevisiae* BY4741 strain was used for experiments. *Green Fluorescent Protein* (*GFP*) gene integrated in the yeast genome was used as the model system for noise quantification. The gene-promoter construct was made as follows. The *GFP* gene was amplified from the yeast strain where the *PDR5* gene was tagged with *GFP,* and the first two codons ATG and TCT were added with the help of primers (Table S2). For PCR amplification, genomic DNA from yeast was isolated by lithium acetate-SDS method and 2 µl genomic DNA was used as template. Amplification using Q5 DNA polymerase and gene-specific primers was performed, and the PCR product subsequently purified using QIA quick PCR Purification Kit (Qiagen).

The strain BY4741 has a truncated and non-functional *HIS3* gene (*his3Δ1*). Therefore, the gene-promoter construct was inserted into the site of the *HIS3* gene through homologous recombination, and *HIS3MX6* cassette was used as the auxotrophic marker for selection of transformed colonies. The *HIS3MX6* cassette was amplified from yeast strain with *PDR5-GFP* fusion. For genomic integration, the genomic regions upstream and downstream region of the *HIS3* gene, denoted by His3L and His3R respectively (Fig. S17), were amplified and were separately cloned into pUC19 vector with restriction digestion and ligation. The constructs were transformed into chemically competent *E. coli* cells^77^ and the transformed bacterial colonies were selected on LB plates supplemented with 100 µg/ml ampicillin. The promoters, the *GFP* gene and the *HIS3MX6* cassette were subsequently cloned step-by-step through restriction digestion with appropriate enzymes (Fig. S17), followed by ligation with T4 DNA ligase. The constructs were transformed into *E. coli* and were selected on LB plates supplemented with ampicillin. The cloned DNA fragments were verified by restriction digestion of the constructed fragment, as well as through Sanger sequencing. Promoters of four genes, namely, *RPG1*, *RPL35A*, *CPA2* and *QCR2* were cloned upstream of the *GFP* gene (Table S3). The genes *CPA2* and *QCR2* showed high protein noise and the genes *RPG1* and *RPL35A* showed low protein noise in genome-wide noise data of yeast^15^. For cloning promoter sequence of a gene, the region between the start codon of the gene and the stop codon of the previous gene on the same DNA strand was considered (Table S3).

In the next step, the full construct was amplified using His3L forward and His3R reverse primers (Table S2) using Q5 polymerase, and 20 µl of the PCR product was used for transformation into competent yeast cells following the protocol of Gietz and Schiestl^78^. Briefly, yeast cells were first grown to saturation in YPD medium at 30°C overnight, and then were diluted into fresh medium, where they were allowed to grow for 4 hours. Cells were precipitated, and were washed once with 0.1M lithium acetate (LiAc) solution. Cells were resuspended in 50µl of 0.1M LiAc and then 5µl salmon sperm carrier DNA along with 20µl of PCR product were added. Further, 300 µl of PLI solution, comprising of 10% (v/v) 1M LiAC, 10% (v/v) water and 80% (v/v) PEG 3350 (50% w/v), was added to each tube. Cells were subjected to heat shock at 42°C for 40 minutes, following which PLI solution was removed, and the cells were plated on synthetic complete medium supplemented with 2% glucose but without any histidine (SD-His). Colonies with genomic integration were confirmed by colony PCR, and subsequently, were verified by Sanger sequencing.

### Generation of single- and multi-mutant variants of GFP

Seven mutant variants of the *GFP* gene were constructed, out of which three variants contained mutations in the N-terminal region and four variants contained mutations in the C-terminal region. Variants with mutations in the N-terminal or C-terminal regions were targeted to prevent disturbing the catalytic core of the GFP protein. The mutant variants were cloned downstream of the four promoters.

The N-terminal single mutant variants consisted of TCT to TCG (S to S) mutation at codon 2, GGA to GGT (G to G) mutation at codon 4, GAA to GAG (E to E) mutation at codon 5 and GAA to GAC (E to D) mutation at codon 6. The C-terminal single mutants consisted of GAT to GAC (D to D) mutation at codon 234, CTA to CTC (L to L) mutation at codon 236, and AAA to AAG (K to K) mutation at codon 238. The multi-mutant variant at the N-terminal end (denoted by N-5) contained five mutations, namely, TCT to TCG (S to S) at codon 2, AAA to AAG (K to K) at codon 3, GGA to GGT (G to G) at codon 4, GAA to GAG (E to E) at codon 5, and GAA to GAC (E to D) at codon 6. The multi-mutant variant at the C-terminal end (denoted by C-5) contained five mutations, namely, GAT to GAC (D to D) at codon 234, GAA to GAC (E to D) mutation at codon 235, CTA to CTC mutation (L to L) at codon 236, TAC to AAC mutation (Y to N) at codon 237, and AAA to AAG mutation (K to K) at codon 238.

To construct and clone the mutant variants of the GFP gene downstream of the promoter regions, an overlap extension PCR method following the protocol described in Higuchi *et al.*^79^ was used. For generating the mutant variants, primers were designed keeping the target mutation at the center with 15 bases flanking them on each side. In the first amplification step, two fragments were generated. The first fragment was from the 5’ end of Hi3L region till the targeted mutation site in the *GFP* gene, and the second fragment was from the targeted mutation site in the *GFP* gene till the 3’ end region of His3R (Fig. S22). Amplification was carried out using high-fidelity Q5 DNA polymerase. The reaction mixture comprised of 2.5 µl of each primer (10 µM), 10 µl of 5x Q5 reaction buffer, 1 µl of 10 mM dNTPs, 0.5 µl of Q5 DNA polymerase (New England Biolabs) and molecular biology grade water to a total volume of 50 µl. The PCR program consisted of incubation at 98 °C for 1 min (initial denaturation) followed by 30 cycles of 98°C for 10 sec (denaturation), 65°C for 30 sec (annealing), 72°C for 1 min 30 sec (extension), and a final extension at 72°C for 10 min.

The products after amplification were treated with 1 µl of ExoSAP-IT PCR product cleanup reagent (Thermo Fisher Scientific) and 0.5 µl DpnI (New England Biolabs) for 1 hr at 37°C, followed by deactivation for 20 min at 80 °C. The products were then purified using QIAquick PCR Purification Kit (Qiagen) and quantified using Qubit 4 Fluorometer (Thermo Fisher Scientific).

As the molecular weight of the amplified fragments were variable, equimolar concentration of the fragments from the previous amplification step were used in the next step. For each reaction, 0.5 to 1 picomole of each purified fragment were used as templates. For assembly, the reaction mixture comprised of equimolar content of two template fragments, 10 µl of 5x Q5 reaction buffer, 1 µl of 10 mM dNTPs, 0.5 µl of Q5 DNA polymerase (New England Biolabs) and molecular biology grade water to a total volume of 45 µl, without any primer. The program consisted of incubation at 98 °C for 1 min (initial denaturation) followed by 10 cycles of 98°C for 10 sec (denaturation), 60°C for 30 sec (annealing) and 72°C for 1 min 30 sec (extension). On completion of 10^th^ cycle, 2.5 µl of forward and reverse primers (10 µM stock of each) were added, and the program was run for next 25 cycles by raising the annealing temperature to 72°C. The reaction was terminated by a final extension at 72°C for 10 min.

The assembled mutant constructs were then directly transformed into competent yeast cells following the LiAC method as described above, and the transformants were selected on SD-His plates. The clones were confirmed by colony PCR and Sanger sequencing.

### Measurement of expression noise using flow cytometry

Three clones of wild-type GFP gene and each mutant variant were picked for noise measurement in flow cytometry. Yeast cultures were grown in SCD medium (0.67% YNB + 0.079% complete synthetic supplement + 2% glucose) at 30℃ with shaking at 220 rpm. Yeast cells from glycerol stock were inoculated in SCD medium, were grown for 24 hours, and were then diluted 1:100 in fresh medium. This process was repeated one more time. The cells were then diluted 1:50 in fresh SCD and were grown for 4 hours. In the next step, cells were centrifuged and washed twice with 1X PBS (Phosphate Buffered Saline) buffer, and were finally resuspended in 1X PBS buffer for flow cytometry. In flow cytometry, the data were acquired for 2,00,000 cells for each sample in a BD LSR Fortessa cell analyzer (Fig. S18). To minimize heterogeneity in signal due to heterogeneity in cell size and complexity, a small homogeneous subset of cells with similar size and complexity parameters (FSC-A and SSC-A) were filtered. Ellipses with different lengths of major and minor axes were fitted to the FSC-A vs SSC-A plot, and the best fit that covered the most of homogeneous cells was chosen (Fig. S19). This also filtered out cell aggregates or budding cells which can have higher GFP signal than single cells. The data from these cells were then used for calculation of expression noise. The data were analyzed using custom R codes based on codes from Silander *et al.*^14^. Normalized mean expression and normalized noise value for a strain in an experiment was calculated by dividing the mean expression and noise values with the corresponding values for the wild-type variant in that experiment.

## Acknowledgments

The authors are thankful to the members of the lab of Dr. Gayatri Mukherjee, School of Medical Science and Technology, IIT Kharagpur for help with flow cytometry.

## Author contributions

Conceptualization: RD

Methodology: SP, UR, RD

Investigation: SP, UR, RD

Visualization: SP, UR, RD

Funding acquisition: RD

Project administration: RD

Supervision: RD

Writing – original draft: SP, UR, RD

Writing – review & editing: SP, UR, RD

## Data availability

Flow cytometry data are available in github: https://github.com/riddhimandhar/TranslationModNoise and dryad: https://doi.org/10.5061/dryad.tb2rbp06x

https://datadryad.org/stash/share/D8AhKWZIiouA5HNG9fjUytzetVlz_AOCagXsKcID0WI

## Code availability

All codes for data analysis and models are available in github: https://github.com/riddhimandhar/TranslationModNoise

## Supplementary Materials

Figs. S1 to S21 Tables S1 to S3

## References

1. N. Q. Balaban, J. Merrin, R. Chait, L. Kowalik, S. Leibler, Bacterial persistence as a phenotypic switch. Science 305, 1622–1625 (2004). doi: 10.1126/science.1099390.

2. Y. Wakamoto, N. Dhar, R. Chait, K. Schneider, F. Signorino-Gelo, S. Leibler, J. D. McKinney, Dynamic persistence of antibiotic-stressed mycobacteria. Science 339, 91–95 (2013). doi: 10.1126/science.1229858.

3. A. Raj, S. A. Rifkin, E. Andersen, A. van Oudenaarden, Variability in gene expression underlies incomplete penetrance. Nature 463, 913–918 (2010). doi: 10.1038/nature08781.

4. A. Burga, M. O. Casanueva, B. Lehner, Predicting mutation outcome from early stochastic variation in genetic interaction partners. Nature 480, 250–253 (2011). doi: 10.1038/nature10665.

5. A. Eldar, V. K. Chary, P. Xenopoulos, M. E. Fontes, O. C. Losón, J. Dworkin, P. J. Piggot, M. B. Elowitz, Partial penetrance facilitates developmental evolution in bacteria. Nature 460, 510–514 (2009). doi: 10.1038/nature08150.

6. Y. Lu, K. Bohn-Wippert, P. J. Pazerunas, J. M. Moy, H. Singh, R. D. Dar, Screening for gene expression fluctuations reveals latency-promoting agents of HIV. Proc Natl Acad Sci USA 118, e2012191118 (2021). doi: 10.1073/pnas.2012191118.

7. M. A. Coomer, L. Ham, M. P. H. Stumpf, Noise distorts the epigenetic landscape and shapes cell-fate decisions. Cell Syst. 13, 83–102.e6 (2022). doi: 10.1016/j.cels.2021.09.002.

8. A. Nguyen, M. Yoshida, H. Goodarzi, S. F. Tavazoie, Highly variable cancer subpopulations that exhibit enhanced transcriptome variability and metastatic fitness. Nat Commun. 7, 11246 (2016). doi: 10.1038/ncomms11246.

9. A. Sharma, E. Merritt, X. Hu, A. Cruz, C. Jiang, H. Sarkodie, Z. Zhou, J. Malhotra, G. M. Riedlinger, S. De, Non-Genetic Intra-Tumor Heterogeneity Is a Major Predictor of Phenotypic Heterogeneity and Ongoing Evolutionary Dynamics in Lung Tumors. Cell Rep. 29, 2164–2174.e5 (2019). doi: 10.1016/j.celrep.2019.10.045.

10. B. L. Emert, C. J. Cote, E. A. Torre, I. P. Dardani, C. L. Jiang, N. Jain, S. M. Shaffer, A. Raj, Variability within rare cell states enables multiple paths toward drug resistance. Nat Biotechnol. 39, 865–876 (2021). doi: 10.1038/s41587-021-00837-3.

11. I. Golding, J. Paulsson, S. M. Zawilski, E. C. Cox, Real-time kinetics of gene activity in individual bacteria. Cell 123,1025–1036 (2005). doi: 10.1016/j.cell.2005.09.031.

12. L. Cai, N. Friedman, X. S. Xie, Stochastic protein expression in individual cells at the single molecule level. Nature 440, 358–62 (2006). doi: 10.1038/nature04599.

13. A. Raj, C. S. Peskin, D. Tranchina, D. Y. Vargas, S. Tyagi, Stochastic mRNA synthesis in mammalian cells. PLoS Biol. 4, e309 (2006). doi: 10.1371/journal.pbio.0040309.

14. O. K. Silander, N. Nikolic, A. Zaslaver, A. Bren, I. Kikoin, U. Alon, M. Ackermann, A genome-wide analysis of promoter-mediated phenotypic noise in *Escherichia coli*. PLoS Genet. 8, e1002443 (2012). doi: 10.1371/journal.pgen.1002443.

15. J. R. Newman, S. Ghaemmaghami, J. Ihmels, D. K. Breslow, M. Noble, J. L. DeRisi, J. S. Weissman, Single-cell proteomic analysis of *S. cerevisiae* reveals the architecture of biological noise. Nature, 441, 840–846 (2006). doi: 10.1038/nature04785.

16. A. Bar-Even, J. Paulsson, N. Maheshri, M. Carmi, E. O’Shea, Y. Pilpel, N. Barkai, Noise in protein expression scales with natural protein abundance. Nat Genet. 38, 636–643 (2006). doi: 10.1038/ng1807.

17. D. M. Suter, N. Molina, D. Gatfield, K. Schneider, U. Schibler, F. Naef, Mammalian genes are transcribed with widely different bursting kinetics. Science 332, 472–4 (2011). doi: 10.1126/science.1198817.

18. R. D. Dar, B. S. Razooky, A. Singh, T. V. Trimeloni, J. M. McCollum, C. D. Cox, M. L. Simpson, L. S. Weinberger, Transcriptional burst frequency and burst size are equally modulated across the human genome. Proc Natl Acad Sci USA 109, 17454–17459 (2012). doi: 10.1073/pnas.1213530109

19. I. Tirosh, A. Weinberger, M. Carmi, N. Barkai, A genetic signature of interspecies variations in gene expression. Nat Genet. 38, 830–834 (2006). doi: 10.1038/ng1819.

20. J. M. Raser, E. K. O’Shea, Control of stochasticity in eukaryotic gene expression. Science 304, 1811–1814 (2004). doi: 10.1126/science.1098641.

21. C. N. Ravarani, G. Chalancon, M. Breker, N. S. de Groot, M. M. Babu, Affinity and competition for TBP are molecular determinants of gene expression noise. Nat Commun. 7, 10417 (2016). doi: 10.1038/ncomms10417.

22. I. Tirosh, N. Barkai, Two strategies for gene regulation by promoter nucleosomes. Genome Res. 18, 1084–1091 (2008). doi: 10.1101/gr.076059.108.

23. J. K. Choi, Y. J. Kim, Intrinsic variability of gene expression encoded in nucleosome positioning sequences. Nat Genet. 41, 498–503 (2009). doi: 10.1038/ng.319.

24. X. Chen, J. Zhang, The Genomic Landscape of Position Effects on Protein Expression Level and Noise in Yeast. Cell Syst. 2, 347–354 (2016). doi: 10.1016/j.cels.2016.03.009.

25. A. J. Faure, J. M. Schmiedel, B. Lehner, Systematic Analysis of the Determinants of Gene Expression Noise in Embryonic Stem Cells. Cell Syst. 5, 471–484.e4 (2017). doi: 10.1016/j.cels.2017.10.003.

26. B. T. Donovan, A. Huynh, D. A. Ball, H. P. Patel, M. G. Poirier, D. R. Larson, M. L. Ferguson, T. L. Lenstra. Live-cell imaging reveals the interplay between transcription factors, nucleosomes, and bursting. EMBO J. 38, e100809 (2019). doi: 10.15252/embj.2018100809.

27. K. Loell, Y. Wu, M. V. Staller, B. Cohen, Activation domains can decouple the mean and noise of gene expression. Cell Rep. 40, 111118 (2022). doi: 10.1016/j.celrep.2022.111118.

28. L. Parab, S. Pal, R. Dhar, Transcription factor binding process is the primary driver of noise in gene expression. PLoS Genet. 18, e1010535 (2022). doi: 10.1371/journal.pgen.1010535.

29. Y. Taniguchi, P. J. Choi, G. W. Li, H. Chen, M. Babu, J. Hearn, A. Emili, X. S. Xie, Quantifying *E. coli* proteome and transcriptome with single-molecule sensitivity in single cells. Science 329, 533–538 (2010). doi: 10.1126/science.1188308.

30. E. M. Ozbudak, M. Thattai, I. Kurtser, A. D. Grossman, A. van Oudenaarden, Regulation of noise in the expression of a single gene. Nat Genet. 31, 69–73 (2002). doi: 10.1038/ng869.

31. W. J. Blake, M. Kaern, C. R. Cantor, J. J. Collins, Noise in eukaryotic gene expression. Nature 422, 633–637 (2003). doi: 10.1038/nature01546.

32. R. Salari, D. Wojtowicz, J. Zheng, D. Levens, Y. Pilpel, T. M. Przytycka, Teasing apart translational and transcriptional components of stochastic variations in eukaryotic gene expression. PLoS Comput Biol. 8, e1002644 (2012). doi: 10.1371/journal.pcbi.1002644.

33. M. dos Reis, R. Savva, L. Wernisch, Solving the riddle of codon usage preferences: a test for translational selection. Nucleic Acids Res. 32, 5036–44 (2004). doi: 10.1093/nar/gkh834.

34. T. Tuller, A. Carmi, K. Vestsigian, S. Navon, Y. Dorfan, J. Zaborske, T. Pan, O. Dahan, I. Furman, Y. Pilpel, An evolutionarily conserved mechanism for controlling the efficiency of protein translation. Cell 141, 344–354 (2010). doi: 10.1016/j.cell.2010.03.031.

35. H. W. Wu, E. Fajiculay, J. F. Wu, C. S. Yan, C. P. Hsu, S. H. Wu, Noise reduction by upstream open reading frames. Nat Plants. 8, 474–480 (2022). doi: 10.1038/s41477-022-01136-8.

36. H. B. Fraser, A. E. Hirsh, G. Giaever, J. Kumm, M. B. Eisen, Noise minimization in eukaryotic gene expression. PLoS Biol. 2, e137 (2004). doi: 10.1371/journal.pbio.0020137.

37. Y. Pilpel, “Noise in biological systems: pros, cons, and mechanisms of control” in Yeast Systems Biology (Humana Press, 2011) pp. 407–25. doi: 10.1007/978-1-61779-173-4_23.

38. B. Wu, C. Eliscovich, Y. J. Yoon, R. H. Singer, Translation dynamics of single mRNAs in live cells and neurons. Science 352, 1430–1435 (2016). doi: 10.1126/science.aaf1084.

39. N. M. Livingston, J. Kwon, O. Valera, J. A. Saba, N. K. Sinha, P. Reddy, B. Nelson, C. Wolfe, T. Ha, R. Green, J. Liu, B. Wu, Bursting translation on single mRNAs in live cells. Mol Cell. 83, 2276–2289.e11 (2023). doi: 10.1016/j.molcel.2023.05.019.

40. M. Nadal-Ribelles, S. Islam, W. Wei, P. Latorre, M. Nguyen, E. de Nadal, F. Posas, L. M. Steinmetz, Sensitive high-throughput single-cell RNA-seq reveals within-clonal transcript correlations in yeast populations. Nat Microbiol. 4, 683–692 (2019). doi: 10.1038/s41564-018-0346-9.

41. A. Riba, N. Di Nanni, N. Mittal, E. Arhné, A. Schmidt, M. Zavolan, Protein synthesis rates and ribosome occupancies reveal determinants of translation elongation rates. Proc Natl Acad Sci USA 116:15023–15032 (2019). doi: 10.1073/pnas.1817299116.

42. J. Peccoud, B. Ycart, Markovian modeling of gene-product synthesis. Theor. Popul. Biol. 48, 222–234 (1995).

43. D. T. Gillespie, Exact stochastic simulation of coupled chemical reactions. J. Phys. Chem. 81, 2340–2361 (1977).

44. J. K. Kim, J. C. Marioni, Inferring the kinetics of stochastic gene expression from single-cell RNA-sequencing data. Genome Biol. 14, R7 (2013). doi: 10.1186/gb-2013-14-1-r7.

45. R.K.P. Zia, J. J. Dong, B. Schmittmann, Modeling Translation in Protein Synthesis with TASEP: A Tutorial and Recent Developments. J Stat Phys 144, 405–428 (2011).

46. D. E. Andreev, M. Arnold, S. J. Kiniry, G. Loughran, A. M. Michel, D. Rachinskii, P. V. Baranov, TASEP modelling provides a parsimonious explanation for the ability of a single uORF to derepress translation during the integrated stress response. Elife 7, e32563 (2018). doi: 10.7554/eLife.32563.

47. J. A. Steitz, Polypeptide chain initiation: nucleotide sequences of the three ribosomal binding sites in bacteriophage R17 RNA. Nature 224, 957–964 (1969). doi: 10.1038/224957a0.

48. N. T. Ingolia, S. Ghaemmaghami, J. R. Newman, J. S. Weissman, Genome-wide analysis in vivo of translation with nucleotide resolution using ribosome profiling. Science 324, 218–223 (2009). doi: 10.1126/science.1168978.

49. D. E. Weinberg, P. Shah, S. W. Eichhorn, J. A. Hussmann, J. B. Plotkin, D. P. Bartel. Improved Ribosome-Footprint and mRNA Measurements Provide Insights into Dynamics and Regulation of Yeast Translation. Cell Rep. 14, 1787–1799 (2016). doi: 10.1016/j.celrep.2016.01.043.

50. C. L. Barrington, G. Galindo, A. L. Koch, E. R. Horton, E. J. Morrison, S. Tisa, T. J. Stasevich, O. S. Rissland. Synonymous codon usage regulates translation initiation. Cell Rep. 42, 113413 (2023). doi: 10.1016/j.celrep.2023.113413.

51. D. Chu, D. J. Barnes, T. von der Haar, The role of tRNA and ribosome competition in coupling the expression of different mRNAs in *Saccharomyces cerevisiae*. Nucleic Acids Res. 39, 6705–14 (2011). doi: 10.1093/nar/gkr300.

52. P. M. Caveney, S. E. Norred, C. W. Chin, J. B. Boreyko, B. S. Razooky, S. T. Retterer, C. P. Collier, M. L. Simpson, Resource Sharing Controls Gene Expression Bursting. ACS Synth Biol. 6, 334–343 (2017). doi: 10.1021/acssynbio.6b00189.

53. Y. Liu, A code within the genetic code: codon usage regulates co-translational protein folding. Cell Commun Signal. 18, 145 (2020). doi: 10.1186/s12964-020-00642-6.

54. P. S. Spencer, E. Siller, J. F. Anderson, J. M. Barral, Silent substitutions predictably alter translation elongation rates and protein folding efficiencies. J Mol Biol. 422, 328–335 (2012). doi: 10.1016/j.jmb.2012.06.010.

55. E. P. O’Brien, M. Vendruscolo, C. M. Dobson, Kinetic modelling indicates that fast-translating codons can coordinate cotranslational protein folding by avoiding misfolded intermediates. Nat Commun.5, 2988 (2014). doi: 10.1038/ncomms3988.

56. F. Trovato, E. P. O’Brien, Fast Protein Translation Can Promote Co- and Posttranslational Folding of Misfolding-Prone Proteins. Biophys J. 112, 1807–1819 (2017). doi: 10.1016/j.bpj.2017.04.006.

57. C. L. Simms, L. L. Yan, H. S. Zaher, Ribosome Collision Is Critical for Quality Control during No-Go Decay. Mol Cell. 68, 361–373.e5 (2017). doi: 10.1016/j.molcel.2017.08.019.

58. S. Juszkiewicz, G. Slodkowicz, Z. Lin, P. Freire-Pritchett, S. Y. Peak-Chew, R. S. Hegde, Ribosome collisions trigger cis-acting feedback inhibition of translation initiation. Elife 9, e60038 (2020). doi: 10.7554/eLife.60038.

59. K. Saito, H. Kratzat, A. Campbell, R. Buschauer, A. M. Burroughs, O. Berninghausen, L. Aravind, R. Green, R. Beckmann, A. R. Buskirk, Ribosome collisions induce mRNA cleavage and ribosome rescue in bacteria. Nature 603, 503–508 (2022). doi: 10.1038/s41586-022-04416-7.

60. E. Mordret, O. Dahan, O. Asraf, R. Rak, A. Yehonadav, G. D. Barnabas, J. Cox, T. Geiger, A. B. Lindner, Y. Pilpel, Systematic Detection of Amino Acid Substitutions in Proteomes Reveals Mechanistic Basis of Ribosome Errors and Selection for Translation Fidelity. Mol Cell. 75, 427–441.e5 (2019). doi: 10.1016/j.molcel.2019.06.041.

61. M. Sun, J. Zhang, Preferred synonymous codons are translated more accurately: Proteomic evidence, among-species variation, and mechanistic basis. Sci Adv. 8, eabl9812 (2022). doi: 10.1126/sciadv.abl9812.

62. P. Dave, G. Roth, E. Griesbach, D. Mateju, T. Hochstoeger, J. A. Chao. Single-molecule imaging reveals translation-dependent destabilization of mRNAs. Mol Cell. 83, 589–606.e6 (2023). doi: 10.1016/j.molcel.2023.01.013.

63. V. Presnyak, N. Alhusaini, Y. H. Chen, S. Martin, N. Morris, N. Kline, S. Olson, D. Weinberg, K. E. Baker, B. R. Graveley, J. Coller. Codon optimality is a major determinant of mRNA stability. Cell 160, 1111–24 (2015). doi: 10.1016/j.cell.2015.02.029.

64. L. Y. Chan, C. F. Mugler, S. Heinrich, P. Vallotton, K. Weis. Non-invasive measurement of mRNA decay reveals translation initiation as the major determinant of mRNA stability. Elife 7, e32536 (2018). doi: 10.7554/eLife.32536.

65. K. Leppek, R. Das, M. Barna, Functional 5’ UTR mRNA structures in eukaryotic translation regulation and how to find them. Nat Rev Mol Cell Biol. 19, 158–174 (2018). doi: 10.1038/nrm.2017.103.

66. C. A. Charneski, L. D. Hurst, Positively charged residues are the major determinants of ribosomal velocity. PLoS Biol. 11, e1001508 (2013). doi: 10.1371/journal.pbio.1001508.

67. R. Sabi, T. Tuller, Modelling the efficiency of codon-tRNA interactions based on codon usage bias. DNA Res. 21, 511–526 (2014). doi: 10.1093/dnares/dsu017.

68. D. C. Liu, J. Nocedal, On the limited memory BFGS method for large scale optimization. Mathematical Programming 45, 503–528 (1989).

69. S. Reuveni, I. Meilijson, M. Kupiec, E. Ruppin, T. Tuller, Genome-scale analysis of translation elongation with a ribosome flow model. PLoS Comput Biol. 7, e1002127 (2011). doi: 10.1371/journal.pcbi.1002127.

70. L. J. Cook, R. K. Zia, B. Schmittmann, Competition between multiple totally asymmetric simple exclusion processes for a finite pool of resources. Phys Rev E Stat Nonlin Soft Matter Phys. 80, 031142 (2009). doi: 10.1103/PhysRevE.80.031142.

71. C. A. Brackley, M. C. Romano, M. Thiel, The dynamics of supply and demand in mRNA translation. PLoS Comput Biol. 7, e1002203 (2011). doi: 10.1371/journal.pcbi.1002203.

72. A. Jain, M. Margaliot, A. K. Gupta, Large-scale mRNA translation and the intricate effects of competition for the finite pool of ribosomes. J R Soc Interface 19, 20220033 (2022). doi: 10.1098/rsif.2022.0033.

73. C. Miller, B. Schwalb, K. Maier, D. Schulz, S. Dümcke, B. Zacher, A. Mayer, J. Sydow, L. Marcinowski, L. Dölken, D. E. Martin, A. Tresch, P. Cramer, Dynamic transcriptome analysis measures rates of mRNA synthesis and decay in yeast. Mol Syst Biol. 7, 458 (2011). doi: 10.1038/msb.2010.112.

74. J. V. Geisberg, Z. Moqtaderi, X. Fan, F. Ozsolak, K. Struhl, Global analysis of mRNA isoform half-lives reveals stabilizing and destabilizing elements in yeast. Cell 156, 812–824 (2014). doi: 10.1016/j.cell.2013.12.026.

75. A. Belle, A. Tanay, L. Bitincka, R. Shamir, E. K. O’Shea, Quantification of protein half-lives in the budding yeast proteome. Proc Natl Acad Sci USA 103, 13004–13009 (2006). doi: 10.1073/pnas.0605420103.

76. A. Garai, D. Chowdhury, T. V. Ramakrishnan, Fluctuations in protein synthesis from a single RNA template: stochastic kinetics of ribosomes. Phys Rev E Stat Nonlin Soft Matter Phys. 79, 011916 (2009). doi: 10.1103/PhysRevE.79.011916.

77. A. Nishimura, M. Morita, Y. Nishimura, Y Sugino, A rapid and highly efficient method for preparation of competent Escherichia coli cells. Nucleic Acids Res. 18, 6169 (1990). doi: 10.1093/nar/18.20.6169.

78. R. D. Gietz, R. H. Schiestl, High-efficiency yeast transformation using the LiAc/SS carrier DNA/PEG method. Nat Protoc. 2, 31–34 (2007). doi: 10.1038/nprot.2007.13.

79. R. Higuchi, B. Krummel, R. K. Saiki, A general method of in vitro preparation and specific mutagenesis of DNA fragments: study of protein and DNA interactions. Nucleic Acids Res. 16, 7351–7367 (1988). doi: 10.1093/nar/16.15.7351.

