## Supplementary material for "Ribosome demand links transcriptional bursts to protein expression noise"

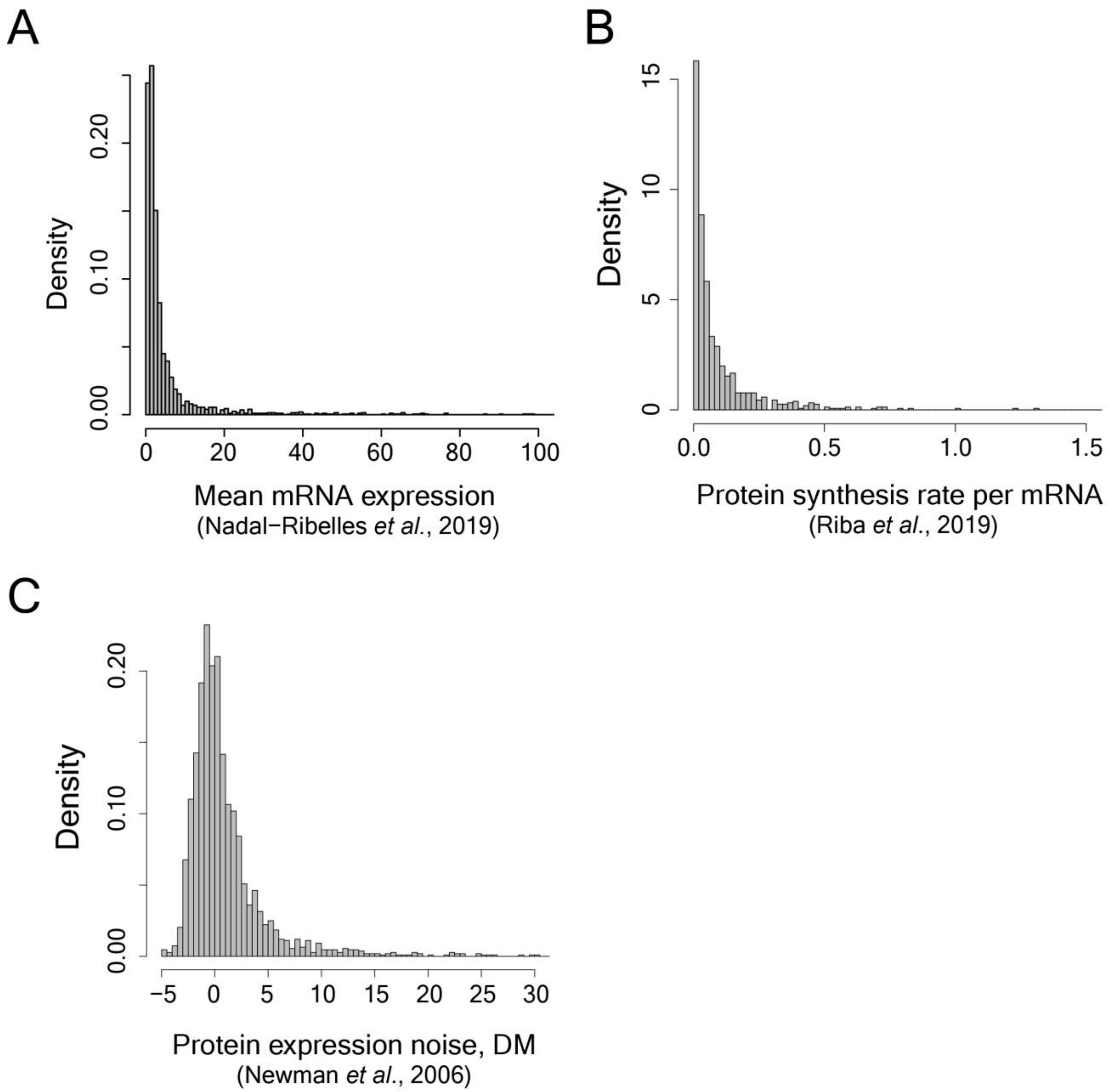


**Fig. S1.**

Distribution of **(A)** mean-mRNA expression from Nadal-Ribelles *et al*.[40], **(B)** protein synthesis rate per mRNA from Riba *et al*.[41], and **(C)** protein expression noise (DM) from Newman *et al*.[15]**.**

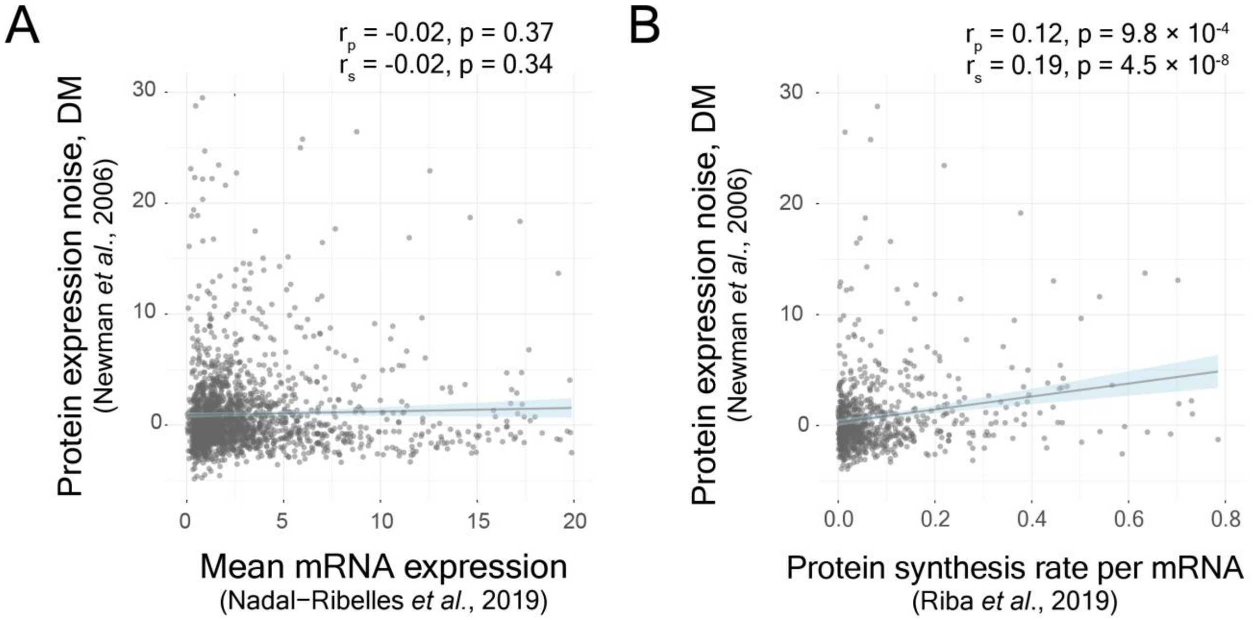


**Fig. S2.**

Correlation of protein expression noise, DM from Newman *et al.*[15], **(A)** with mean mRNA expression, calculated from Nadal-Ribelles *et al.*[40], and **(B)** with protein synthesis rate per mRNA from Riba *et al.*[41]

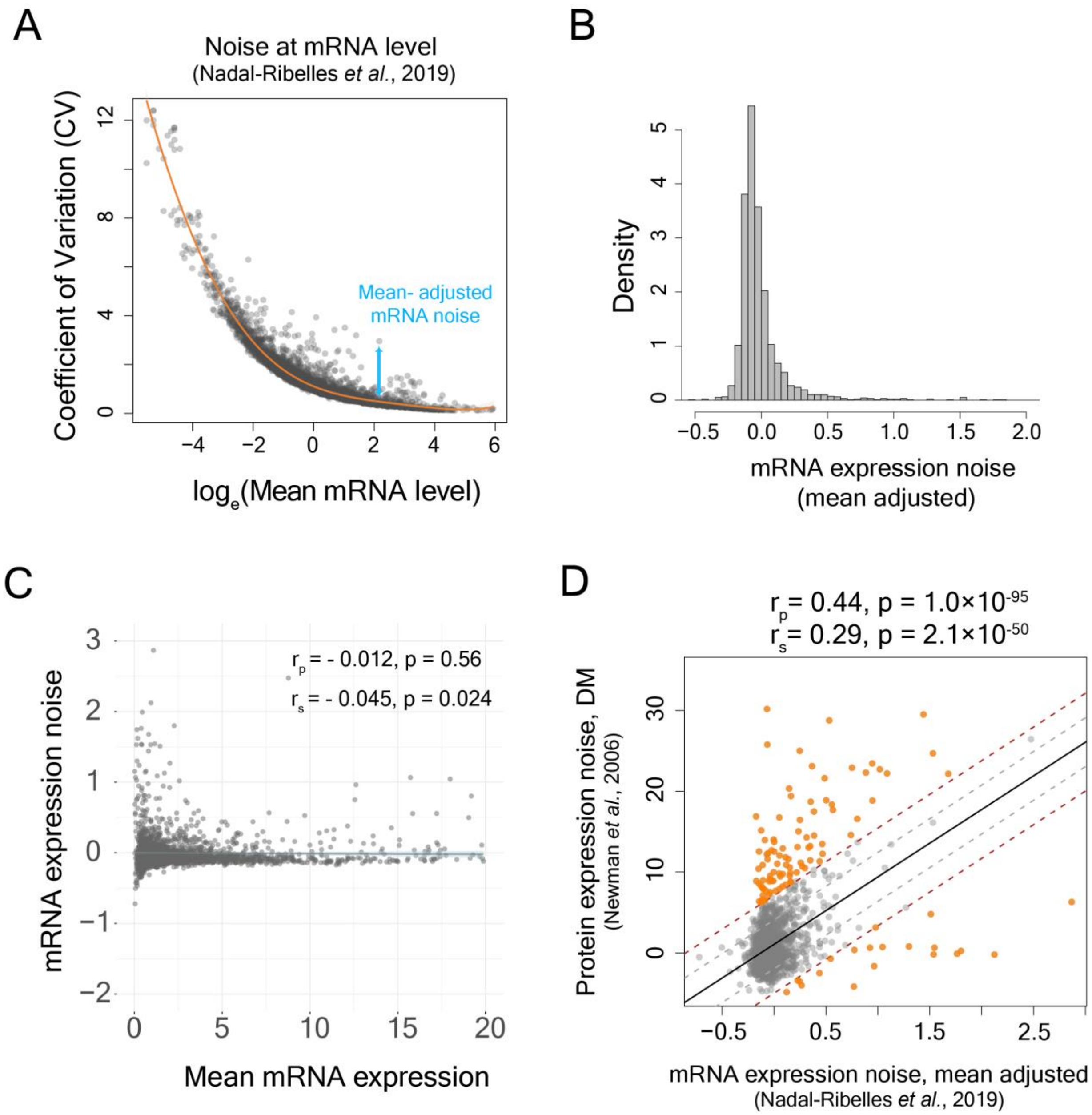


**Fig. S3.**

**(A)** Calculation of mean-adjusted mRNA expression noise from the plot of coefficient of variation vs ln(mean mRNA expression), calculated from[40] **(B)** Distribution of mean-adjusted mRNA expression noise for 5500 genes in yeast **(C)** Correlation between mean RNA expression and mRNA expression noise, calculated from [40] **(D)** Correlation between protein expression noise, DM [15], and mean-adjusted mRNA expression noise. The solid black line represents the linear regression line between mean-adjusted mRNA expression noise and protein expression noise (DM). The grey dotted lines represent ±1 s.d. lines and the brown dotted lines represent ±2 s.d. lines. The orange points show genes which fall outside the ±2 s.d. lines.

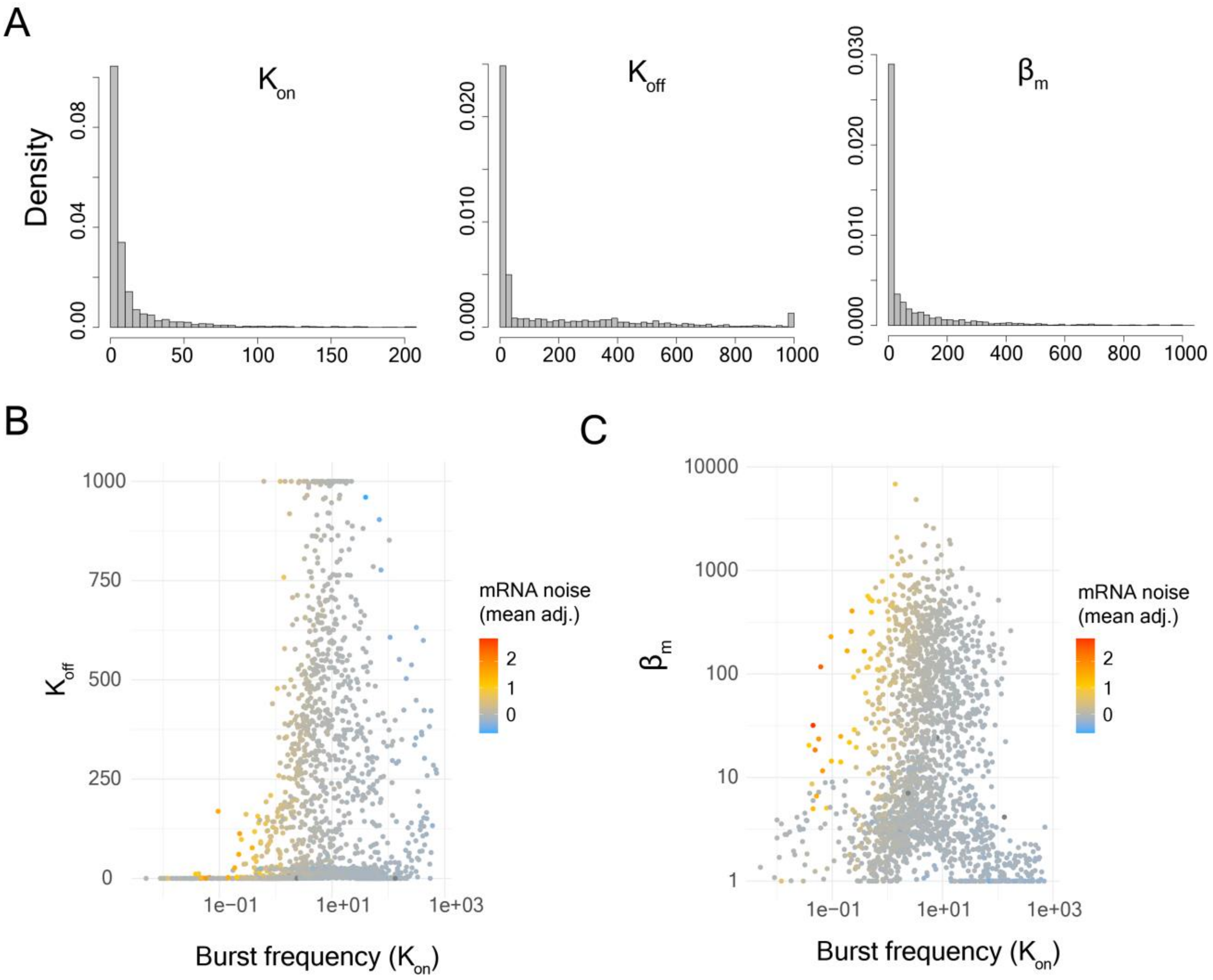


**Fig. S4.**

**(A)** Distribution of values of parameters of transcriptional bursts ($K_{on}$, $K_{off}$ and $\beta_m$) for yeast genes estimated from [40]. **(B)** Relationship between $K_{on}$, $K_{off}$ and mean-adjusted mRNA expression noise for yeast genes. **(C)** Relationship between $K_{on}$, $\beta_m$ and mean-adjusted mRNA expression noise for yeast genes.

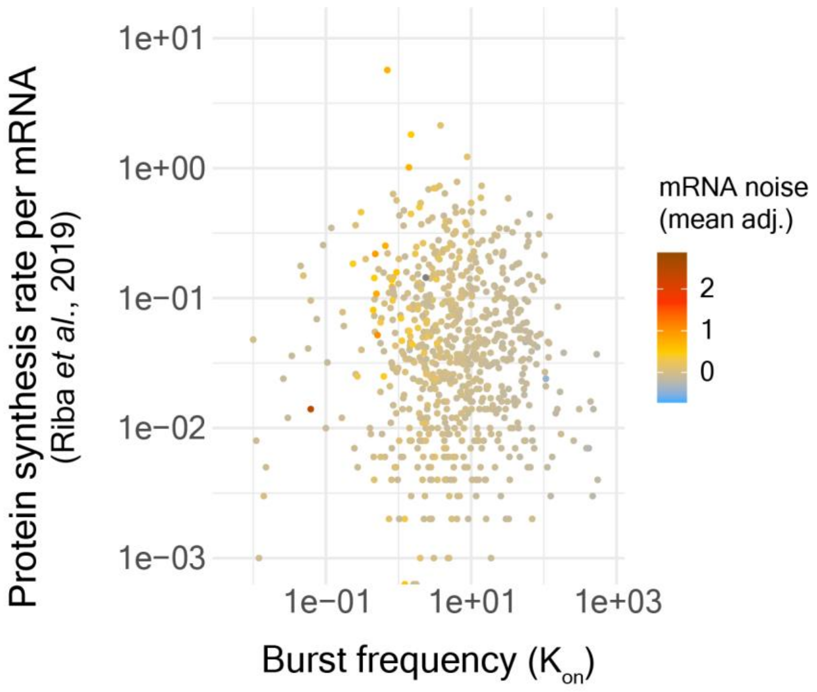


**Fig. S5.**

Relationship between transcriptional burst frequency ($K_{on}$), protein synthesis rate per mRNA[41], and mean-adjusted mRNA expression noise.

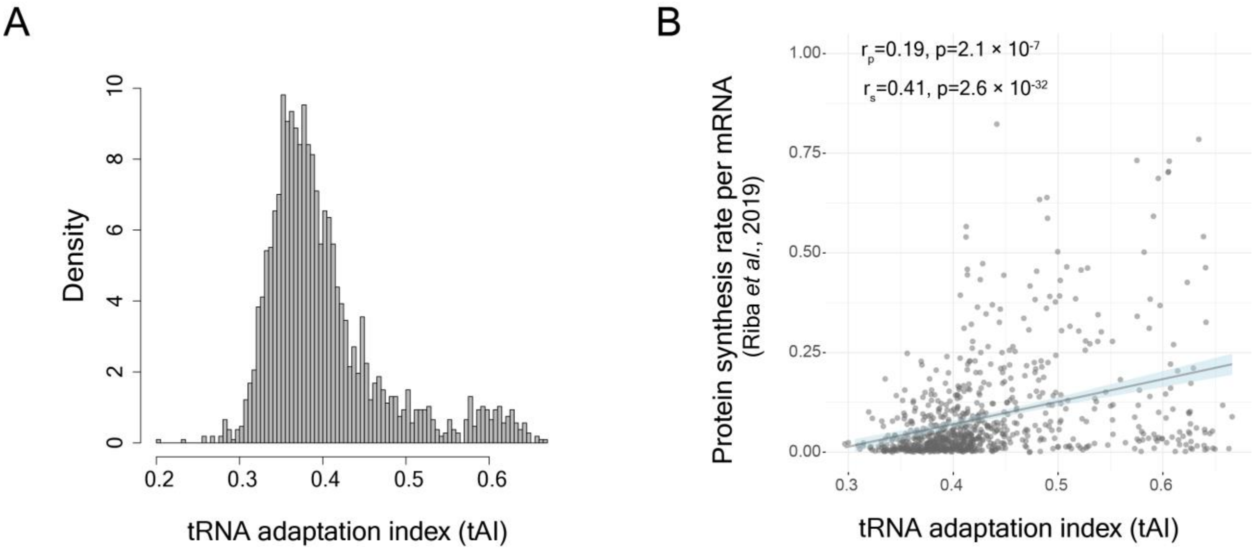


**Fig. S6.**

**(A)** Distribution of tRNA adaptation index (tAI) values of yeast genes calculated from [67] according to the method described in [34]. **(B)** Correlation between tAI and protein synthesis rate per mRNA[41] for yeast genes.

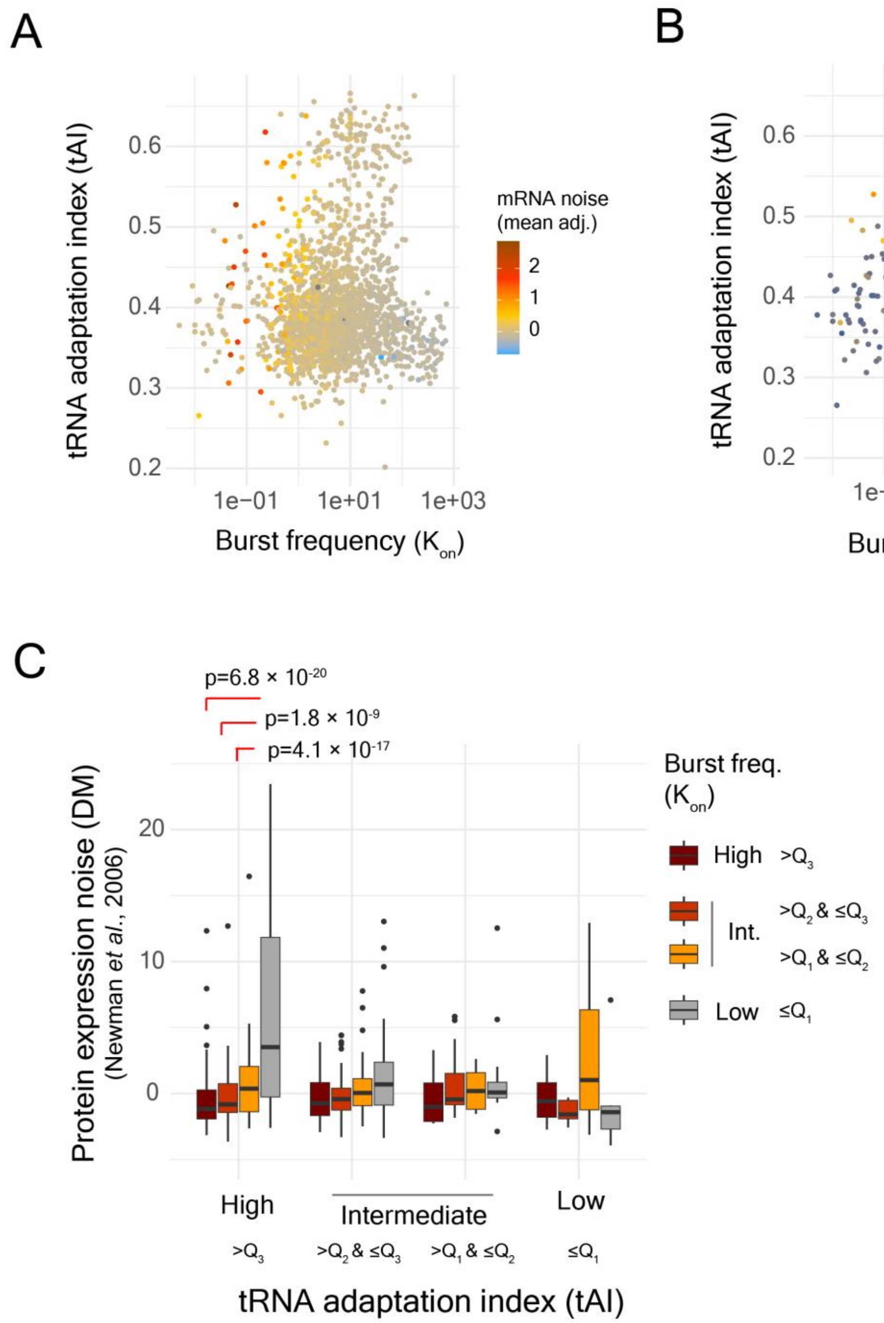


**Fig. S7.**

**(A)** Relationship between transcriptional burst frequency ($K_{on}$), tAI, and mean-adjusted mRNA expression noise **(B)** Relationship between transcriptional burst frequency ($K_{on}$), tAI, and protein expression noise (DM)[15] **(C)** Protein expression noise (DM) in 16 classes of yeast genes classified based on the quartiles of transcriptional burst frequency ($K_{on}$) and then by the quartiles of tAI.

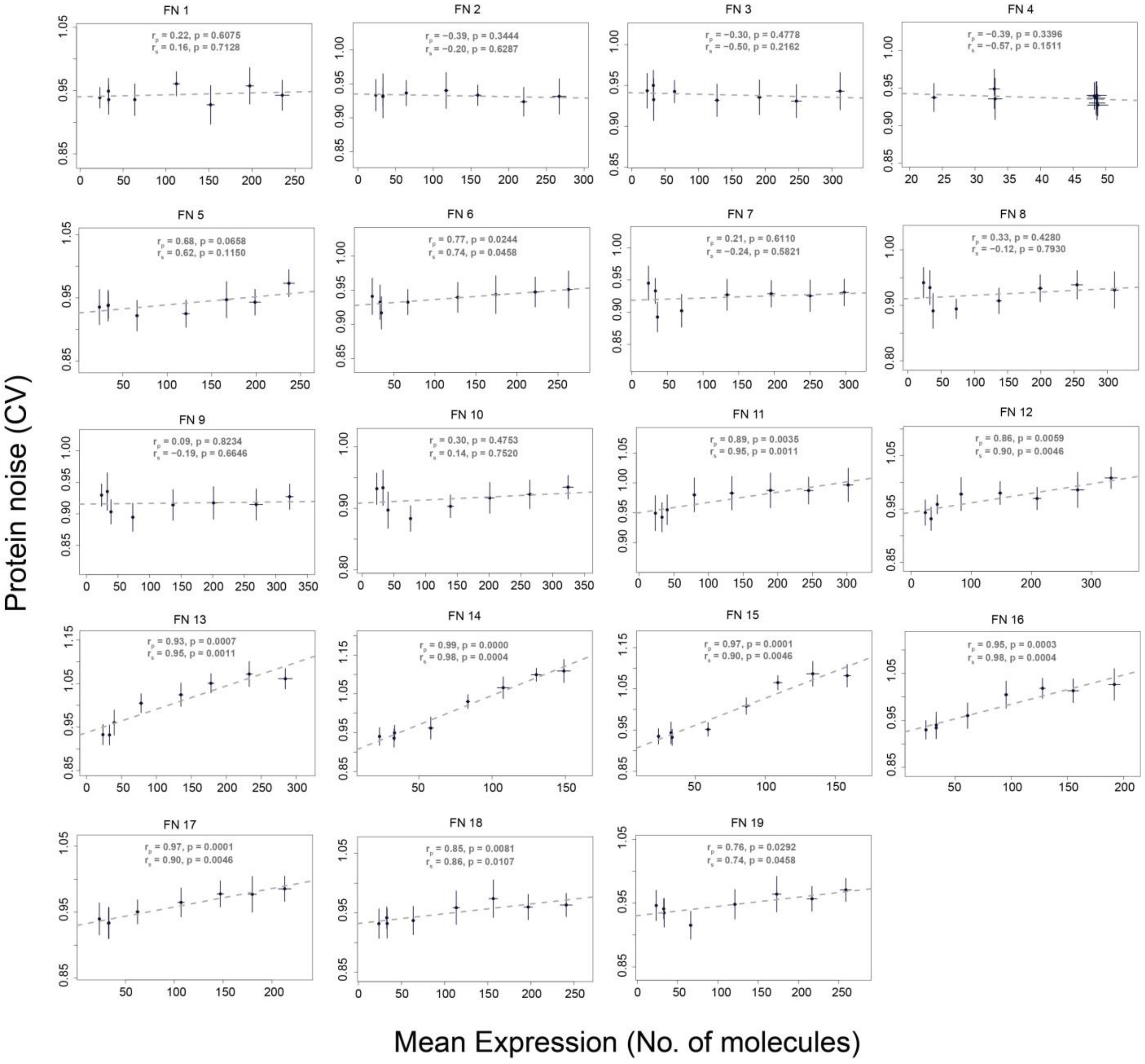


**Fig. S8.**
Relationship between mean protein expression and protein noise (CV) derived from stochastic simulations based on different mathematical functions to model ribosome demand (Table S1). Functions 11, 12 and 16 are repeated from Fig. 4 for sake of completeness.

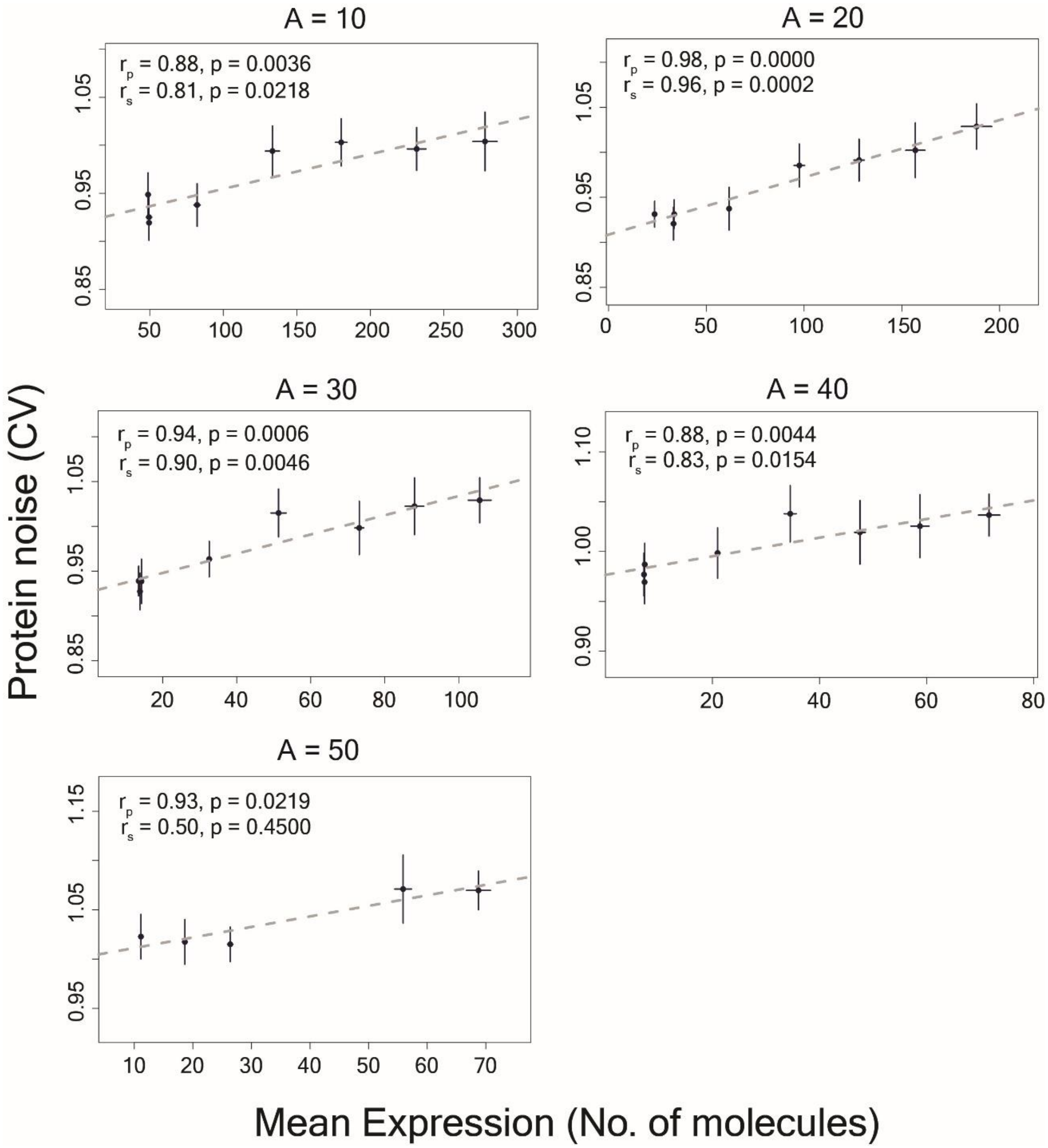


**Fig. S9.**

Relationship between mean protein expression and protein noise (CV) derived from stochastic simulations for different values of the parameter A.

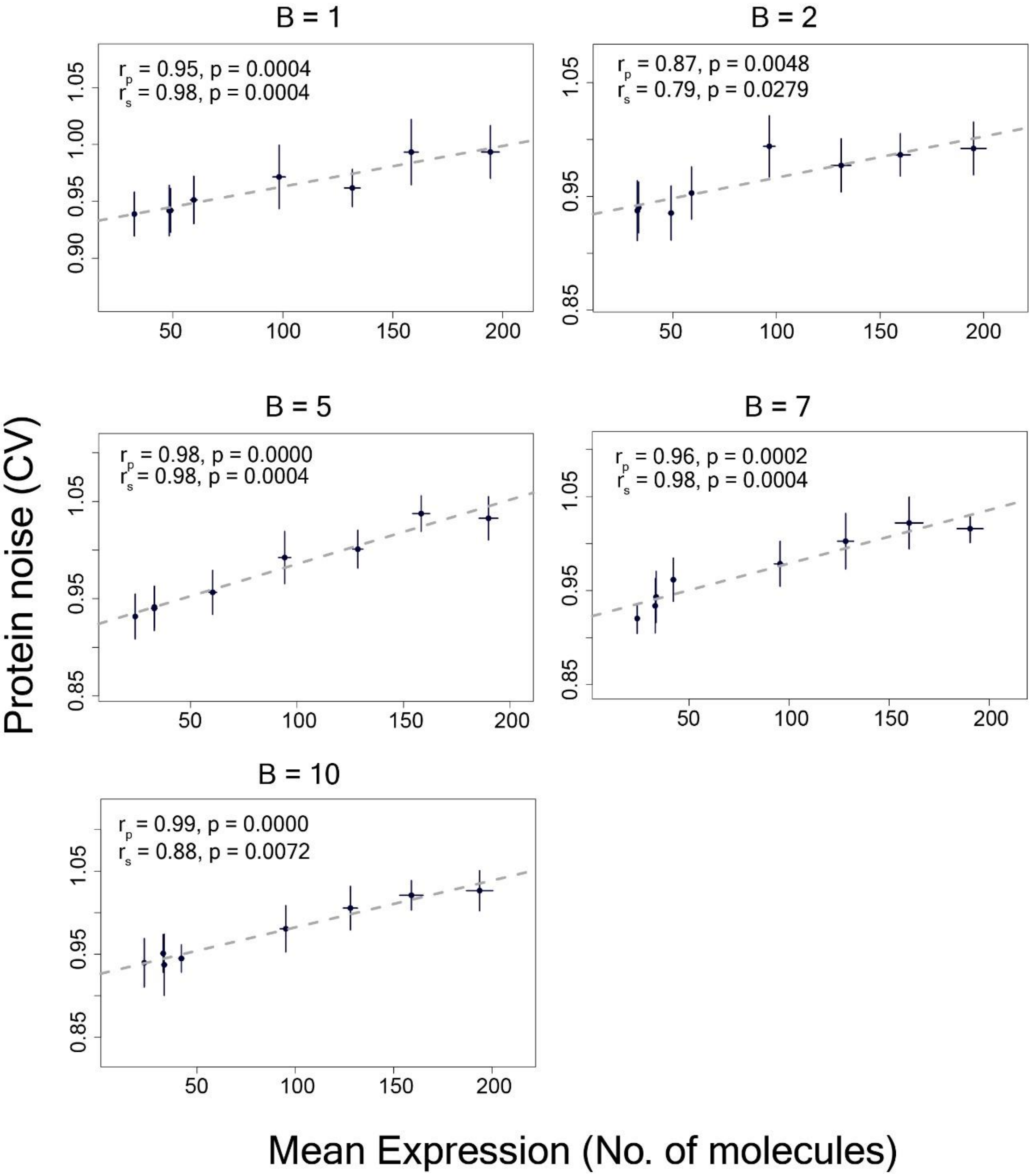


**Fig. S10.**

Relationship between mean protein expression and protein noise (CV) derived from stochastic simulations for different values of the parameter B.

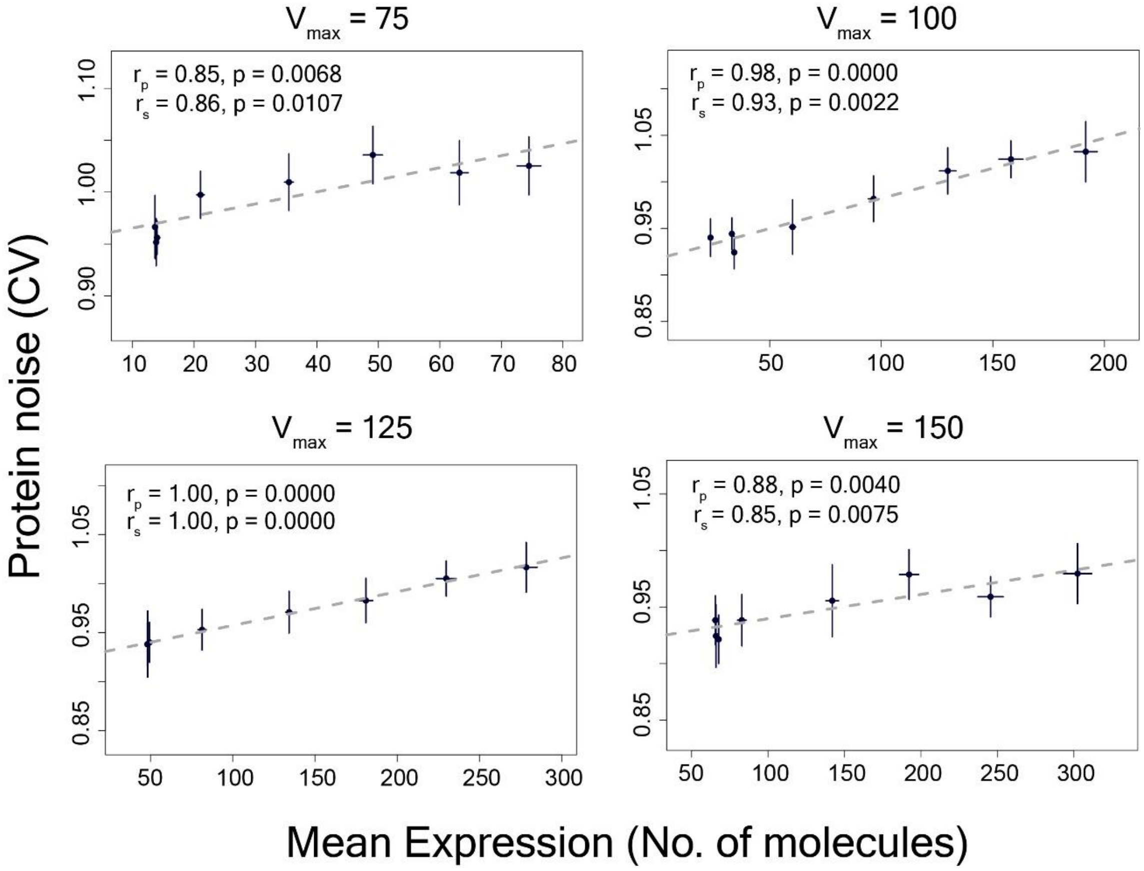


**Fig. S11.**

Relationship between mean protein expression and protein noise (CV) derived from stochastic simulations for different values of the parameter $V_{max}$

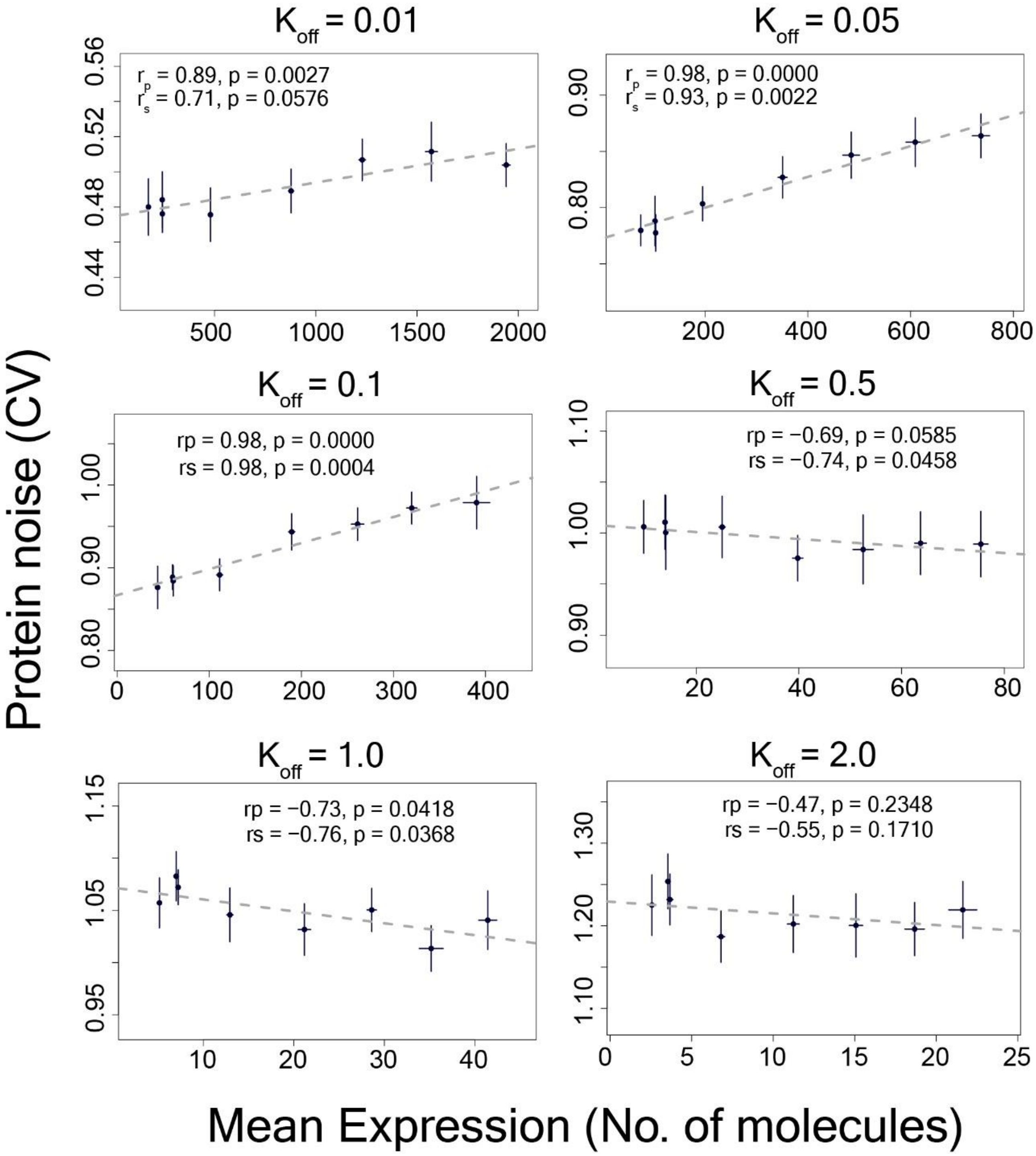


**Fig. S12.**

Relationship between mean protein expression and protein noise (CV) derived from stochastic simulations for different values of the parameter $K_{off}$

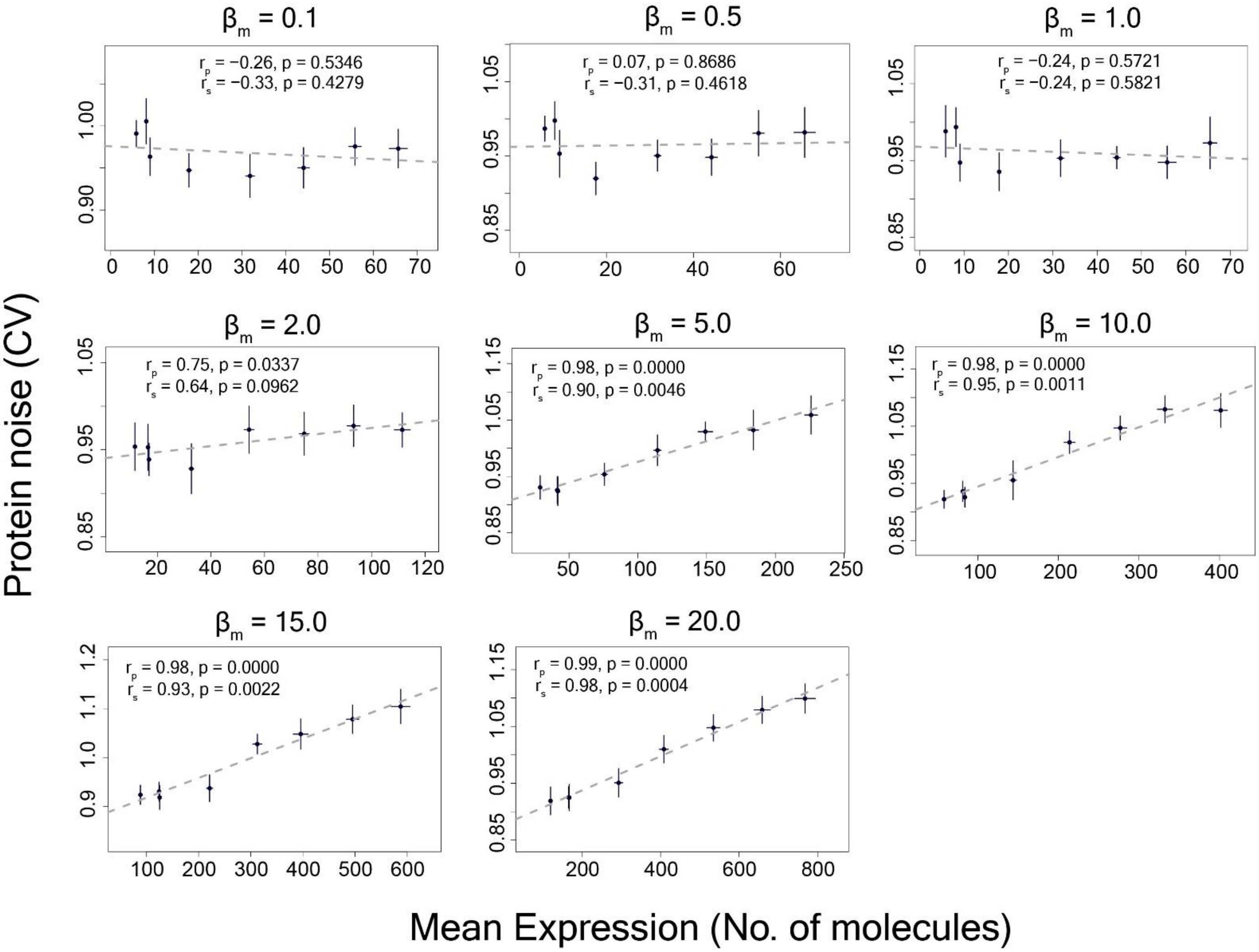


**Fig. S13.**

Relationship between mean protein expression and protein noise (CV) derived from stochastic simulations for different values of the parameter $\beta_m$

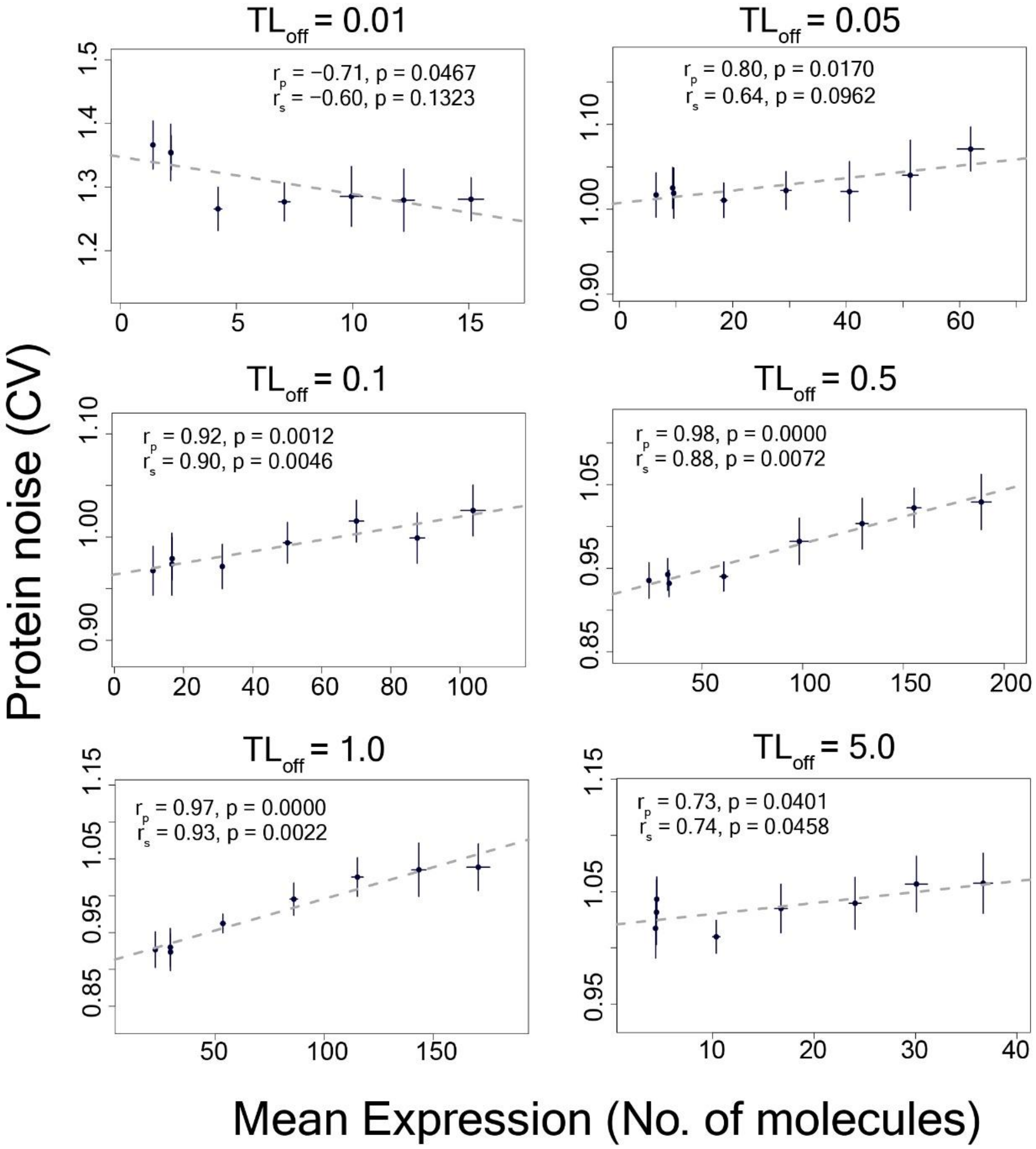


**Fig. S14.**

Relationship between mean protein expression and protein noise (CV) derived from stochastic simulations for different values of the parameter $TL_{off}$

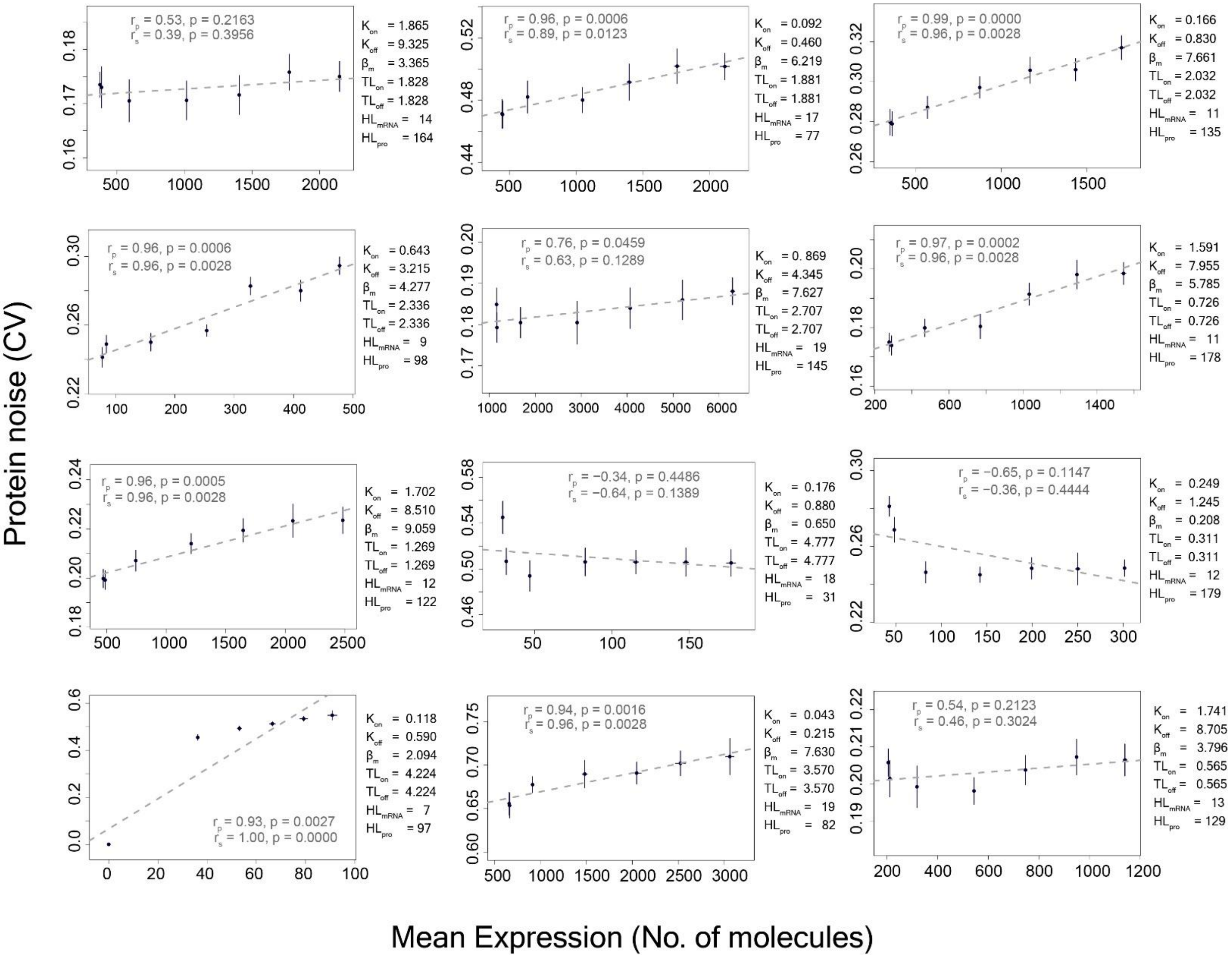


**Fig. S15.**

Relationship between mean protein expression and protein noise (CV) derived from stochastic simulations for sets of parameter values obtained by random sampling of parameter space

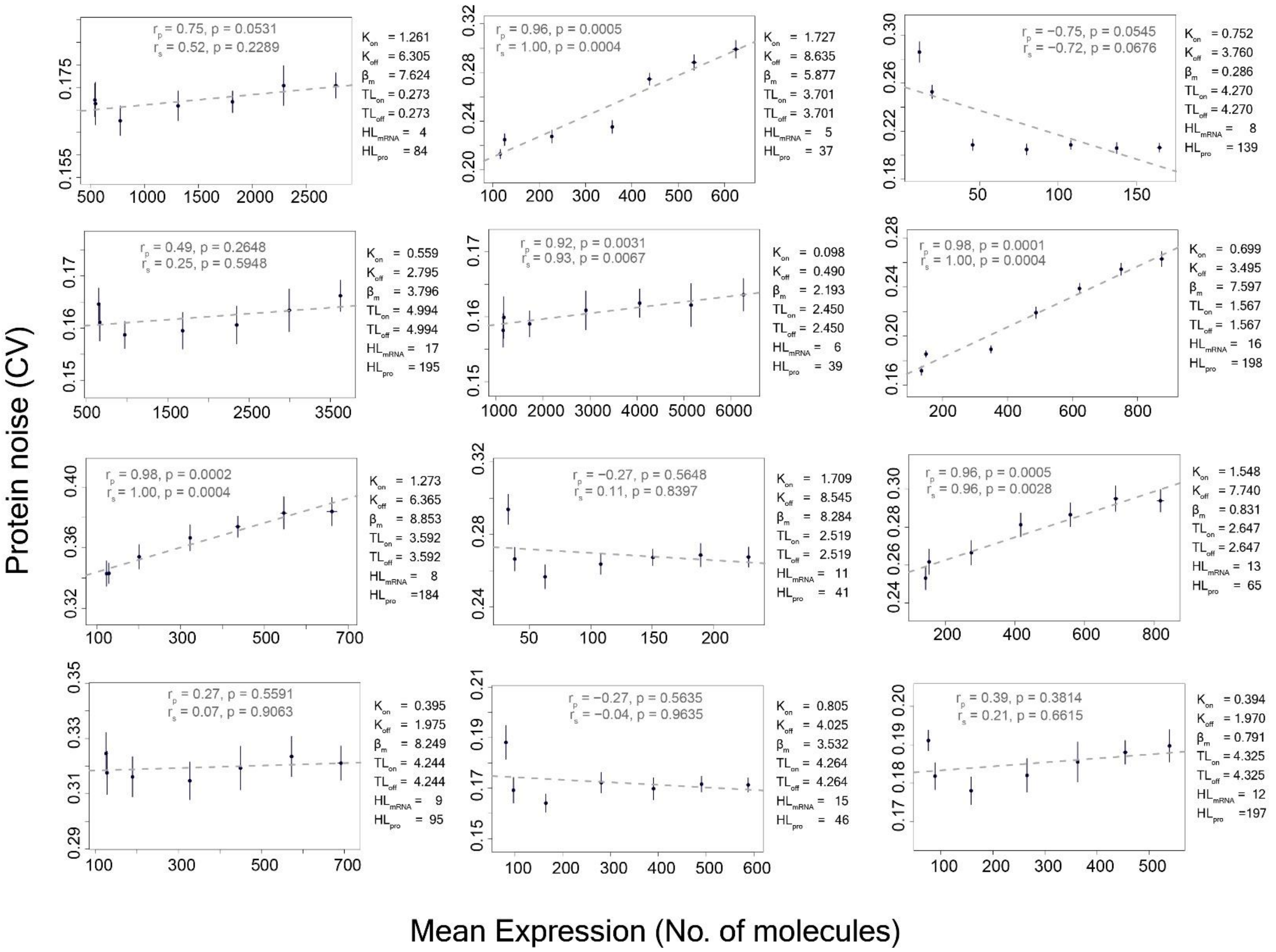


**Fig. S15 (continued)**

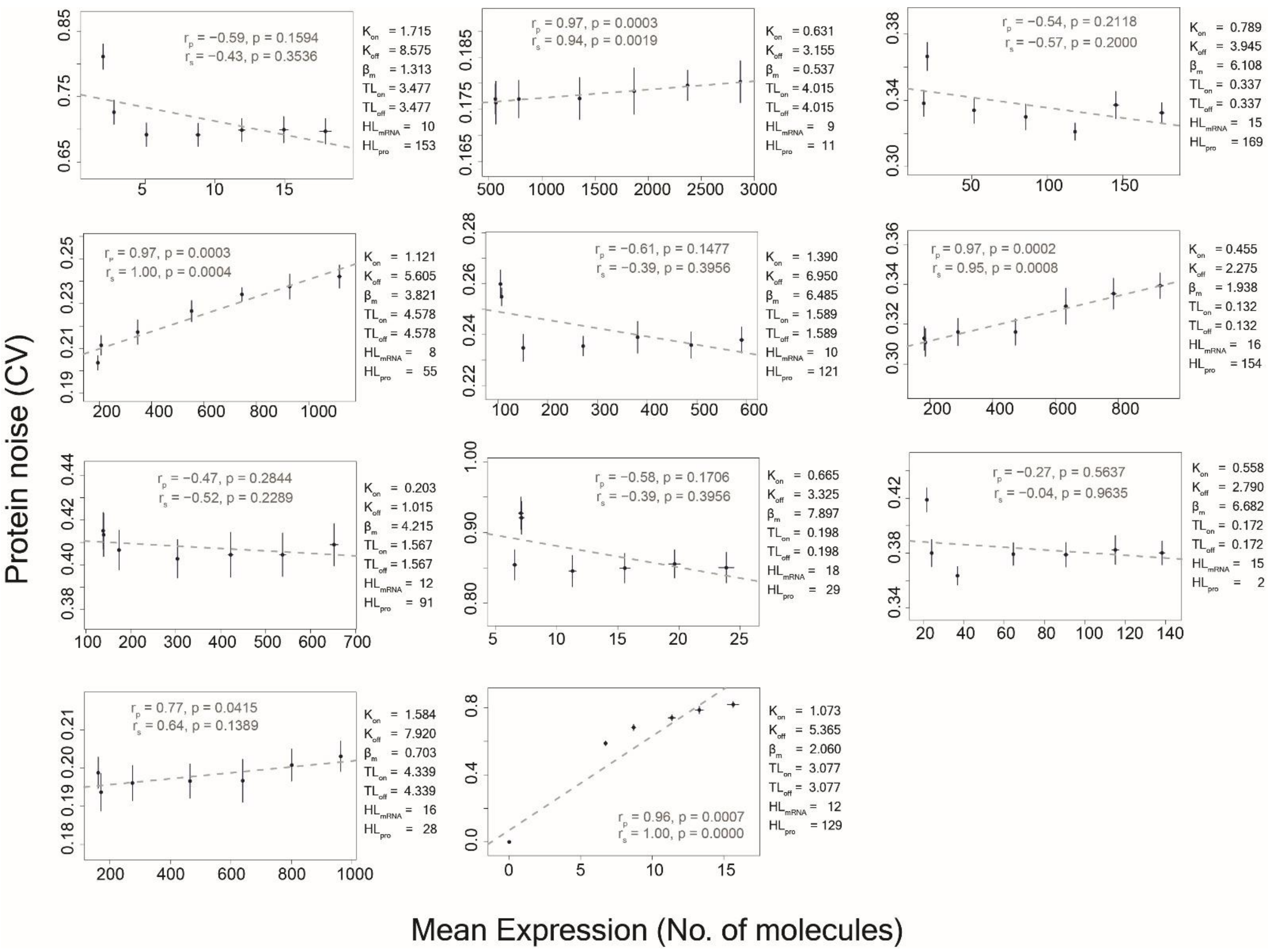


**Fig. S15 (continued)**

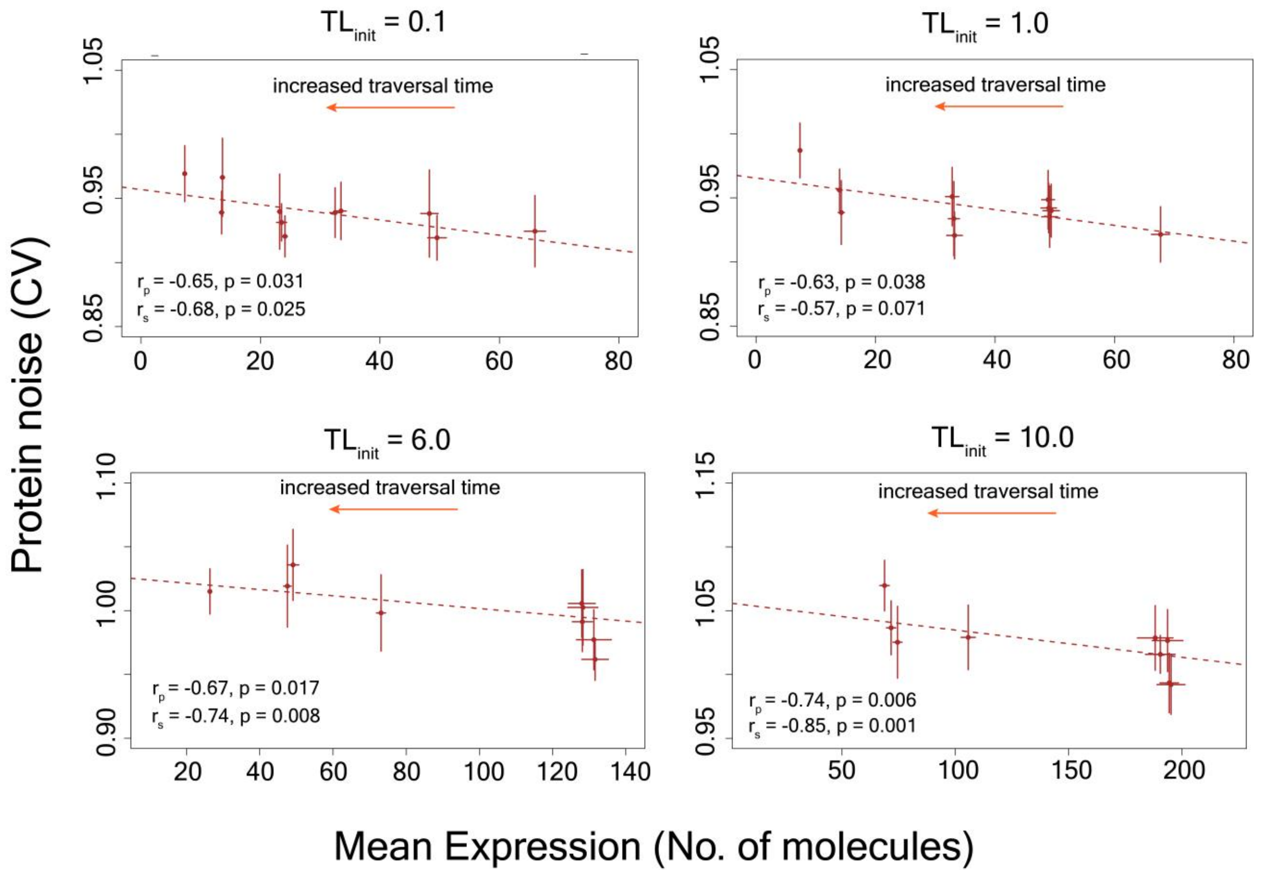


**Figure S16.**

Stochastic simulations where mean protein expression was altered by changing the ribosome traversal speed (Eq. 10) but keeping the base translation initiation rate ($TL_{init}$) constant, and thus, not allowing variation in ribosome demand with changes in ribosome traversal speed, abolished the positive correlation between mean protein expression and protein noise.

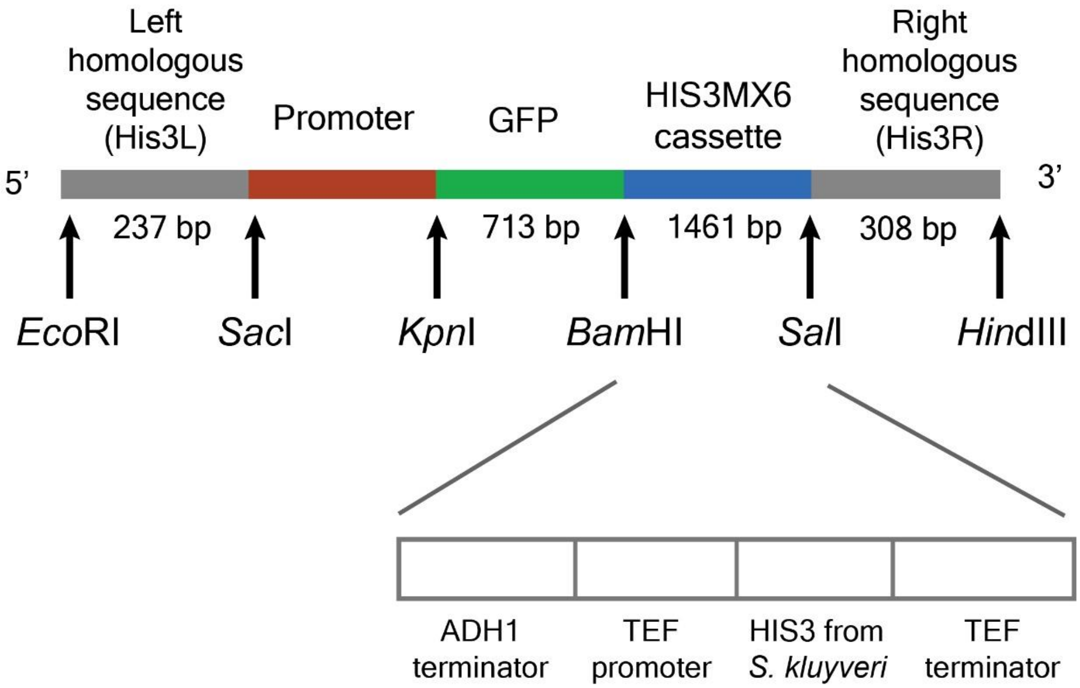


**Fig, S17.**

Structure of the promoter-GFP constructs along with auxotrophic marker (*HIS3MX6*) and homologous recombination sites (His3L and His3R) for genomic integration into yeast. The restriction sites for building the construct are shown by arrows.

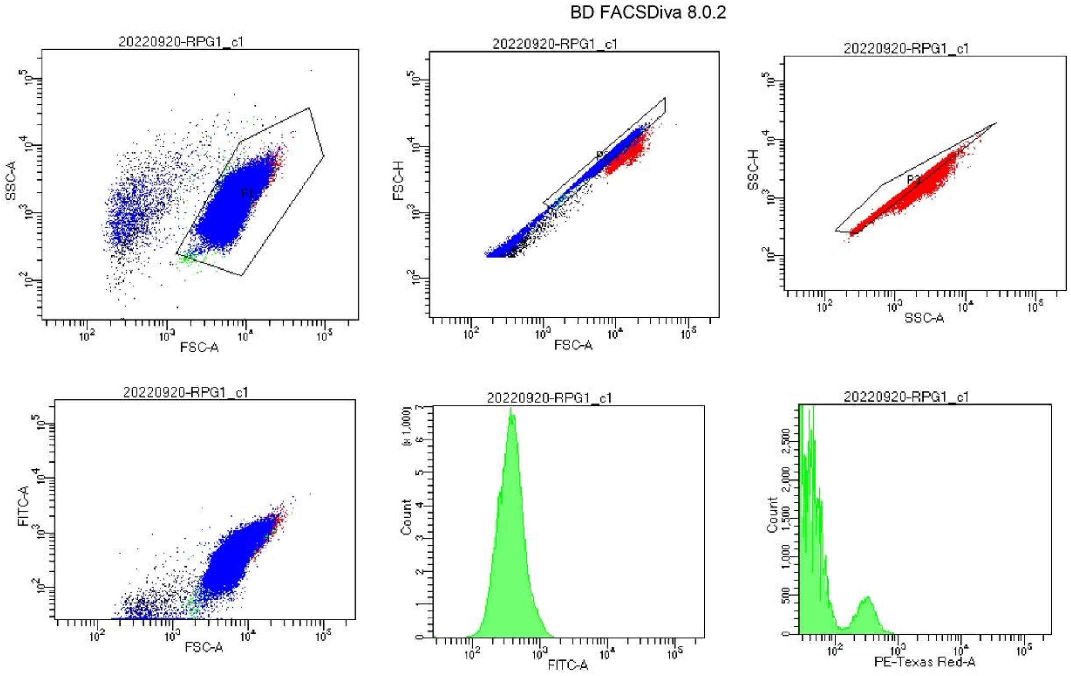


Tube Name: RPG1_c1
GUID: 7e290436-55ce-4f51-bf64-dcc3bc77e0d3

| Population | #Events | %Parent | FSC-A Mean | SSC-A Mean | FITC-A Mean | FITC-A Median | FITC-A SD | FITC-A %CV | PE-Texas Re... Mean | PE-Texas Re... Median |
|---|---|---|---|---|---|---|---|---|---|---|
| All Events | 200,000 | #### | 6,443 | 1,393 | 399 | 356 | 212 | 53.3 | 33 | 14 |
| P1 | 197,616 | 98.8 | 6,511 | 1,384 | 403 | 359 | 209 | 52.0 | 32 | 14 |
| P2 | 190,875 | 95.4 | 6,270 | 1,335 | 387 | 352 | 189 | 48.8 | 31 | 13 |
| P3 | 190,838 | 95.4 | 6,211 | 1,330 | 384 | 351 | 190 | 49.5 | 31 | 13 |

**Fig. S18.**
Snapshot of a flow cytometry experiment for measurement of mean protein expression and protein noise.

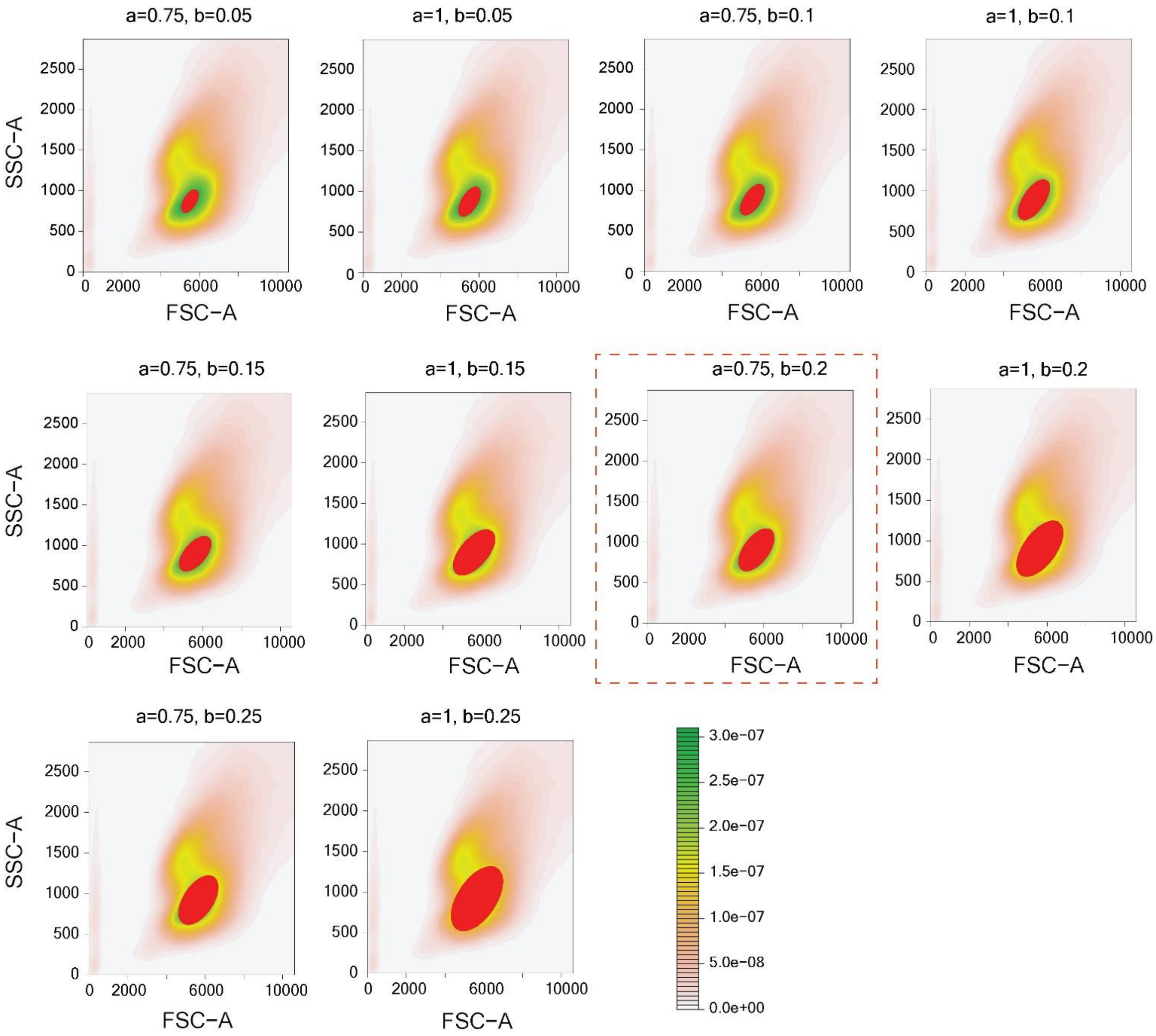


**Fig. S19.**

Fitting of ellipses (shown by red color) with different values of major and minor axes (‘a’ and ‘b’) and identification of the best fit that chooses a homogenous set of cells and contain at least ~20,000 cells for quantification of noise.

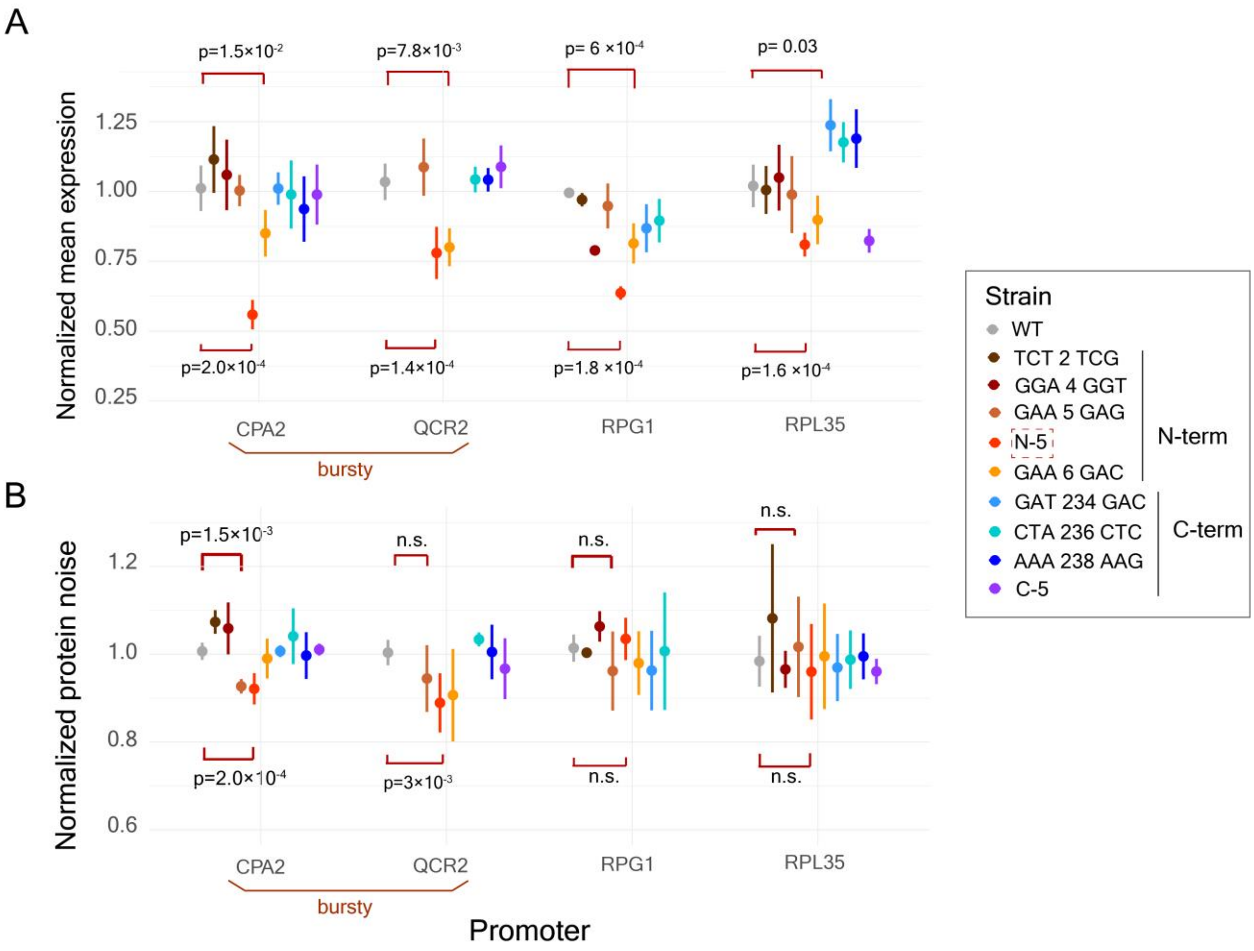


**Figure S20.**

**(A-B)** Normalized mean protein expression (A) and normalized protein noise (B) of GFP variants under the regulation of two bursty promoters *CPA2* and *QCR2*, and two non-bursty promoters *RPG1* and *RPL35A*. The p-values shows the results of Mann-Whitney U test to test for differences in normalized mean expression of normalized protein noise between a GFP variant and the wild-type.

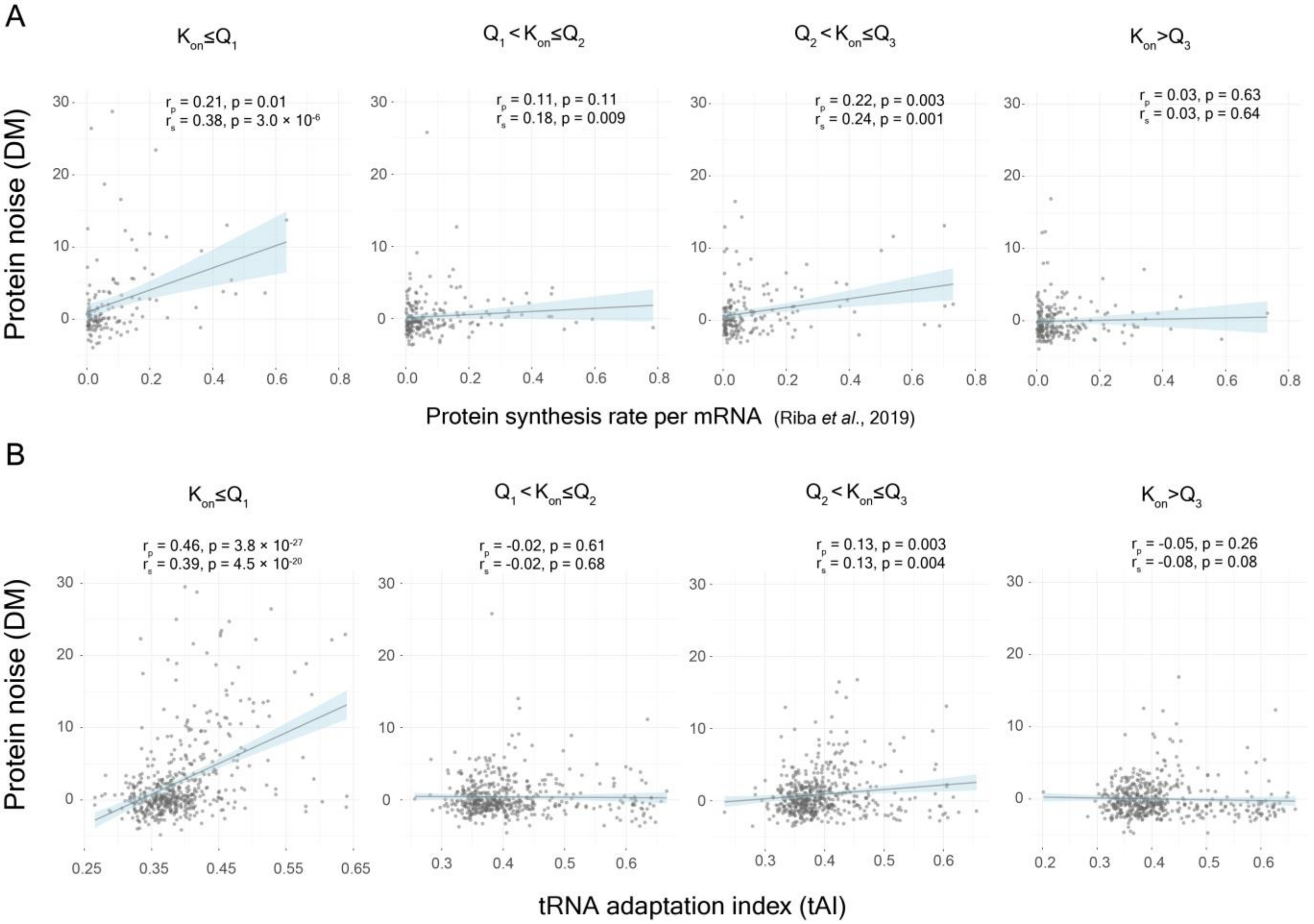


**Fig. S21.**

**(A)** Correlation between protein synthesis rate per mRNA[41] and protein noise, DM[15], for groups of genes with different values of transcriptional burst frequency ($K_{on}$). **(B)** Correlation between tRNA adaptation index (tAI) and protein noise, DM[15], for groups of genes with different values of transcriptional burst frequency ($K_{on}$). $Q_1$, $Q_2$, and $Q_3$ represent quartiles.

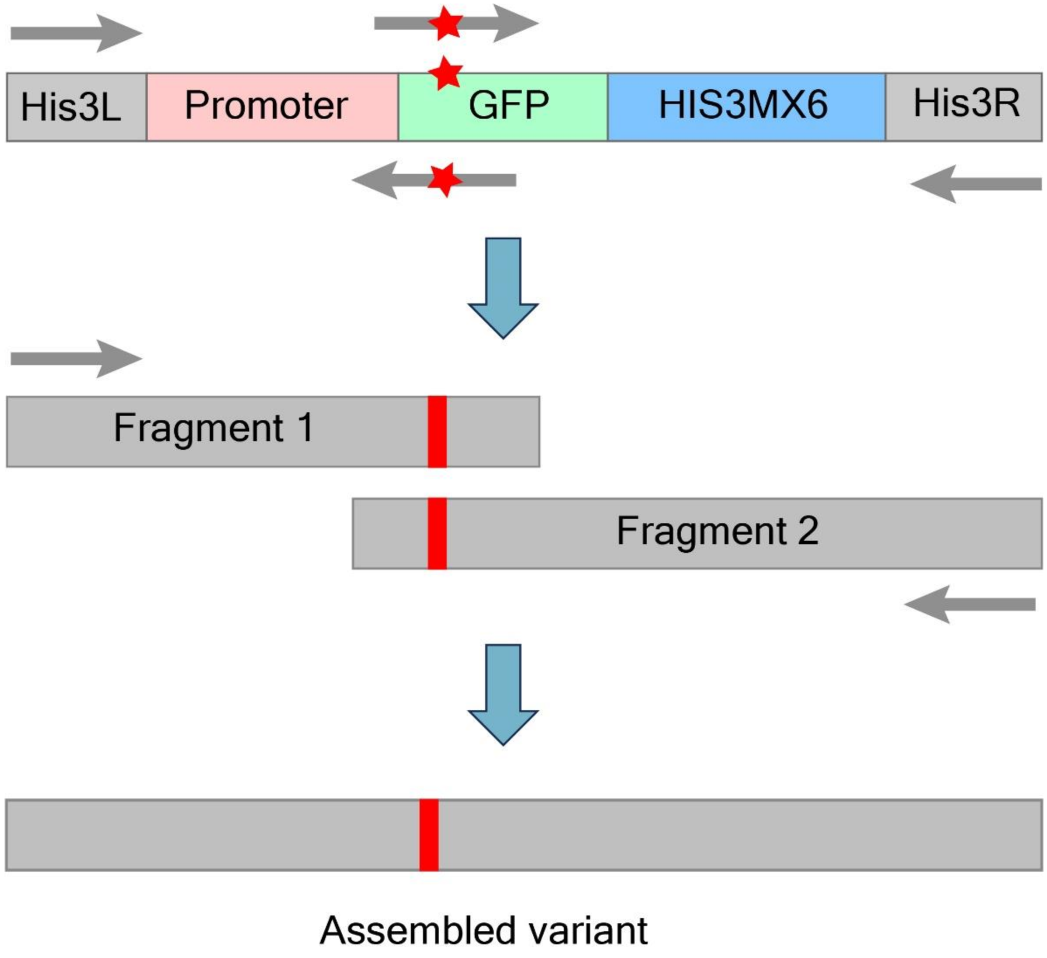


**Fig. S22.**

Construction of GFP mutant variants along with different promoters for genomic integration into yeast by overlap-extension PCR

**Table S1.**

List of mathematical functions explored to model ribosome demand

$mRNA_{prev}$ - Number of mRNA molecules at the previous time-point during simulation
$mRNA_{curr}$ - Number of mRNA molecules present at the present time-point during simulation
$ribo_{prev}$ - Number of ribosomes that were bound to mRNA molecules at the previous time-point during simulation
$ribo_{curr}$ - Number of ribosomes that are bound to mRNA molecules at the current time-point during simulation

$tlinitr_{base}$ – The basal translation initiation rate
$tlinitr_{curr}$ - the translation initiation rate at the current time-point during simulation

| Function number (FN) | Function |
|---|---|
| 1 | av = $\frac{1+ mRNA_{curr}}{1+ mRNA_{prev}}$ <br> kc = $1 + \frac{5}{(1+ mRNA_{prev})}$ <br> $tlinitr_{curr} = tlinitr_{base} \times \frac{kc^{10}}{kc^{10} + av^{10}}$ |
| 2 | av = $\frac{1+ mRNA_{curr}}{1+ mRNA_{prev}}$ <br> kc = $1 + \frac{6}{(1+ mRNA_{prev})}$ <br> $tlinitr_{curr} = tlinitr_{base} \times \frac{kc^{10}}{kc^{10} + av^{10}}$ |
| 3 | av = $\frac{1+ mRNA_{curr}}{1+ mRNA_{prev}}$ <br> kc = $1 + \frac{10}{(1+ mRNA_{prev})}$ <br> $tlinitr_{curr} = tlinitr_{base} \times \frac{kc^{10}}{kc^{10} + av^{10}}$ |
| 4 | av = $\frac{1+ mRNA_{curr}}{1+ mRNA_{prev}}$ <br> kc = $[1 + \frac{10}{(1+ mRNA_{prev})}] \times \frac{1}{(1+ \frac{ribo_{curr}}{50})^{10}}$ <br> $tlinitr_{curr} = tlinitr_{base} \times \frac{kc^{10}}{kc^{10} + av^{10}}$ |
| 5 | av = $\frac{1+ ribo_{curr}}{1+ ribo_{prev}}$ <br> kc = $1 + \frac{5}{(1+ ribo_{prev})}$ |

| | |
|---|---|
| | $tlinitr_{curr} = tlinitr_{base} \times \frac{kc^{10}}{kc^{10} + av^{10}}$ |
| 6 | av = $\frac{1+ ribo_{curr}}{1+ ribo_{prev}}$<br>kc = 1 + $\frac{10}{(1+ ribo_{prev})}$<br>$tlinitr_{curr} = tlinitr_{base} \times \frac{kc^{10}}{kc^{10} + av^{10}}$ |
| 7 | av = $\frac{1+ ribo_{curr}}{1+ ribo_{prev}}$<br>kc = 1 + $\frac{20}{(1+ ribo_{prev})}$<br>$tlinitr_{curr} = tlinitr_{base} \times \frac{kc^{10}}{kc^{10} + av^{10}}$ |
| 8 | av = $\frac{1+ ribo_{curr}}{1+ ribo_{prev}}$<br>kc = 1 + $\frac{30}{(1+ ribo_{prev})}$<br>$tlinitr_{curr} = tlinitr_{base} \times \frac{kc^{10}}{kc^{10} + av^{10}}$ |
| 9 | av = $\frac{1+ ribo_{curr}}{1+ ribo_{prev}}$<br>kc = 1 + $\frac{40}{(1+ ribo_{prev})}$<br>$tlinitr_{curr} = tlinitr_{base} \times \frac{kc^{10}}{kc^{10} + av^{10}}$ |
| 10 | av = $\frac{1+ ribo_{curr}}{1+ ribo_{prev}}$<br>kc = 1 + $\frac{50}{(1+ ribo_{prev})}$<br>$tlinitr_{curr} = tlinitr_{base} \times \frac{kc^{10}}{kc^{10} + av^{10}}$ |
| 11 | av = $\frac{1+ mRNA_{curr}}{1+ mRNA_{prev}}$<br>$tlinitr_{curr} = tlinitr_{base} \times \frac{1}{av}$ |
| 12 | av = $\frac{1+ ribo_{curr}}{1+ ribo_{prev}}$<br>$tlinitr_{curr} = tlinitr_{base} \times \frac{1}{av}$ |
| 13 | av = $\left(\frac{1+ mRNA_{curr}}{1+ mRNA_{prev}}\right) \times \left(\frac{1+ ribo_{curr}}{1+ ribo_{prev}}\right)$ |

| | |
|---|---|
| | $tlinitr_{curr} = tlinitr_{base} \times \frac{1}{av}$ |
| 14 | $av = \left(\frac{1+ mRNA_{curr}}{1+ mRNA_{prev}}\right) \times \left(\frac{1+ ribo_{curr}}{1+ ribo_{prev}}\right)$<br>$kc = 1 + \frac{1}{(1+ mRNA_{prev} + ribo_{prev})}$<br>$tlinitr_{curr} = tlinitr_{base} \times \frac{kc^{10}}{kc^{10} + av^{10}}$ |
| 15 | $av = \left(\frac{1+ mRNA_{curr}}{1+ mRNA_{prev}}\right) \times \left(\frac{1+ ribo_{curr}}{1+ ribo_{prev}}\right)$<br>$kc = 1 + \frac{2}{(1+ mRNA_{prev} + ribo_{prev})}$<br>$tlinitr_{curr} = tlinitr_{base} \times \frac{kc^{10}}{kc^{10} + av^{10}}$ |
| 16 | $av = \left(\frac{1+ mRNA_{curr}}{1+ mRNA_{prev}}\right) \times \left(\frac{1+ ribo_{curr}}{1+ ribo_{prev}}\right)$<br>$kc = 1 + \frac{5}{(1+ mRNA_{prev} + ribo_{prev})}$<br>$tlinitr_{curr} = tlinitr_{base} \times \frac{kc^{10}}{kc^{10} + av^{10}}$ |
| 17 | $av = \left(\frac{1+ mRNA_{curr}}{1+ mRNA_{prev}}\right) \times \left(\frac{1+ ribo_{curr}}{1+ ribo_{prev}}\right)$<br>$kc = 1 + \frac{10}{(1+ mRNA_{prev} + ribo_{prev})}$<br>$tlinitr_{curr} = tlinitr_{base} \times \frac{kc^{10}}{kc^{10} + av^{10}}$ |
| 18 | $av = \left(\frac{1+ mRNA_{curr}}{1+ mRNA_{prev}}\right) \times \left(\frac{1+ ribo_{curr}}{1+ ribo_{prev}}\right)$<br>$kc = 1 + \frac{15}{(1+ mRNA_{prev} + ribo_{prev})}$<br>$tlinitr_{curr} = tlinitr_{base} \times \frac{kc^{10}}{kc^{10} + av^{10}}$ |
| 19 | $av = \left(\frac{1+ mRNA_{curr}}{1+ mRNA_{prev}}\right) \times \left(\frac{1+ ribo_{curr}}{1+ ribo_{prev}}\right)$<br>$kc = 1 + \frac{25}{(1+ mRNA_{prev} + ribo_{prev})}$<br>$tlinitr_{curr} = tlinitr_{base} \times \frac{kc^{10}}{kc^{10} + av^{10}}$ |

**Table S2.**

List of primers used

| Primer name | Sequence (5′ to 3′) |
|---|---|
| GFP_for | AGCGGTACCATGTCTAAAGGAGAAGAACTTTTCACTGGAG |
| GFP_rev | AGCGGATCCCTATTTGTATAGTTCATCCATGCCATGTG |
| His3L_for | AGCGAATTCAACACAGTCCTTTCCCGCA |
| His3L_rev | AGCGAGCTCCTTTGCCTTCGTTTATCTTGCCTG |
| His3R_for | AGCGGATCCGTCGACTGACACCGATTATTTAAAG |
| His3R_rev | AGCAAGCTTAATATGAAATGCTTTTCTTGTTGTTCTTAC |
| HIS3MX_for | AGCGGATCCGGCGCGCCACTTCTAAATAAG |
| HIS3MX_rev | AGCAAGCTTGTCGACCAGTATAGCGACCAG |
| RPL35Ap_for | AGCGAGCTCAAATTCTAGAATATGGATCAAATACGCTTG |
| RPL35Ap_rev | AGCGGTACCTTCGAATATACTGTCTCACTGTACC |
| CPA2p_for | AGCGAGCTCCTTTGCGGAATGGTATACTATTTCTTTTC |
| CPA2p_rev | AGCGGTACCTCTTTTCTTCCTGTCTATGTACTGTATTG |
| QCR2p_for | AGCGAGCTCTTGGATCTTGCTCAACAAAAATTTTTC |
| QCR2p_rev | AGCGGTACCCAACGTTCTTCTTTTTTCCTTTTAATAATTTTAAC |
| RPG1p_for | AGCGAGCTCTTATTTCAAATTTTTCATGTCCTTGTATTTTTATTC |
| RPG1p_rev | AGCGGTACCCTTGATGTTTCCTGGGCGTTTG |
| NGFP_TCT2TCG_for | AGCGGTACCATGTCGAAAGGAGAAGAACTTTTCACTGGA |
| NGFP_GGA4GGT_for | AGCGGTACCATGTCTAAAGGTGAAGAACTTTTCACTGGA |
| RPG1-TCT2TCG_rev | GAAAAGTTCGTCTCCCTTAGACATGGTACCCTTGATGTTTCC |
| RPG1-GGA4GGT_rev | GAAAAGTTCCTCTCCTTTAGACATGGTACCCTTGATGTTTCC |
| RPL35A-TCT2TCG_rev | GAAAAGTTCGTCTCCCTTTGACATGGTACCTTCGAATATACTGTC |
| RPL35A-GGA4GGT_rev | GAAAAGTTCCTCTCCTTTTGACATGGTACCTTCGAATATACTGTC |
| CPA2-TCT2TCG_rev | GAAAAGTTCGTCTCCCTTAGACATGGTACCTCTTTTCTTCCTGTC |
| CPA2-GGA4GGT_rev | GAAAAGTTCCTCTCCTTTAGACATGGTACCTCTTTTCTTCCTGTC |
| QCR2-TCT2TCG_rev | GAAAAGTTCGTCTCCCTTAGACATGGTACCCAACGTTCTTC |
| QCR2-GGA4GGT_rev | GAAAAGTTCCTCTCCTTTAGACATGGTACCCAACGTTCTTC |
| NGFP_GAA5GAG_for | AGCGGTACCATGTCTAAAGGAGAGGAACTTTTCACTGGA |
| NGFP_GAA6GAC_for | AGCGGTACCATGTCTAAAGGAGACGAACTTTTCACTGGA |
| NGFP_GAT234GAC_rev | AGCGGATCCCTATTTGTATAGTTCGTCCATGCCATGTGT |
| NGFP_CTA236CTC_rev | AGCGGATCCCTATTTGTAGAGTTCATCCATGCCATGTGT |
| NGFP_AAA238AAG_rev | AGCGGATCCCTACTTGTATAGTTCATCCATGCCATGTGT |
| N-5_for | AGCGGTACCATGTCGAAGGGTGAGGACCTTTTCACTGGA |
| C-5_rev | AGCGGTACCATGTCGAAGGGTGAGGACCTTTTCACTGGA |
| RPG1-GAA5GAG_rev | GAAAAGTTCCTCTCCTTTAGACATGGTACCCTTGATGTTTCC |
| RPG1- GAA6GAC_rev | GAAAAGGTCGTCTCCTTTAGACATGGTACCCTTGATGTTTCC |
| RPG1-N-5_rev | GAAAAGGTCCTCACCCTTCGACATGGTACCCTTGATGTTTCC |
| QCR2- GAA5GAG _rev | GAAAAGTTCCTCTCCTTTAGACATGGTACCCAACGTTCTTC |
| QCR2- GAA6GAC_rev | GAAAAGGTCGTCTCCTTTAGACATGGTACCCAACGTTCTTC |
| QCR2-N-5_rev | GAAAAGGTCCTCACCCTTCGACATGGTACCCAACGTTCTTC |
| CPA2- GAA5GAG _rev | GAAAAGTTCCTCTCCTTTAGACATGGTACCTCTTTTCTTCCTGTC |
| CPA2- GAA6GAC_rev | GAAAAGGTCGTCTCCTTTAGACATGGTACCTCTTTTCTTCCTGTC |
| CPA2-N-5_rev | GAAAAGGTCCTCACCCTTCGACATGGTACCTCTTTTCTTCCTGTC |
| GAT234GAC-HMX_for | GGCATGGACGAACTATACAAATAGGGATCCGGCGCGCCACTTC |

| | |
|---|---|
| CTA236CTC-HMX_for | GGCATGGATGAACTCTACAAATAGGGATCCGGCGCGCCACTTC |
| AAA238AAG-HMX_for | GGCATGGATGAACTATACAAGTAGGGATCCGGCGCGCCACTTC |
| C-5-HMX_for | GGCATGGACGACCTCAACAAGTAGGGATCCGGCGCGCCACTTC |

**Table S3**
Fragment size and nucleotide sequence of the parts of the promoter-GFP constructs

| **Fragment** | **Sequence** | **Size (bps)** |
|---|---|---|
| His3L | AACACAGTCCTTTCCCGCAATTTTCTTTTTCTATTACTCTTGGCCTCCTCTA<br>GTACACTCTATATTTTTTATGCCTCGGTAATGATTTTCATTTTTTTTTTTTC<br>CACCTAGCGGATGACTCTTTTTTTTTCTTAGCGATTGGCATTATCACATAAT<br>GAATTATACATTATATAAAGTAATGTGATTTCTTCGAAGAATATACTAAAAA<br>ATGAGCAGGCAAGATAAACGAAGGCAAAG | 237 |
| RPL35A promoter | AAATTCTAGAATATGGATCAAATACGCTTGTATAAACTAAATGAAACA<br>TAAAGATTAAGAACTTAAGAGGCCAACGTCGATGGATTTATTGACGAT<br>CACCAGCCAACACATATAGATTTTAGTGTAAAAGCAATAAAAACCAAG<br>ATAATAAAATAAAAAAATACTGAAGAAGCCTAACTAGTATAAACTACT<br>TTAACTAATAATGGCAATTTGATATAGAAACAAAGAAACATGATATAT<br>TTAGGATATTATACAACGCATTTTCATTTGTTTTACAGCACCCCTGCG<br>TGAATCATATATTGACGTTTCGCTCTCAGGTCCACCGTGTTCTCAAAA<br>GATACTTTTAAAACCTAAAACACACGAAATCATATTATGATAATTCAG<br>AATGATAGTGTGGTACTGTGTCAATTGACTGTTCAAGACTGAAGAGGA<br>TCTTTGATTTGTTGTTACTCAACAAATAATCTTCACGAAAACTTTCTC<br>AATCTGGGGACTGTATTAATCTCAGACCCATACATATCTACACCCATA<br>ACTTTTTACATTTAATTTTTTATCACATAATAGGTAGCTTAAATTGTA<br>AAGTCGCAAAAAAAAATGGCAGCGCAGCCTCTCCGGGTGAACCCCACG<br>ACAACTTACCTGGCACTCCATGCACTAACGGGCGGGTTTGGGCAGGAT<br>TCCAGCATCAATTTTGCAAAATTCACACCTGAGTAATTCATATATGTA<br>ATATAATGTTAAGCATACGCTGTCGATTAGCACTATTATTGACCGTAG<br>AATAGGTACAGTGAGACAGTATATTCGAA | 797 |
| RPG1 promoter | TTATTTCAAATTTTTCATGTCCTTGTATTTTTATTCTTTATCCCTTCCAATC<br>AGAAAGGATCTAGTGAACAAGTTCTTTCCTCTATGGTATATATTTTAGTGAT<br>AAATTTTATAAAATTATCAAAACCAAGGCATCCTTTCCTTTTATTCTGTCAT<br>TGGAATCTGCCTGTCATAATTATCACCCACCGGGTAAAGATGATAATTTTTC<br>AGTCGCTTTGCCCGAGAAGCTTTGCCAGGTGAAAAATTTTTCTTGGTGAACT<br>AAATCGAAAGTAGATATACTTACAACTATAGAGAGAGTTCCAAAATAAACCA<br>TACAAACGCCCAGGAAACATCAAG | 336 |
| CPA2 promoter | CTTTGCGGAATGGTATACTATTTCTTTTCCCTCTTTTTAGCACTATTACACC<br>CCGCCCACAAAATAAAAATAATAACGACGACCTAATCTCACCAAGTGACCCT<br>TGTAAAACCTCCTTTTCTTTATAATGTTTCTTTTCTTACTAATATTTGGTAC | 673 |

| | | |
|---|---|---|
| | ATTTAGGGTAGTGATAAAAGAATGGCAACATTGTTATTATTGTGAAAAATGA<br>GGAAAATAGAAAATCAGAAACCCTAAAAAGTGATTTTACCCTATCAGAAATA<br>TTCAAATGTCCTAATTAAAAATAGTAAATCCCCTAAACATTCAGATTGTAAA<br>CTAGGGTTGAGAAAATGACTCATCCACCACTGTCTTCTTTCCTGCGGCATTC<br>TATAGATTATTGTGAATGACTCTTATTGATGAGATGGCAATAACTTTTGAAT<br>ATCAGAGATAGGAACCTCCATGTCGTAACGATTGTGTCACCTTGAGTAAGCA<br>TCGAGAAAATCCAATCTTTTTTTTTCCGTCATAAGCATTTCTGCCATGCTAT<br>TTGTATATATATAATTACTAATACGTCTTCTATAGTATGCCTTATCTCTTTT<br>TTGAAGCGCTATTTAAGTTTAAGCATCGAAAAACTAACATCTATAGTTAAAA<br>TTAGTTCTATAAAGGAAGAGCAATACAGTACATAGACAGGAAGAAAAGA | |
| QCR2 promoter | TTGGATCTTGCTCAACAAAAATTTTTCCCTATTCCTGTGTCAGCAATTCTCT<br>ATTGTTCTCAATATTTCTTCCACTATTATTATTTTGATCCAAAATTATTTTT<br>TTCCTTCAATGCGATGAGCTTTTGAAAAATTTCTGATCATTCCCAACGAACC<br>AATAGAAGGCCCGCCCCGTCTTATATCCGTTAGCCTACCAAATATATATATA<br>AAGAACAAGGGCCTTTCCTCAGAGCGTTTGCTGACGAAGTTTTAGAAGTTAA<br>TAAGGTTTTTAACAGCAGTGTGCTCGAACGATTAGGACGGGAGAGTTAAAAT<br>TATTAAAAGGAAAAAAGAAGAACGTTG | 339 |
| GFP | ATGTCTAAAGGAGAAGAACTTTTCACTGGAGTTGTCCCAATTCTTGTTGAAT<br>TAGATGGTGATGTTAATGGGCACAAATTTTCTGTCAGTGGAGAGGGTGAAGG<br>TGATGCAACATACGGAAAACTTACCCTTAAATTTATTTGCACTACTGGAAAA<br>CTACCTGTTCCATGGCCAACACTTGTCACTACTTTTACGTATGGTGTTCAAT<br>GCTTTTCAAGATACCCAGATCATATGAAACGGCATGACTTTTTCAAGAGTGC<br>CATGCCCGAAGGTTATGTACAGGAAAGAACTATATTTTTCAAAGATGACGGG<br>AACTACAAGACACGTGCTGAAGTCAAGTTTGAAGGTGATACCCTTGTTAATA<br>GAATCGAGTTAAAAGGTATTGATTTTAAAGAAGATGGAAACATTCTTGGACA<br>CAAATTGGAATACAACTATAACTCACACAATGTATACATCATGGCAGACAAA<br>CAAAAGAATGGAATCAAAGTTAACTTCAAAATTAGACACAACATTGAAGATG<br>GAAGCGTTCAACTAGCAGACCATTATCAACAAAATACTCCAATTGGCGATGG<br>CCCTGTCCTTTTACCAGACAACCATTACCTGTCCACACAATCTGCCCTTTCG<br>AAAGATCCCAACGAAAAGAGAGACCACATGGTCCTTCTTGAGTTTGTAACAG<br>CTGCTGGGATTACACATGGCATGGATGAACTATACAAATAG | 711 |
| HIS3MX6 | GGCGCGCCACTTCTAAATAAGCGAATTTCTTATGATTTATGATTTTTATTAT<br>TAAATAAGTTATAAAAAAAATAAGTGTATACAAATTTTAAAGTGACTCTTAG<br>GTTTTAAAACGAAAATTCTTATTCTTGAGTAACTCTTTCCTGTAGGTCAGGT<br>TGCTTTCTCAGGTATAGCATGAGGTCGCTCTTATTGACCACACCTCTACCGG<br>CAGATCTGTTTAGCTTGCCTCGTCCCCGCCGGGTCACCCGGCCAGCGACATG<br>GAGGCCCAGAATACCCTCCTTGACAGTCTTGACGTGCGCAGCTCAGGGGCAT<br>GATGTGACTGTCGCCCGTACATTTAGCCCATACATCCCCATGTATAATCATT<br>TGCATCCATACATTTTGATGGCCGCACGGCGCGAAGCAAAAATTACGGCTCC<br>TCGCTGCAGACCTGCGAGCAGGGAAACGCTCCCCTCACAGACGCGTTGAATT<br>GTCCCCACGCCGCGCCCCTGTAGAGAAATATAAAAGGTTAGGATTTGCCACT<br>GAGGTTCTTCTTTCATATACTTCCTTTTAAAATCTTGCTAGGATACAGTTCT | 1455 |

| | | |
|---|---|---|
| | CACATCACATCCGAACATAAACAACCATGGGTAGGAGGGCTTTTGTAGAAAG<br>AAATACGAACGAAACGAAAATCAGCGTTGCCATCGCTTTGGACAAAGCTCCC<br>TTACCTGAAGAGTCGAATTTTATTGATGAACTTATAACTTCCAAGCATGCAA<br>ACCAAAAGGGAGAACAAGTAATCCAAGTAGACACGGGAATTGGATTCTTGGA<br>TCACATGTATCATGCACTGGCTAAACATGCAGGCTGGAGCTTACGACTTTAC<br>TCAAGAGGTGATTTAATCATCGATGATCATCACACTGCAGAAGATACTGCTA<br>TTGCACTTGGTATTGCATTCAAGCAGGCTATGGGTAACTTTGCCGGCGTTAA<br>AAGATTTGGACATGCTTATTGTCCACTTGACGAAGCTCTTTCTAGAAGCGTA<br>GTTGACTTGTCGGGACGGCCCTATGCTGTTATCGATTTGGGATTAAAGCGTG<br>AAAAGGTTGGGGAATTGTCCTGTGAAATGATCCCTCACTTACTATATTCCTT<br>TTCGGTAGCAGCTGGAATTACTTTGCATGTTACCTGCTTATATGGTAGTAAT<br>GACCATCATCGTGCTGAAAGCGCTTTTAAATCTCTGGCTGTTGCCATGCGCG<br>CGGCTACTAGTCTTACTGGAAGTTCTGAAGTCCCAAGCACGAAGGGAGTGTT<br>GTAAAGGATACTGACAATAAAAAGATTCTTGTTTTCAAGAACTTGTCATTTG<br>TATAGTTTTTTTATATTGTAGTTGTTCTATTTTAATCAAATGTTAGCGTGAT<br>TTATATTTTTTTTCGCCTCGACATCATCTGCCCAGATGCGAAGTTAAGTGCG<br>CAGAAAGTAATATCATGCGTCAATCGTATGTGAATGCTGGTCGCTATACTG | |
| His3R | TGACACCGATTATTTAAAGCTGCAGCATACGATATATATACATGTGTATATA<br>TGTATACCTATGAATGTCAGTAAGTATGTATACGAACAGTATGATACTGAAG<br>ATGACAAGGTAATGCATCATTCTATACGTGTCATTCTGAACGAGGCGCGCTT<br>TCCTTTTTTCTTTTTGCTTTTTCTTTTTTTTTCTCTTGAACTCGAGAAAAAA<br>AATATAAAAGAGATGGAGGAACGGGAAAAAGTTAGTTGTGGTGATAGGTGGC<br>AAGTGGTATTCCGTAAGAACAACAAGAAAAGCATTTCATATT | 302 |
